# Sex-specific plasticity and the nutritional geometry of insulin-signaling gene expression in *Drosophila melanogaster*

**DOI:** 10.1101/2020.11.16.385708

**Authors:** Jeanne M.C. McDonald, Pegah Nabili, Lily Thorsen, Sohee Jeon, Alexander Shingleton

## Abstract

1.

**Background:** Sexual-size dimorphism (SSD) is replete among animals, but while the selective pressures that drive the evolution of SSD have been well studied, the developmental mechanisms upon which these pressures act are poorly understood. Ours and others’ research has shown that SSD in *Drosophila* reflects elevated levels of nutritional plasticity in females versus males, such that SSD increases with dietary intake and body size, a phenomenon called sex-specific plasticity (SSP). Additional data indicate that while body size in both sexes responds to variation in protein level, only female body size is sensitive to variation in carbohydrate level. Here we explore whether these difference in sensitivity at the morphological level are reflected by differences in how the insulin/IGF-signaling (IIS) and TOR-signaling pathways respond to changes in carbohydrates and proteins in females versus males, using a nutritional geometry approach.

**Results:** The IIS-regulated transcripts of *4E-BP* and *InR* most strongly correlated with body size in females and males respectively, but neither responded to carbohydrate level and so could not explain the sex-specific response to body size to dietary carbohydrate. Transcripts regulated by TOR-signaling did, however, respond to dietary carbohydrate in a sex-specific manner. In females, expression of *dILP5* positively correlated with body size, while expression of *dILP2,3* and *8,* was elevated on diets with a low concentration of both carbohydrate and protein. In contrast, we detected lower levels of dILP2 and 5 protein in the brains of females fed on low concentration diets. We could not detect any effect of diet on *dILP* expression in males.

**Conclusion:** Although females and males show sex-specific transcriptional responses to changes in protein and carbohydrate, the patterns of expression do not support a simple model of the regulation of body-size SSP by either insulin-or TOR-signaling. The data also indicate a complex relationship between carbohydrate and protein level, *dILP* expression and dILP peptide levels in the brain. In general, diet quality and sex both affect the transcriptional response to changes in diet quantity, and so should be considered in future studies that explore the effect of nutrition on body size.

## 2. Background

Sexual Size Dimorphism (SSD), the difference in body size between males and females, is perhaps the most familiar and widespread form of sexual dimorphism. This condition is extremely variable among species. For example, a female blanket octopi can weigh 10,000-20,000 times more than male [1], while a male southern elephant seals can weigh seven times more than a female [2]. Further, the degree of SSD is highly evolutionarily labile, and can vary between closely related species or among populations within species, sometimes dramatically. For example, among populations of the Australian carpet python, females range from being less than 1.5x to more than 10x the size of males [3]. While intraspecific variation in SSD is likely due, in part, to genetic differences between populations, it may also be a consequence of sex-specific differences in the phenotypic plasticity of body size. Environmental variation can account for the vast majority of variation in body size within a population [4, 5], and if males and females differ in the extent to which the environment affects body size, this will generate changes in SSD across environments [6]. While there is considerable evidence that males and females differ in the extent of their body size response to a variety of environmental variables – a phenomenon called *Sex-Specific Plasticity* (SSP)– the role that SSP plays in the developmental generation and evolution of sexual size dimorphism is largely unknown. This is compounded by a general lack of knowledge regarding the proximate developmental mechanisms that generate differences in body size between males and females.

Perhaps the most important, and certainly the best understood, environmental factor that regulates body size is developmental nutrition. In all animals where it has been studied, the nutritional regulation of growth is mediated via the insulin/IGF-signaling (IIS) pathway, a highly conserved receptor tyrosine kinase pathway, the components of which pre-date the Metazoa [7]. Insulin-like peptides are released in response to nutrition and bind to receptors of dividing cells to initiate a signal-transduction cascade that regulates the expression of positive and negative growth regulator. The IIS pathway positively regulates the activity of TOR-signaling, which also responds directly to cellular levels of amino acid. Activation of IIS/TOR-signaling results in an increase in both growth rate and (typically) adult body size at high levels of nutrition, while deactivation of IIS/TOR-signaling does the opposite at low levels of nutrition.

Although both males and female typically display nutritional plasticity, in many species one sex is more nutritionally plastic than the other. In birds and mammals, male growth tends to be affected more by nutritional stress than female growth [8], while in arthropods, the reverse appears to be true [9]. This is accompanied by the general trend of male-biased SSD in birds and mammals versus female-biased SSD in arthropods [10, 11]. Generally, therefore, the larger sex tends to be more nutritionally plastic than the smaller sex, and SSD increases with body size within a population [11]. This suggests that the mechanisms that generate SSD at least partially overlap with those that generate sex-specific nutritional plasticity.

Several adaptive hypotheses that explain why selection acts differently on males versus females may explain a relationship between SSD and SSP [9]. Under the adaptive canalization hypothesis [12], traits that are under strong stabilizing selection should be more canalized and less plastic than traits that are not. Thus, if there were strong stabilizing selection on the body size of one sex but not the other, then this could lead to differences in size plasticity. In contrast, the condition dependence hypothesis predicts that traits under strong directional sexual selection will exhibit greater sensitivity to environmental condition [13, 14] Consistent with this latter hypothesis is the observation that exaggerated traits used by males to compete for and attract mates are often typically highly plastic relative to the same trait in females (e.g. [14]). The condition-dependence hypothesis was formulated to explain the condition dependence of individual traits within the body and their response to directional sexual selection. Whether it can explain the correlation between SSP and SSD for the body as a whole, and where body size is not obviously subject to sexual selection (for example for female-biased SSD) is unclear.

While these adaptive hypotheses provide an ultimate evolutionary mechanism for a correlation between SSD and SSP, several studies have hinted at the proximate developmental mechanism linking the two phenomena. A study by Emlen et al [15] found that exaggerated male traits – specifically the horns of the beetle *Trypoxylus dichotomus –* are more insulin sensitive than other traits in the body. This accounted for both the trait’s increase in size in males relative to females, and their elevated nutritional plasticity. Consequently, the trait becomes more sexually dimorphic as trait size increases. In *Drosophila melanogaster*, where females are larger and more nutritionally plastic than males, SSD also appears to be regulated by the IIS pathway. This is based on three pieces of evidence: (1) SSD is eliminated in flies with suppressed IIS through hypomorphic mutation of the insulin receptor (*InR*) [16]; (2) well-fed females have higher IIS activity than males [17], and; (3) SSD requires sex-specific difference in the neurosecretory cells that produce insulin-like peptides (dILPs), the hormone that activates the IIS pathway [18]. Because SSD is eliminated in flies with a loss of InR activity, it follows that female body size is more sensitive to changes in insulin-signaling than male body size. Consequently, as for the beetle horn, SSP of body size in *Drosophila* may result from sex-specific differences in insulin-sensitivity. What is unknown, however, is whether the differences in nutritional plasticity between males and females is evident at the level of IIS/TOR-signaling activity itself.

To address this question, we looked at the expression of genes that regulate and are transcriptionally regulated by the IIS and TOR-signaling pathways across a nutritional landscape, where nutritional quality and quantity varies. In a previous study [19], we looked at the effect of changes in protein-to-carbohydrate ratio (diet quality) and total food concentration (diet quantity) on the size of the wing, maxillary palp, femur of the first leg and thorax. For all traits, female trait size was more sensitive to changes in either diet quality or quantity than male trait size. Intriguingly, however, these sex-specific differences could be attributed to the each trait’s size response to changes in carbohydrate versus protein concentration: males and female trait sizes were equally sensitive to changes in protein concentration, but only female trait size was detectibly sensitive to changes in carbohydrate concentration. Here we test the hypothesis that these observed sex-specific differences in body-size plasticity are reflected by corresponding differences in response to diet at the level of the IIS/TOR-signaling pathways.

## 3. Results

### 3.1. Female and male body size responds differently to changes in carbohydrates but not proteins

We found that, consistent with prior results [19], males and females differed in their response to dietary carbohydrate versus protein (Figure 1, Supplementary Figure 1, Table 1). Specifically, both male and female body size responded to changes in dietary protein as a negative quadratic, with body size increasing as protein concentration increased, but at a decreasing rate (Table 1). In contrast, only female body size responded to dietary carbohydrate concentration, this time as a positive quadratic, such that body size declined with increasing carbohydrate, but at a decreasing rate (Table 1). Correspondingly, including an interaction between sex and carbohydrate when modeling the relationship between diet and body size significantly improved model fit, while including an interaction between sex and protein did not further improve fit (Table 2). Thus male and female body size appears to respond differently to carbohydrate but not to protein. As a result, sexual size dimorphism (SSD) varied across the nutritional geometry landscape (Figure 1, Supplementary Figure 1).

**Table 1:**
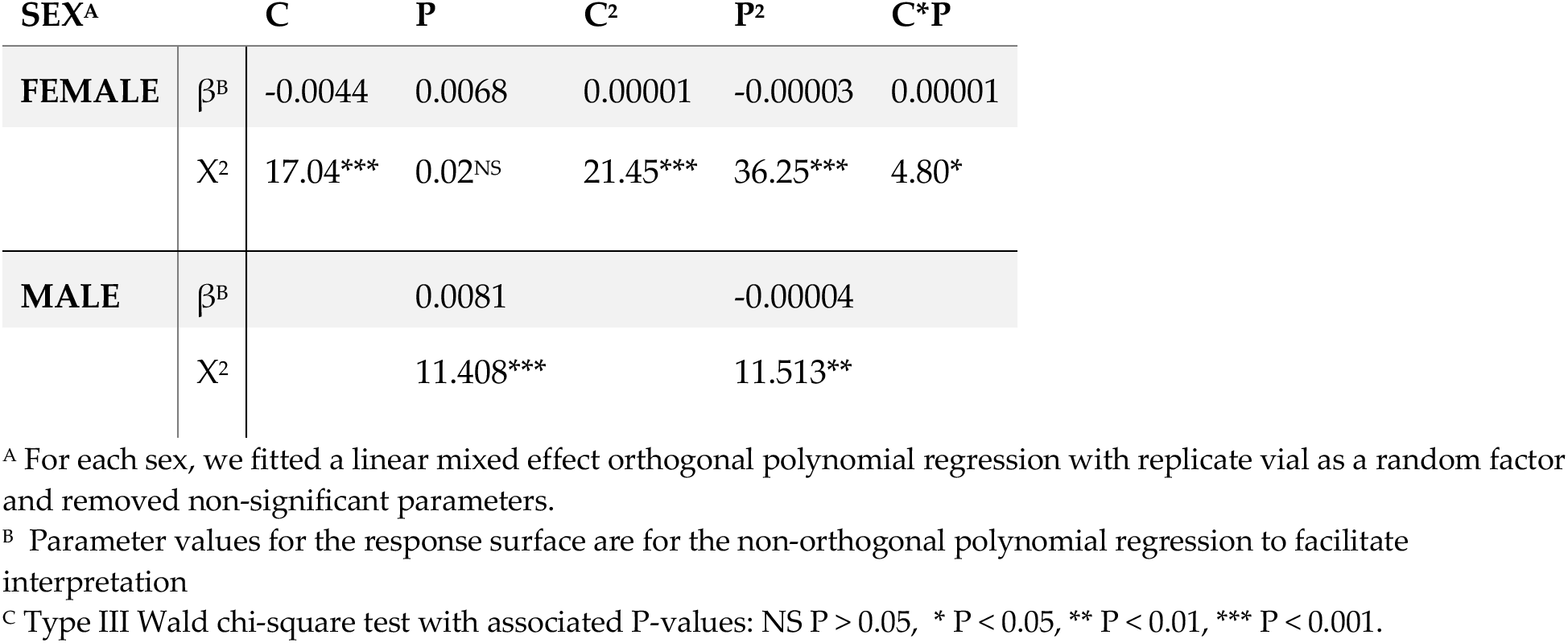
Effects of protein (P), carbohydrate (C), and their squares and product on body size in females and males.

| SEX<sup>A</sup> |  | C | P | C<sup>2</sup> | P<sup>2</sup> | C*P |
| --- | --- | --- | --- | --- | --- | --- |
| FEMALE | $\beta^B$ | -0.0044 | 0.0068 | 0.00001 | -0.00003 | 0.00001 |
| | $\chi^2$ | 17.04*** | 0.02<sup>NS</sup> | 21.45*** | 36.25*** | 4.80* |
| MALE | $\beta^B$ | | 0.0081 | | -0.00004 | |
| | $\chi^2$ | | 11.408*** | | 11.513** | |

<sup>A</sup> For each sex, we fitted a linear mixed effect orthogonal polynomial regression with replicate vial as a random factor and removed non-significant parameters.
<sup>B</sup> Parameter values for the response surface are for the non-orthogonal polynomial regression to facilitate interpretation
<sup>C</sup> Type III Wald chi-square test with associated P-values: NS $P > 0.05$ , \* $P < 0.05$ , \*\* $P < 0.01$ , \*\*\* $P < 0.001$ .

**Table 2:**
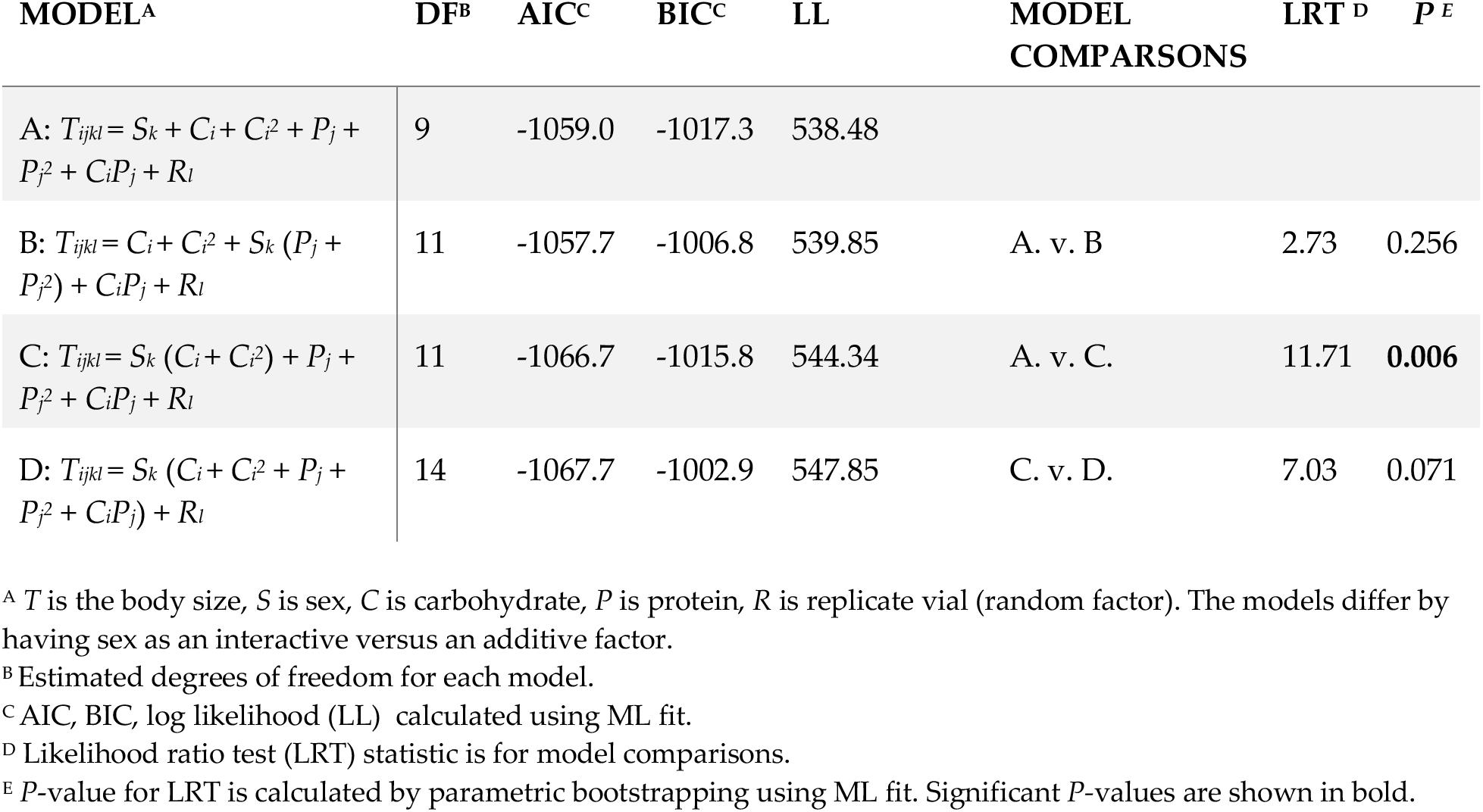
Effect of including sex as an interactive versus additive term when modeling the influence of diet on body size.

**Figure 1:**
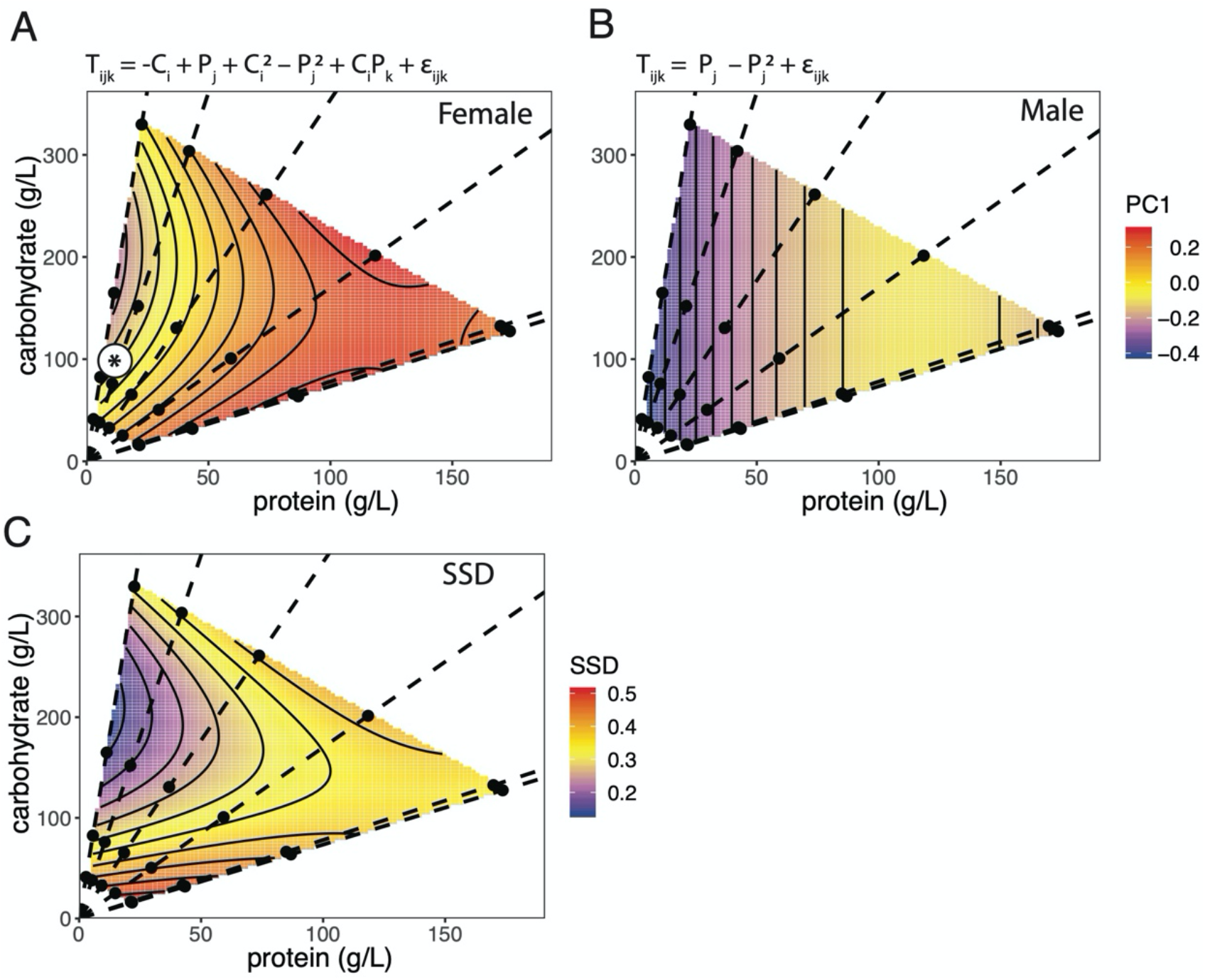
The effect of protein and carbohydrate concentration on female and male body size and sexual size dimorphism (SSD). (A, B) Surfaces show the fitted relationship between body size, carbohydrate level and protein level in female and male flies, based on the statistical model specified by the equation above each chart. T = body size, P = protein, C = carbohydrate, *ε* = error, subscripts refer to levels within each factor. (C) Surface shows the difference in female and male body size (SSD) across the same nutritional landscape, using fitted values from A and B. Points indicate diets tested and dotted lines connect diets with equal protein-to-carbohydrate ratios (1:14.6, 1:7.2, 1:3.5, 1:1.7, 1.3:1, 1.4:1). * indicates approximate composition of standard cornmeal-molasses medium. Corresponding thin-plate spline plots are shown in Supplementary Figure 1.

### 3.2. Female and male IIS/TOR gene expression responds differently to changes in diet

We collected expression data for eight genes that either regulate or are regulated by the IIS/TOR signaling pathway: insulin receptor (*InR*); eukaryotic translation initiation factor 4E binding protein (*4E-BP/Thor*); Drosophila insulin like peptides 2, 3, 5 and 8 (*dILP 2, 3, 5,8),* Absent, small, or homeotic discs-like protein (*Ash2L*), and *CG3071* (the Drosophila homolog of UTP15/SAW). Expression of both *InR* and *4E-BP* is negatively regulated by the IIS pathway [26], *dILPs 2,3,5* encode peptides that ostensibly activate the IIS pathway, while *dILP8* retards development by inhibiting the synthesis and release of ecdysone [27]. Finally expression of both *Ash2L* and *CG3071* has been shown to be negatively and positive regulated by *TOR* signaling, respectively [28]. Expression of only a few of these genes correlated directly with body size: *4EBP* and *dILP5* negatively and positively correlated with body size in females, respectively, while expression of *InR* negatively correlated with body size in males (Table 3). Nevertheless, a multivariate analysis of gene expression across all diets supported a significant interaction between the effects of sex and carbohydrate, as a linear factor, and sex and protein, as a quadratic factor, on gene expression (Table 4). We therefore explored how sex and diet affected the expression of individual genes. Specifically, we looked to see whether sex-specific responses in the expression of genes that either regulate or are regulated by the IIS/TOR-signalling pathways reflected the sex-specific response of body size to changes in diet.

**Table 3:** Coefficients and partial-eta-squared for multiple regression of body size against gene expression in males and females.

| GENE | FEMALE |  |  | MALE |  |  |
| --- | --- | --- | --- | --- | --- | --- |
| | Coefficient <sup>A</sup> | $\eta_p^2$ | *P* <sup>B</sup> | Coefficient | $\eta_p^2$ | *P* |
| *InR* | -0.014 | 0.004 | 0.638 | -0.038 | 0.168 | **0.047** |
| *4E-BP* | -0.123 | 0.147 | **0.003** | 0.018 | 0.013 | 0.593 |
| *Ash2L* | 0.014 | 0.003 | 0.660 | 0.019 | 0.034 | 0.391 |
| *CG3071* | -0.053 | 0.038 | 0.137 | -0.020 | 0.017 | 0.545 |
| *dILP2* | 0.039 | 0.018 | 0.308 | -0.019 | 0.072 | 0.206 |
| *dILP3* | -0.010 | 0.001 | 0.866 | -0.018 | 0.080 | 0.181 |
| *dILP5* | 0.054 | 0.071 | **0.042** | 0.047 | 0.146 | 0.066 |
| *dILP8* | -0.025 | 0.076 | 0.035 | <0.001 | <0.001 | 0.970 |

<sup>A</sup> Coefficients for the multiple regression $T_{ij...p} = A_i + B_j + \dots + H_p$ where *T* is body size (fitted values from the linear mixed model of the relationship between body size and diet) and *A* through *H* is the expression of the genes measured in our analysis.
<sup>B</sup> Significant *P*-values are shown in bold.

**Table 4:** Effect of including sex as an interactive versus additive term when modeling the influence of diet on the expression of multiple genes.

| MODEL<sup>A</sup> | DF <sup>B</sup> | VAR <sup>C</sup> | PILLAI<sup>D</sup> | F <sup>E</sup> | P <sup>F</sup> |
| --- | --- | --- | --- | --- | --- |
| $T_{ijk} = S_k + (C_i + P_j + P_j^2)$ | 92 | 0.53979 | | | |
| $T_{ijk} = S_k (C_i + P_j + P_j^2)$ | 89 | 0.51856 | 0.5284 | 2.1254 | **0.002** |
| $T_{ijk} = S_k (C_i + C_i^2 + P_j + P_j^2 + C_i P_j)$ | 85 | 0.51583 | 0.3791 | 1.0599 | 0.384 |

<sup>A</sup> $T$ is a matrix of expression levels across eight genes, $S$ is sex, $C$ is carbohydrate, $P$ is protein, $R$ is replicate vial (random factor). The models differ by having sex as an interactive versus an additive factor.
<sup>B</sup> Estimated degrees of freedom for each model.
<sup>C</sup> Generalized variance
<sup>E</sup> Pillai's Trace is against model specified in row above.
<sup>D</sup> Approximate $F$ -statistic is against model specified in row above.
<sup>E</sup> $P$ -value for LRT is calculated by parametric bootstrapping using ML fit. Significant $P$ -values are shown in bold.

#### Genes Regulated by IIS

Both *InR* and *4E-BP* expression responded to changes only in dietary protein but not carbohydrate (Figure 2). *InR* expression in males and females had a positive quadratic response to protein level, decreasing as protein level increased, but at a declining rate (Table 5). The expression of *InR* was lower across all diets in males than in females, but there was no significant interaction between the effects of sex and protein level on gene expression (Table 5). In contrast, *4E-BP* expression was sensitive to protein level only in females, decreasing linearly as protein increased, but was unaffected by diet in males (Table 4). Consequently, there was a significant interaction between the effect of sex and protein level on *4E-BP* expression (Table 5).

**Table 5:**
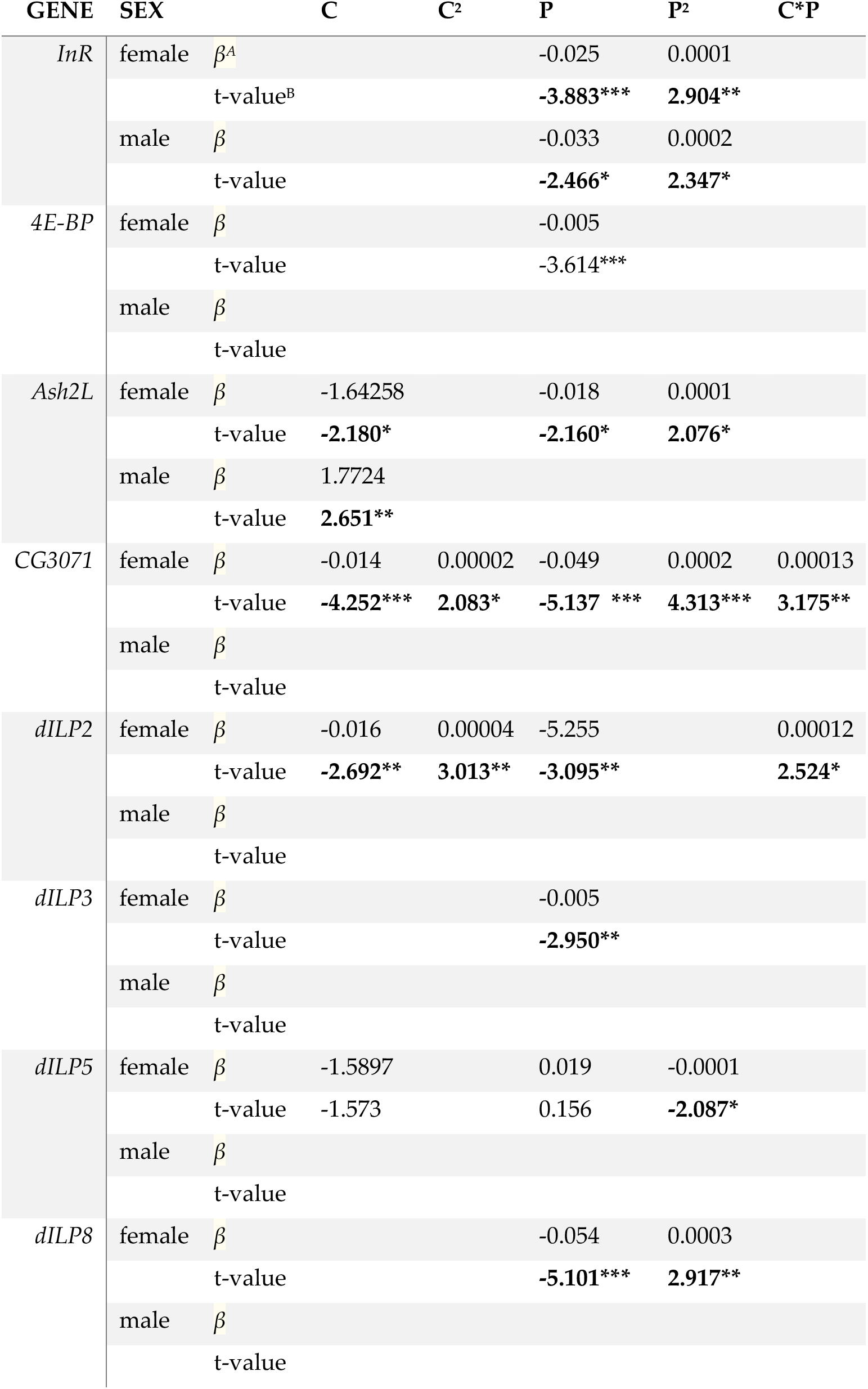
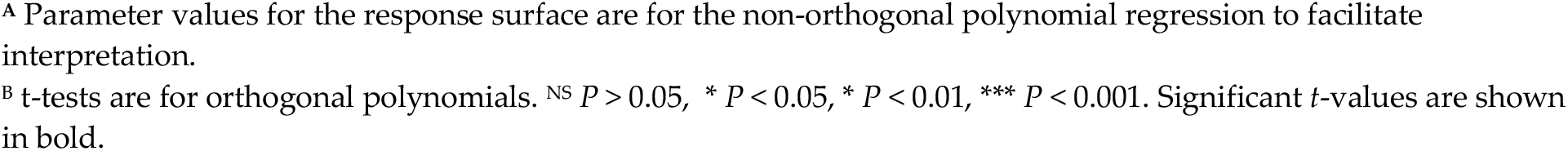
Effects of protein (*P*), carbohydrate (C), and their squares and product on gene expression in females and males.

| GENE | SEX |  | C | C<sup>2</sup> | P | P<sup>2</sup> | C*P |
| --- | --- | --- | --- | --- | --- | --- | --- |
| *InR* | female | $\beta^A$ | | | -0.025 | 0.0001 | |
|  |  | t-value<sup>B</sup> |  |  | **-3.883\*\*\*** | **2.904\*\*** |  |
| | male | $\beta$ | | | -0.033 | 0.0002 | |
|  |  | t-value |  |  | **-2.466\*** | **2.347\*** |  |
| *4E-BP* | female | $\beta$ | | | -0.005 | | |
|  |  | t-value |  |  | -3.614*** |  |  |
| | male | $\beta$ | | | | | |
|  |  | t-value |  |  |  |  |  |
| *Ash2L* | female | $\beta$ | -1.64258 | | -0.018 | 0.0001 | |
|  |  | t-value | **-2.180\*** |  | **-2.160\*** | **2.076\*** |  |
| | male | $\beta$ | 1.7724 | | | | |
|  |  | t-value | **2.651\*\*** |  |  |  |  |
| *CG3071* | female | $\beta$ | -0.014 | 0.00002 | -0.049 | 0.0002 | 0.00013 |
|  |  | t-value | **-4.252\*\*\*** | **2.083\*** | **-5.137 \*\*\*** | **4.313\*\*\*** | **3.175\*\*** |
| | male | $\beta$ | | | | | |
|  |  | t-value |  |  |  |  |  |
| *dILP2* | female | $\beta$ | -0.016 | 0.00004 | -5.255 | | 0.00012 |
|  |  | t-value | **-2.692\*\*** | **3.013\*\*** | **-3.095\*\*** |  | **2.524\*** |
| | male | $\beta$ | | | | | |
|  |  | t-value |  |  |  |  |  |
| *dILP3* | female | $\beta$ | | | -0.005 | | |
|  |  | t-value |  |  | **-2.950\*\*** |  |  |
| | male | $\beta$ | | | | | |
|  |  | t-value |  |  |  |  |  |
| *dILP5* | female | $\beta$ | -1.5897 | | 0.019 | -0.0001 | |
|  |  | t-value | -1.573 |  | 0.156 | **-2.087\*** |  |
| | male | $\beta$ | | | | | |
|  |  | t-value |  |  |  |  |  |
| *dILP8* | female | $\beta$ | | | -0.054 | 0.0003 | |
|  |  | t-value |  |  | **-5.101\*\*\*** | **2.917\*\*** |  |
| | male | $\beta$ | | | | | |
|  |  | t-value |  |  |  |  |  |

**Table 6:** Effect of including sex as an interactive versus additive term when modeling the influence of diet on the expression of individual genes.

| GENE | MODEL<sup>A</sup> | DF<sup>B</sup> | RSS<sup>C</sup> | F<sup>D</sup> | P<sup>E</sup> |
| --- | --- | --- | --- | --- | --- |
| *InR* | $T_{jk} = P_j + P_j^2$ | 148 | 100.209 | | |
| | $T_{jk} = S_k + P_j + P_j^2$ | 147 | 96.398 | 5.7500 | **0.0178** |
| | $T_{jk} = S_k (P_j + P_j^2)$ | 145 | 96.120 | 0.2098 | 0.811 |
| *4E-BP* | $T_{jk} = S_k + P_j$ | 153 | 52.219 | | |
| | $T_{jk} = S_k P_j$ | 152 | 48.604 | 11.305 | **0.001** |
| *Ash2l* | $T_{ik} = S_k + C_i$ | 152 | 80.009 | | |
| | $T_{ik} = S_k * C_i$ | 151 | 72.610 | 15.888 | **0.0001** |
| | $T_{ik} = S_k (C_i + P_j + P_j^2)$ | 147 | 68.459 | 2.229 | 0.069 |
| *CG3071* | $T_{ijk} = S_k + C_i + C_i^2 + P_j + P_j^2$ | 123 | 52.497 | | |
| | $T_{ijk} = S_k (C_i + C_i^2 + P_j + P_j^2)$ | 118 | 43.322 | 4.998 | **0.0003** |
| *dILP2* | $T_{ijk} = C_i + C_i^2 + P_j$ | 123 | 157.57 | | |
| | $T_{ijk} = S_k + C_i + C_i^2 + P_j$ | 122 | 156.47 | 0.8623 | 0.3550 |
| | $T_{ijk} = S_k (C_i + C_i^2 + P_j + C_i P_j)$ | 119 | 152.96 | 0.9097 | 0.4386 |
| *dILP3* | $T_{jk} = P_j$ | 152 | 136.38 | | |
| | $T_{jk} = S_k + P_j$ | 151 | 134.35 | 2.2632 | 0.1346 |
| | $T_{jk} = S_k P_j$ | 150 | 134.35 | <0.0001 | 0.9803 |
| *dILP5* | $T_{ijk} = C_i + P_j + P_j^2$ | 148 | 147.40 | | |
| | $T_{ijk} = S_k + C_i + P_j + P_j^2$ | 147 | 142.85 | 4.6873 | **0.0320** |
| | $T_{ijk} = S_k (C_i + P_j + P_j^2)$ | 144 | 139.80 | 1.0458 | 0.37431 |
| *dILP8* | $T_{jk} = S_k + P_j + P_j^2$ | 121 | 205.98 | | |
| | $T_{jk} = S_k (P_j + P_j^2)$ | 119 | 190.40 | 4.8693 | **0.0093** |

<sup>A</sup> $T$ is a expression level of gene, $S$ is sex, $C$ is carbohydrate, $P$ is protein. The models differ by having sex as an interactive versus an additive factor. The models chosen are the simplest where one or either sex have significant parameter values.
<sup>B</sup> Degrees of freedom for each model.
<sup>C</sup> Residual Sum Square
<sup>D</sup> $F$ -statistic is against model specified in row above.
<sup>E</sup> Significant P-values are shown in bold.

**Figure 2:**
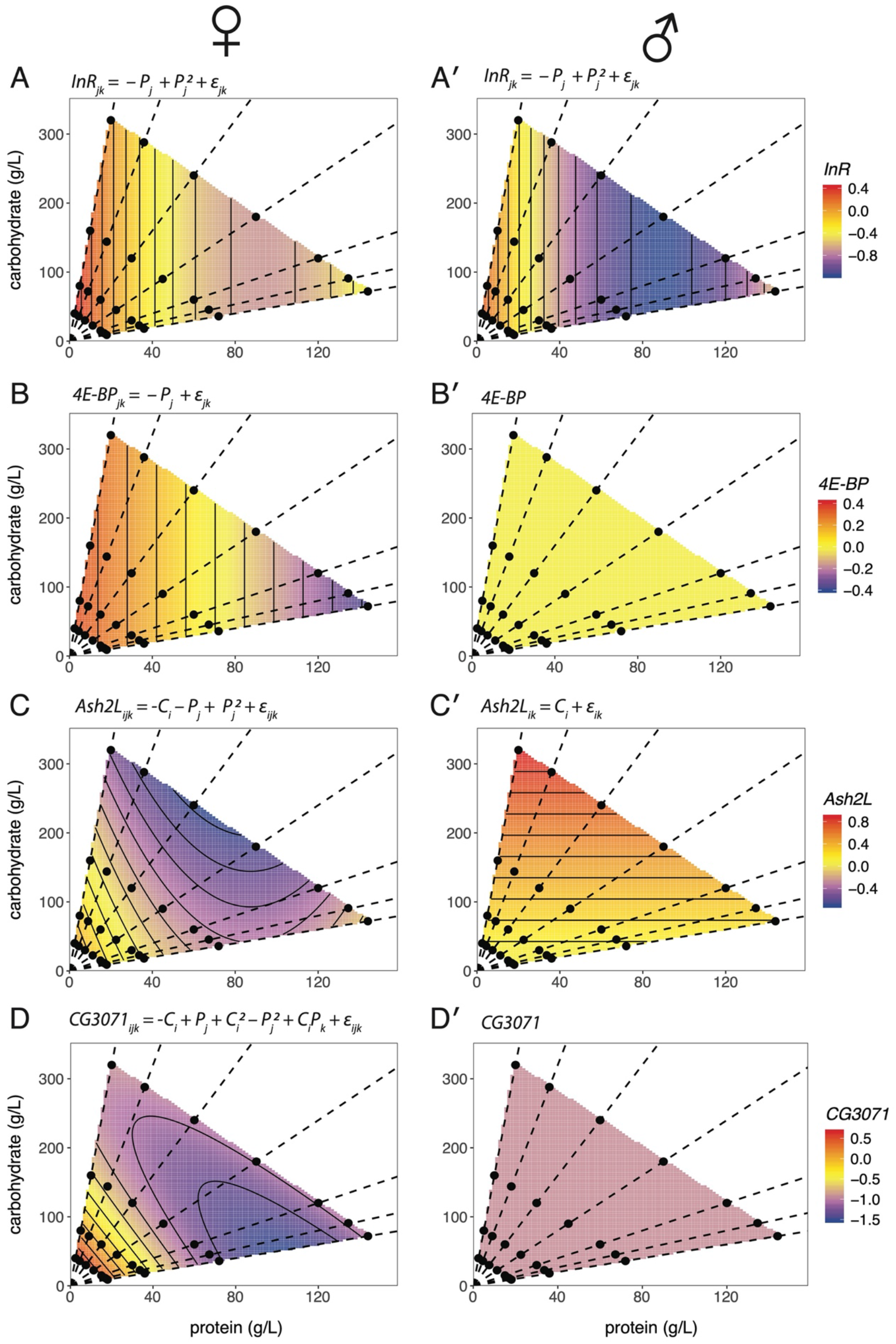
The effect of protein and carbohydrate concentration on the expression of IIS and TOR transcriptionally-regulated genes in females and males. Surfaces show the fitted relationship between gene expression, carbohydrate level and protein level in female and male flies, based on the statistical model specified by the equation above each chart (Table 4). *P* = protein, *C* = carbohydrate, *∊* = error, subscripts refer to levels within each factor. Expression of (A, A′) *InR*, and (B, B′) *4E-BP*, both negatively regulated by the activity of the IIS via the Forkhead transcription factor FOXO. (C, C′) Expression of *Ash2L*, ostensibly negatively regulated by the activity of TOR signaling. (D, D′) Expression of *CG3071*, ostensibly positively regulated by the activity of TOR signaling. Points indicate diets tested and dotted lines connect diets with equal protein-to-carbohydrate ratios (1:16, 1:8, 1:4, 1:2, 1:1, 2:1). Corresponding thin-plate spline plots are shown in Supplementary Figure 2.

**Figure 3:**
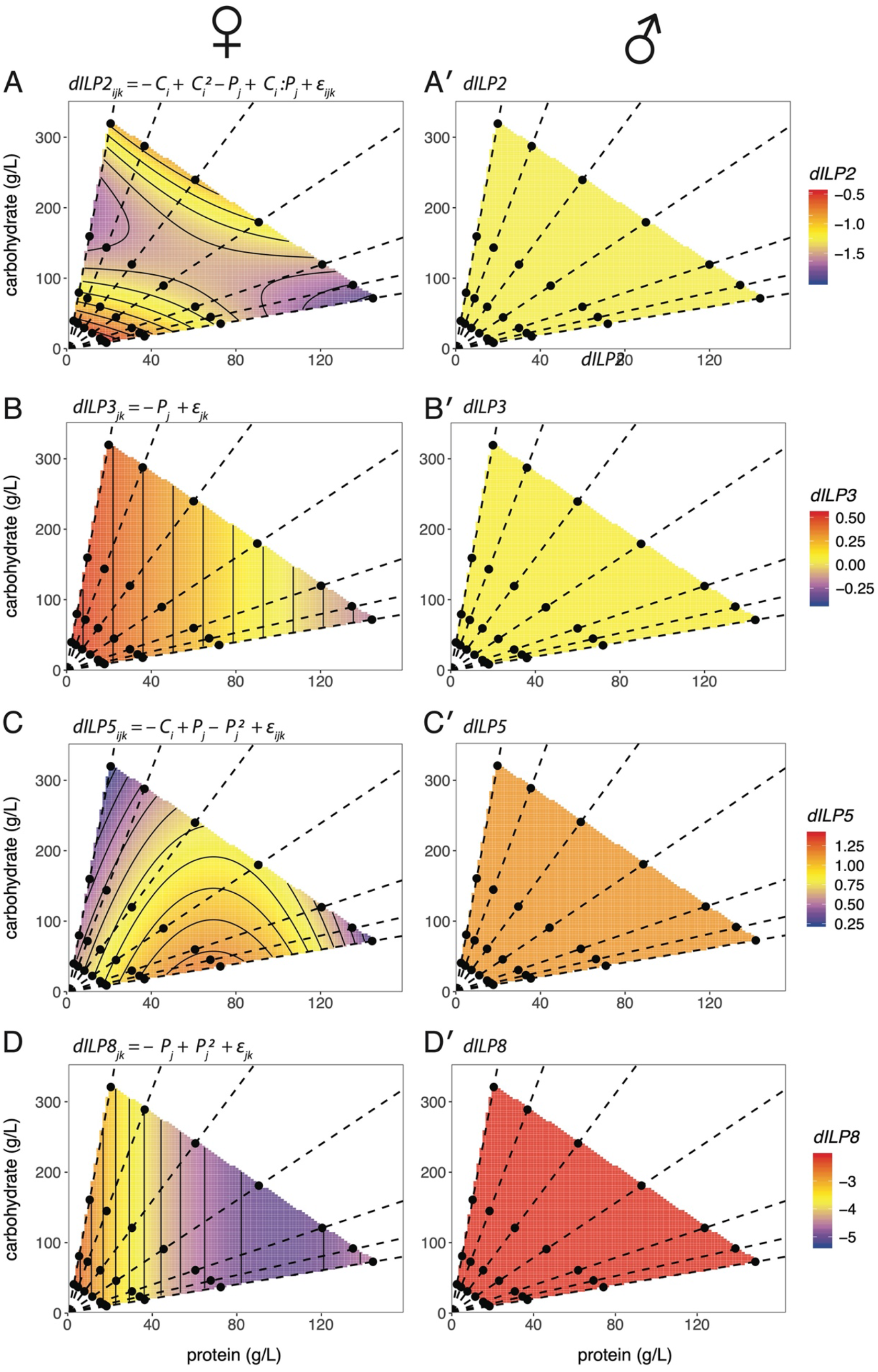
The effect of protein and carbohydrate concentration on the expression of dILPs in females and males. Surfaces show the fitted relationship between gene expression, carbohydrate level and protein level in female and male flies, based on the statistical model specified by the equation above each chart (Table 4). *P* = protein, *C* = carbohydrate, *ε* = error, subscripts refer to levels within each factor. (A) *dILP2*. (B) *dILP3.* (C) *dILP 5.* (D) *dILP8*. Points indicate diets tested and dotted lines connect diets with equal protein-to-carbohydrate ratios (1:16, 1:8, 1:4, 1:2, 1:1, 2:1). Corresponding thin-plate spline plots are shown in Supplementary Figure 3.

#### Genes Regulated by TOR Signaling

Expression of both *Ash2L* and *CG3071* had a complex and sex-specific response to diet. Carbohydrate significantly affected *Ash2L* expression as a linear relationship, but in opposite directions in either sex: Increased carbohydrate increased *Ash2L* expression in males but decreased *Ash2L* expression in females (Figure 2, Table 4). *Ash2L* expression was not affected by protein in males, but had a positive quadratic response to protein in females, decreasing as protein increased but at a declining rate. There was therefore a significant interaction between sex and diet on the expression of *Ash2L,* although only between sex and carbohydrate and not sex and protein (Table 5). *CG3071* also showed a sex-specific expression response to diet. In females, *CG3071* expression had a negative quadratic response to protein level (increasing at a declining rate as protein increased), a positive quadratic response to carbohydrate level (decreasing at a declining rate as carbohydrate increased), with a significant carbohydrate: protein interaction (Table 4). In contrast, *CG3071* expression was not affected by diet in males, which lead to a significant diet-by-sex interaction effect on *CG3071* expression (Table 5).

#### Genes That Regulate IIS/TOR Signaling

The expression of *dILP2* showed a significant response to diet in females, declining linearly with increasing protein, and at a decreasing rate with increasing carbohydrate, as a positive quadratic. There was also a significant carbohydrate-by-protein interaction effect on *dILP2* expression in females. There was no significant effect of diet on *dILP2* expression in males, although this appears to be a consequence of low statistical power, due to large amount of within-diet variation in *dILP2* expression among male samples compared to females sample. Fitting a one-way ANOVA of *dILP2* expression against diet, where diet is each protein-to-carbohydrate ratio at each concentration, the residual variance (a measure of average variance within diets) was significantly higher in males than in females, by a ratio of 8.68 (Supplementary Table 1). Correspondingly, we had very low power (0.22) to detect a significant sex-by-diet interaction for *dILP2* expression, based on the observed effect size (Cohen’s *f^2^* = 0.023, see Supplementary Material for details). Further, there was no evidence that expression of *dILP2* was different between males and females, independent of diet (Table 5).

The expression of *dILP3* decreased linearly with protein in females, but there was no detectable effect of diet on *dILP3* expression in males (Figure 2, Table 4). There was also no detectable difference in expression between the sexes (Table 5). As for *dILP2*, variation in expression of *dILP3* among samples within diets was significantly higher in males than in females (Supplementary Table 1), and a power analysis gave a power of only 0.05 to detect a sex-by-protein interaction on *dILP3* expression, based on the observed effect size (Cohen’s *f^2^* <0.001).

The expression of *dILP5* was only marginally affected by diet in females, with a significant negative quadratic effect of protein. We could detect no significant effect of either carbohydrate or protein on *dILP5* expression in males. As for *dILP2* and *3*, *dILP5* expression was more variable among male samples within diets than among female samples (Supplementary Table 1), and a power analysis gave a power of only 0.23 to detect a sex-by-protein interaction on *dILP5* expression, based on the observed effect size (Cohen’s *f^2^* = 0.022). Nevertheless, there was sex-specific expression independent of diet, such that *dILP5* expression was higher in males than in females (Table 5).

The expression of *dILP8* responded as a positive quadratic in response to protein in females, declining as protein increased at a decreasing rate, but did not show any response to any aspect of diet in males (Figure 2, Table 4). The lack of a corresponding response to protein in males was not due to a lack of statistical power: a power analysis suggested that the power to detect the female effect size of protein on *dILP8* expression (Cohen’s *f^2^* = 0.46) in males was 0.99. Further, *dILP8* expression was not more variable among male samples within diets than among female samples (Supplementary Table 1). Consequently, we detected a significant sex-by-protein interaction on *dILP8* expression (Table 5).

### 3.3 dILP protein levels in the brain increase with increasing food concentration

Our finding that *dILP* expression in females was generally higher in larvae fed on lower food concentrations was surprising, given that dILPs (apart from dILP8) are canonically positive regulators of growth in response to nutrition. We therefore explored the relationship between diet, *dILP* expression and the levels of dILP peptide in the insulin-producing cells (IPCs) of the brain. Staining for dILP2 and dILP5 peptide was higher in larvae fed a higher food concentration (360g/l) versus a lower food concentration (45g/l), with a 1:2 protein: carbohydrate ratio (Figure 4).

**Figure 4:**
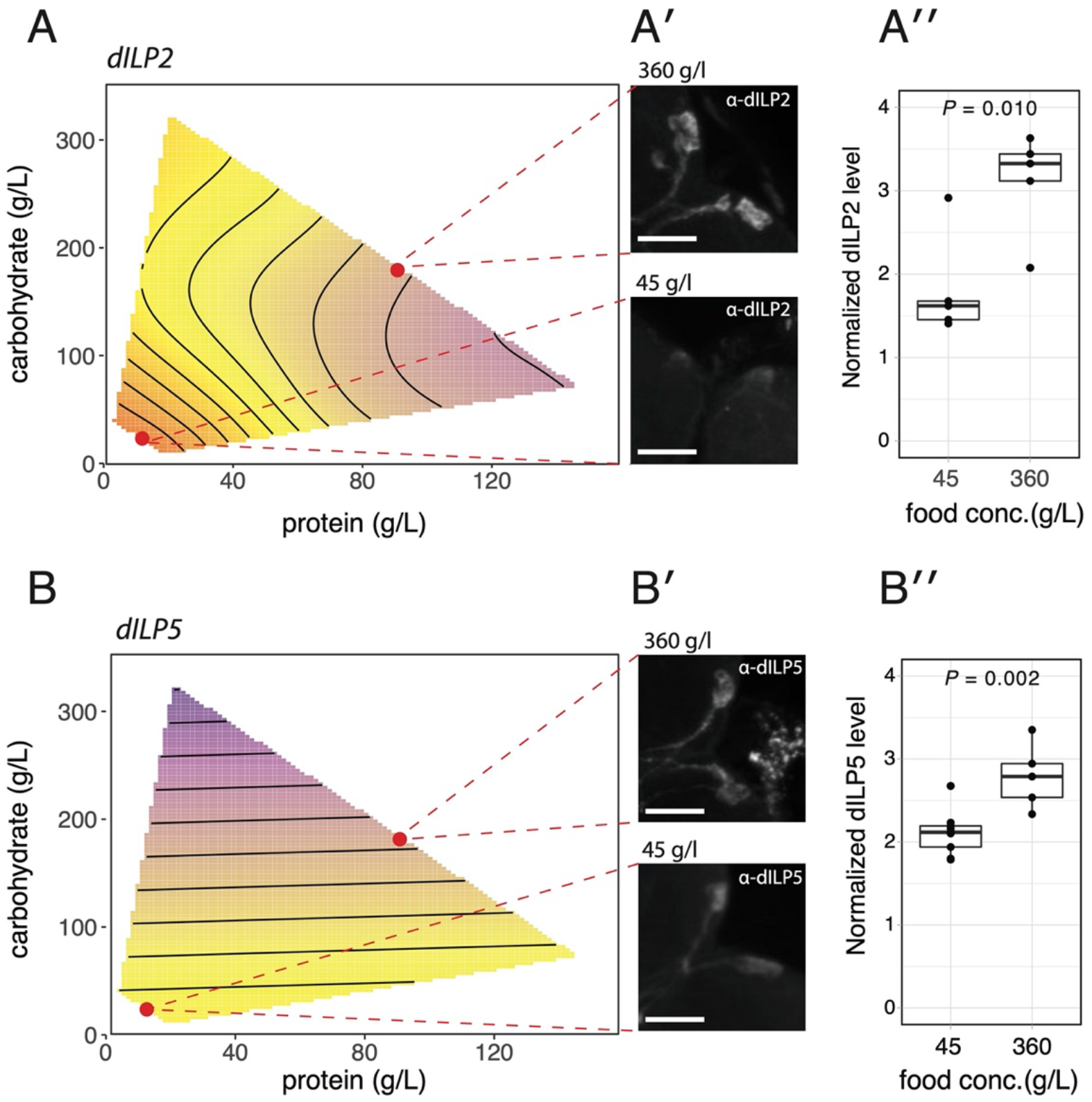
The effect of diet on dILP2/5 expression and peptide levels in the brain in females and males. (A,B) The relationship between diet and *dILP2* and *dILP5* expression. Points indicate 45 and 360 g/l food concentration at 1:2 protein: carbohydrate ratio. (A′, B′) Representative images of dILP2 and dILP5 levels in the insulin-producing cells (IPCs) of female larvae reared on different diets. (A″, B″) Mean normalized level of dILP2 and dILP5 in the IPCs of larvae reared on different diets. *P-*values are for a pooled t-test.

## 4. Discussion

Previous studies indicate that in *Drosophila*, body size is more nutritionally plastic in females than in males, a phenomenon referred to sex-specific plasticity (SSP) [19]. Nutritional plasticity is canonically regulated by the insulin/IGF-signaling and TOR signaling pathways in *Drosophila* and most other animals. We hypothesized, therefore, that differences in nutritional plasticity between the sexes reflects sex-specific differences in the activation and activity of these pathways. In this study, we applied a nutritional geometry framework to test this hypothesis, and explored the sex-specific effects of protein and carbohydrates on body size and the expression of genes that activate and are transcriptionally regulated by IIS/TOR. Our results indicate that, broadly, gene expression is more nutritionally sensitive in females than in males, although the effect of nutrition on the expression of IIS and TOR-signaling genes is both quantitively and qualitatively different between males and females.

The simplest hypothesis to account for sex-specific differences in the nutritional plasticity of body size in *Drosophila* is that females have the same response to nutrition as males, but at a higher sensitivity. Under this hypothesis, the response surfaces of body size and gene expression across a nutritional landscape of varying protein and carbohydrate concentrations should be the same shape in both sexes, but with lower gradients in males. This would be evident as lower parameter values in males versus females. This is not supported by the data, however. In particular, body size in males shows more or less the same sensitivity to changes in protein concentration as females, but has no detectable sensitivity to changes in carbohydrate concentration. This in turn suggests that an important aspect of sex-specific nutritional plasticity in *Drosophila* is differences in the response to variation in dietary carbohydrates.

The best understood regulator of body size with respect to nutrition are the insulin/IGF-signaling and TOR-signaling pathways [29, 30]. The IIS pathway is activated by circulating dILPs, some of which are released in a nutrition-dependent manner [27]. The TOR-signaling pathway is in part regulated by IIS, but also responds directly to circulating amino acids [31–33]. Because the SSP of body size in flies appears to primarily driven by a differential response to carbohydrates but not to protein, we might expect that the effect of diet on body size is mediated primarily by IIS rather than TOR-signaling. Our data do not support this hypothesis. We used the expression of *InR* and *4E-BP* as a measure of IIS activity: Both are transcriptionally regulated by the forkhead transcription factor FOXO, which is activated when nutrition and IIS activity is low [26, 34]. If sex-specific plasticity is mediated by IIS then we would expect to see sex-specific differences in *Inr* and *4EBP* expression in response to changes in carbohydrate but not to protein. In both males and females, however, *InR* expression responded only to protein level – and to the same extant – but did not respond to carbohydrate level. Expression of *4E-BP* only responded to protein level in females, but did not respond to carbohydrate level in either sex. Collectively, variation in the expression of *4E-BP* and *InR* were the strongest correlates with body size in females and males respectively (Table 3), consistent with the hypothesis that the IIS is the major regulator of body size with respect to nutrition in *Drosophila*. However, the expression of these genes do not reflect sex-specific differences in the effect of carbohydrate on body size, and so do not explain sex-specific plasticity.

In contrast to *InR* and *4E-BP*, the expression of *Ash2L* and *CG3071* both responded to carbohydrates in a sex-specific manner. However, the response is not consistent with a previous report on the regulation of the these genes’ expression by TOR-signaling [28]. This study reported that *Ash2L* is negatively regulated by TOR-signaling while *CG3071* is positively regulated. If TOR-signaling increases with protein, then we would expect *Ash2L* expression to decrease correspondingly, which is true in females. However, *Ash2L* expression also decreases with increasing carbohydrate, which is difficult to reconcile with the negative effects that carbohydrate has on female growth. Even more challenging to interpret is the observation that *Ash2L* expression increases with carbohydrate but not protein in males, even though male do not appear to have a growth response to carbohydrate. Thus the relationship between *Ash2L* expression and body size is not a simple one. The expression pattern of *CG3071* is similarly unclear. While *CG3071* also has a carbohydrate and protein response in females, this response is qualitatively similar to the response of *Ash2L,* which is not expected if TOR-signaling positively regulates *CG3071* expression but negatively regulates *Ash2L* expression. Thus the role that TOR signaling plays in regulating SSD and SSP is equivocal based on our data. Problematically, the mechanism by which TOR-signaling regulates *Ash2L* and *CG3071* expression has not been fully explored, which makes interpreting changes in *Ash2L* and *CG3071* expression with diet even more challenging.

Our data suggests a complex relationship between sex, diet, *dILP* expression and dILP retention levels in the IPCs. In females, there were stark differences in the pattern of expression among the dILPS. Of all the dILPs, *dILP5* expression was most strongly correlated with body size, increasing with increasing protein level but decreasing with increasing carbohydrate level. We also found that dILP5 levels in the IPCs were lower in larvae fed on lower food concentrations. Thus, chronic reductions in nutrition reduce *dILP5* expression and dILP5 IPCs level. This is consistent with dILP5 being a positive regulator of growth in response to nutrition [35]. The negative effect of low nutrition on *dILP5* expression has been observed in previous studies [30, 35, 36]. However, at least one of these studies reported an increase in dILP5 retention in the IPCs of larvae that were either starved for 24 hours or reared on a low protein diet since birth [36], which is inconsistent with our data. The low protein diet in this study was generated by reducing the amount of protein from ~16g/l to ~0.8g/l, while keeping the concentration of carbohydrate at ~60g/l, thereby decreasing the protein-to-carbohydrate ratio. In our study we reduced overall food concentration from 360g/l to 45g/l while retaining a 1:2 protein-to-carbohydrate ratio. It is possible, therefore, that the retention of dILP5 in the IPCs in the earlier study is due to a low protein-to-carbohydrate ratio. Indeed, a second earlier study showed that a chronic high sugar diet with low protein-to-carbohydrate ratio also increased dILP5 levels in the IPCs [37], although this time accompanied by an increase in *dILP5* expression. Thus the relationship between *dILP5* expression and dILP5 levels in the IPCs may depend not only on the overall food concentration, but also on food composition, in particular the protein-to-carbohydrate level.

Previous studies suggest that *dILP2* expression is unaffected by reduced nutrition [30, 36] (although see supplementary data in [38]), but that dILP2 peptides are retained in the IPCs in larvae that have been starved 24h or are reared on a low protein diet [36, 39, 40]. A high sugar diet is also associated with increased retention of dILP2 peptide in the IPCs, but with an increase in *dILP2* expression [37]. In our study, however, we found that *dILP2* expression declined with an increase in both protein and carbohydrate, and that dILP2 peptide was lower in the IPCs of larvae fed on lower food concentrations. Our observed expression of *dILP3* was also inconsistent with previous studies that found that acute starvation reduced expression [30], while rearing on a high protein diet increased expression [41]. In contrast, we found that, as for *dILP2*, *dILP3* expression declined with increasing protein, although was unaffected by carbohydrate level, which is surprising given that circulating sugars promote the release of dILP3 from the IPCs [40]. Our finding that *dILP2* and *dILP3* expression is lowest on low protein diets that restrict growth, and that dILP2 peptides levels are higher when *dILP2* expression is lower, suggests that dILP2 and dILP3 may negatively regulate their own expression, which is a common characteristic of hormone regulation.

Finally, the expression of *dILP8* showed a similar pattern to *dILP3,* increasing with a decrease in protein concentration. dILP8 is involved in regulating growth and developmental timing through its inhibitory effects on ecdysone synthesis [42, 43]. It is released from imaginal discs in response to damage or growth perturbation, as well as showing periodic changes in expression throughout development. Ecdysone synthesis is also inhibited by low nutrition early in the third larval instar, leading to a delay in metamorphosis [44]. The observation that females on low protein diets also show elevated levels of *dILP8* expression, suggests that dILP8 may play a role in regulating growth and/or development in response to nutrition, at least in females.

We did not detect an effect of diet on the expression of any of the *dILP*s in male larvae. This did not, however, translate into a significant interaction between diet and sex on *dILP* expression, and therefore does not help explain the sex-specific differences in nutritional plasticity of body size. This was because variation in *dILP 2,3,5* and *8* expression among samples within diets was significantly higher for male samples than for females samples (Supplementary Table 1), reducing the statistical power to detect diet-by-sex interactions. The only exception was for *dILP8,* such that *dILP8* expression was significantly more plastic in females than in males. Nevertheless, *dILP8* expression in females did not correlate with carbohydrate level, and so is unlikely to explain the differential effect of carbohydrate on body size in females versus males. The elevated variation in *dILP* expression among male samples was not seen for expression of *4eBP*, *InR*, *Ash2L*, or *CG3072*, and is therefore not likely due to sample degradation. It is also not due to lower levels of expression in males relative to females. It is possible, therefore, that *dILP* expression levels are more developmentally dynamic in males then females, and that we are capturing aspects of that instability by measuring gene expression at only a single point in development.

Collectively, there appears to be no simple explanatory relationship between the sex-specific nutritional geometry of body size with the sex-specific nutritional geometry of IIS/TOR-signaling gene expression. Further, many of our findings do not replicate what has been reported in previous studies. An earlier study looking at *dILP* expression in adults across a nutritional landscape saw similarly complex and non-intuitive relationships between diet and gene expression [23]. What is very clear from both studies is that the protein-to-carbohydrate ratio affects the transcriptional response of IIS/TOR-signaling genes to changes in total diet, and our study indicates that it does so in a sex-specific way. This has important implications for interpreting those studies that have looked at changes in the expression of IIS/TOR-signaling genes in response to changes in nutrition. First, many studies simply dilute diet to reduce overall nutrition. Problematically, while most *Drosophila* labs use ‘standard’ food recipes, the use of different types of yeast, cornmeal, mollasses *etc* means that the composition of these diets may be unique to most research groups [45]. Our data indicate that the effect of diet dilution on both body size and IIS/TOR-signaling gene expression – and potentially the activity of the IIS/TOR-signaling pathways – will depend on the P:C ratio of that specific diet. Second, these responses are sex-specific, and studies that do not consider sex may come to different conclusions than studies that do. Third, the relationship between *dILP* expression, dILP retention and diet is similarly complex, and may also depend on the composition of the diet being manipulated.

One important caveat with our, and almost all other studies of IIS/TOR-signaling gene expression during development, is that we measured expression at a single developmental time point, at the very beginning of larval wandering. Previous studies have shown that the activity of the IIS and TOR-signaling pathways change dynamically during development, and that different dILPs are expressed at different life stages. This may also account for differences between our and other studies of the effects of nutrition on IIS/TOR-signaling gene expression. Ideally, one would like to conduct a multidimensional study of gene expression across both time and a nutritional landscape, and to tie this with a corresponding study of dILP levels both in the IPCs and circulating in the hemolymph. Such a study would not only help elucidate the relationship between IIS/TOR-signaling and the sex-specific effects of nutrition on body size, but also help us understand better how nutrition regulates the transcription, translation, storage and release of dILPs.

## 5. Conclusions

Our study provides a foundation for future work that looks at the sex-specific effects of diet quantity and quality on body size and on the developmental mechanisms that regulate body size. Our data show significant differences in the effect of diet on gene expression in males and females, although they do not suggest a simple explanatory relationship between gene expression and body size. Future studies on sex-specific plasticity in *Drosophila*, and other animals, should therefore consider not only the nutritional geometry of gene expression, but also of protein levels, pathway activity and other developmental parameters such as growth rate and growth duration, ideally at multiple time points throughout development.

## 6. Materials and Methods

The goal of the study was to test the hypothesis that sex-specific differences in body-size plasticity across a nutritional landscape in *Drosophila* correspond to sex-specific differences in the response to diet at the level of the IIS/TOR-signaling pathways, using the expression of genes that regulate or are regulated by IIS/TOR-signaling to assay pathway activity.

### 6.1. Fly Stocks and Maintenance

Flies used for the nutritional geometry of body size were as described in [19]. Flies used for the nutritional geometry of gene expression were an isogenic line of white-eyed but otherwise wild-type flies (VDRC 60000). Flies were maintained as a stock at 17°C on standard on standard media of 45 g of molasses, 75 g of sucrose, 70 g of cornmeal, 20 g of yeast extract, 10 g of agar, 1100 ml of water, and 25 ml of a 10% Nipagin solution per liter of fly food, under constant light.

### 6.2. Diet Manipulation for Nutritional Geometry of Body Size

Flies were reared on 24 diets: six protein-to-carbohydrate ratios (1:14.6, 1:7.2, 1:3.5, 1:1.7, 1.3:1, 1.4:1), at four food concentrations: (45, 90, 180 and 360 g l−1), with six replicate vials per diet, as described in [19]. We measured four non-genital traits in each fly: the length of the first femur, the area of the wing, the area of the maxillary palp and the length of the thorax (see [19] for details). Data were collected from, on average, 17 flies from each sex at each diet, with no more than 30 flies coming from any one vial. To assay the effect of diet on overall body size, we conducted a principle component analysis on all log-transformed non-genital morphological data from all flies, and used each fly’s value for the first principle component (PC1) as a proxy for overall body size [20, 21].

### 6.3. Diet Manipulation for Nutritional Geometry of Gene Expression

Flies were reared on 28 different diets: seven protein-to-carbohydrate ratios (1:16, 1:8, 1:4, 1:2, 1:1, 1.5:1 and 2:1) at four food concentrations (45, 90, 180, and 360 g/l). To generate the diets we made an instant-yeast solution (Lesaffre SAF-Instant Red: 44% protein, 33% carbohydrate) and a sucrose solution (Mallinokrodt, Paris, Kentucky) at each food concentration. We also added 0.5% agar, 10% nipagen (10% p-hydroxy benzoic acid methyl ester in 95% ethanol) and 1% propionic acid (Sigma Life Science, St. Louis, Missouri) to each solution, to control consistency and prevent fungal growth. The solutions were autoclaved and mixed at different ratios to achieve the desired protein-to-carbohydrate ratios for each diet (supplemental Table 1). 10 ml of each of the 28 different diets were aliquoted into separate 25 x 95 mm vials.

Populations of 50 flies were placed in mating cages with 60mm petri dishes of standard lab food. The flies were allowed to lay eggs for 24 hrs. The adult flies were removed and the eggs were left on the petri dish for another 24hrs until the larvae had hatched. The first instar larvae were then transferred in groups of 50 into each food vial. There were at least five replicate vials for each diet.

All cultures were maintained at 25°C at constant light under 60-70% humidity. Larvae were collected from the food vials when they had just initiated wandering, to ensure that all larvae were collected at the same developmental stage regardless of diet. All larvae were sexed and flash-frozen to −80°C within five minutes of being removed from the vial. The larvae were stored at −80°C until they were used for RNA extractions.

### 6.4. Gene Expression Quantification

We used qPCR to assay gene expression in male and female larvae collected at the beginning of larval wandering. Larvae were pooled into cohorts of 15 individuals, and we collected 3 cohorts of each sex for each of the 28 diets, with the exception of the 1:16 360g/L diet, which only generated enough larvae for 2 cohorts of each sex. For each sample, RNA was extracted using Trizol (Invitrogen, Grand Island, NY, USA) and treated with DNase 1 (Invitrogen) before being reverse-transcribed with High Capacity cDNA Reverse Transcription Kit synthesis according to manufactures instructions (Thermo Fischer Scientific, Carlsbad, CA, USA). We used SYBR Green PCR master mix (Thermo Fischer Scientific, Carlsbad, CA, USA) to conduct qPCR using a CFX Connect Real-time System (Bio-Rad Laboratories, Inc., Hercules, CA). mRNA abundance was calculated using the standard curve method [22]. Relative gene expression was calculated by comparing the measured mRNA abundance of the gene of interest to that of ribosomal protein 49 (RP49). Prior studies have indicated that RP49 is insensitive to changes in nutritional conditions [23]. In two samples (1:16 180g and 1:4 45g male), gene expression levels were an order of magnitude above all other samples across all amplicons. Data from these samples was therefore excluded from subsequent analysis. The complete protocol is provided in the supplementary material, along with primer sequences.

### 6.5. dILP2 and dILP5 Levels in the Brain

The following antibodies were used: dILP2 rat (1:800); dILP5 rabbit (1:800) (both gifts of P. Leopold); AlexaFluor 594 Donkey anti-Rat IgG (1:500); AlexaFluor 594 Goat anti-Rabbit IgG (1:500) (both from Life Technologies).

Larvae were reared on either 45g/l or 360g/l at a protein:carbohydrate ratio of 1:2, as described above. Brains were dissected in 1X PBS on ice from wandering female third instar larvae. The brains were fixed for 30 minutes in 4% paraformaldehyde and washed in 0.3% Triton-X in PBS (PBT) overnight at 4oC. They were then washed for 1 hr in 2% NGS in 0.2% I-Block in PBT (BBT/NGS) and incubated with primary antibody diluted in BBT/NGS overnight at 4oC. Subsequently, brains were washed four time in 0.2% BSA/I-Block in PBT (BBT), 15 minutes per wash. Brain were then incubated in secondary antibody at room temperature for 2 hours. Finally, brains were washed in four times in PBT for 15 minutes per wash and mounted with vector shield + DAPI.

We acquired images using Olympus Fluoview FV101 confocal microscope. For both 45g/L and 360g/L dILP2 images, 35% DAPI and 35% Alexa 594 intensity settings were used. For both 45g/L and 360g/L dILP5 images, 30% DAPI and 30% Alexa 594 intensity settings were used. We used a 1.5 micron step size to capture a Z-stack of images for each IPC (one per brain), which was flattened to a single image using a maximum-intensity projection in FIJI software. The intensity of the IPC in both the Alexa 594 and DAPI channel was then used to generate a normalized measure of the level of dILP2 or dILP5 in each brain. We repeated this for five brains for dILP2 and dILP5 at each diet level (nine for dILP5 at 45g/l). dILP levels were then compared between diets using a standard pooled *t*-test.

### 6.6. Statistical Analysis

Nonlinear response surfaces are routinely modelled using the second-order polynomial regression [19, 23–25]. To determine the effect of protein and carbohydrate level on body size or gene expression we therefore fit the model:

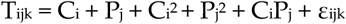

where *T* is trait (body size or expression level), *C* is carbohydrate concentration, *P* is protein concentration and *∊* is error (subscripts are levels within variables). Models for body size included replicate vial as a random factor and were fit using *lmer* in the *lme4* package in *R.* Models for expression level were fit using *lm* in the *base* package of *R.* For each trait, the significance of each parameter was tested with a type III ANOVA using the *Anova* function in the *car* package for standard models and the *summary* function in the *lmerTest* package for mixed models. Any non-significant parameters were subsequently removed from the model before the data were reanalyzed using the simplified model. Models that did not contain interactions were analyzed with a type II ANOVA. We subsequently plotted the fitted values of the simplest model against carbohydrate and protein levels. We also plotted thin-plate splines of the raw measurements against carbohydrate and protein levels. Finally, we plotted sexual size dimorphism (SSD) as the difference between the female and male fitted values against carbohydrate and protein level, as well a thin-plate splines of the difference between mean female and male values at each diet.

To test whether the response to carbohydrates or protein differed between the sexes we fit two models:

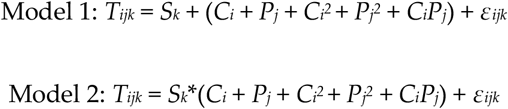

where *S* is sex. Models were compared using ANOVA (for standard models of expression level) or parametric boostrapping (for mixed models of body size). If inclusion of sex as an interactive factor rather than an additive factor significantly improved the fit of the model, we concluded that the relationship between trait and diet varied between males and females. If the relationship between trait and diet could be explained by a simpler model, we tested whether inclusion of sex as an interactive rather than an additive factor in the simplified model improved the fit of the model. We also conducted a similar multivariate analysis on the effect of diet and sex on the expression of multiple genes, using MANOVA to compare models. Finally, we used the fitted values from the relationship between diet and body size to test the relationship between body size and the expression of each gene, using the linear model:

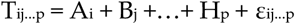

where *T* is body size (fitted values from the linear mixed model of the relationship between body size and diet) and *A* through *H* is the expression of the genes measured in our analysis.

All analyses were conducted in *R* and the data and scripts for the analyses are provided on Dryad. Gene expression levels were log-transformed prior to analysis. For all analyses, we plotted residual against fitted values to confirm homogeneity of variance and generated a QQ plot to confirm that the residuals were approximately normally distributed.

## Supporting information

R Markdown of Body Size Analysis

R Markdown of Gene Expression Analysis

## 7. Declarations

- Ethics approval and consent to participate: Not applicable
- Consent for publication: Not applicable
- Availability of data and materials: All the data and the R scripts used to analyse them will be made available on Dryad upon publication.
- Competing interests: The authors declare that they have no competing interests
- Funding: This research was funded by NSF grant IOS-1952385 awarded to AWS
- Authors’ contributions: JMCM, PN, LT and AWS conceived the study and its design; JMCM, PN, LT and SJ collected the data; JMCM, PN, LT, SJ and AWS analyzed the data and drafted the manuscript. All authors read and approved the final manuscript.
- Acknowledgements: The authors would like to thank the members of the Shingleton Lab for comments on early drafts of the manuscript.
- Authors’ information (optional): Not applicable

## 9. Supplementary Figures

**Supplementary Figure 1:**
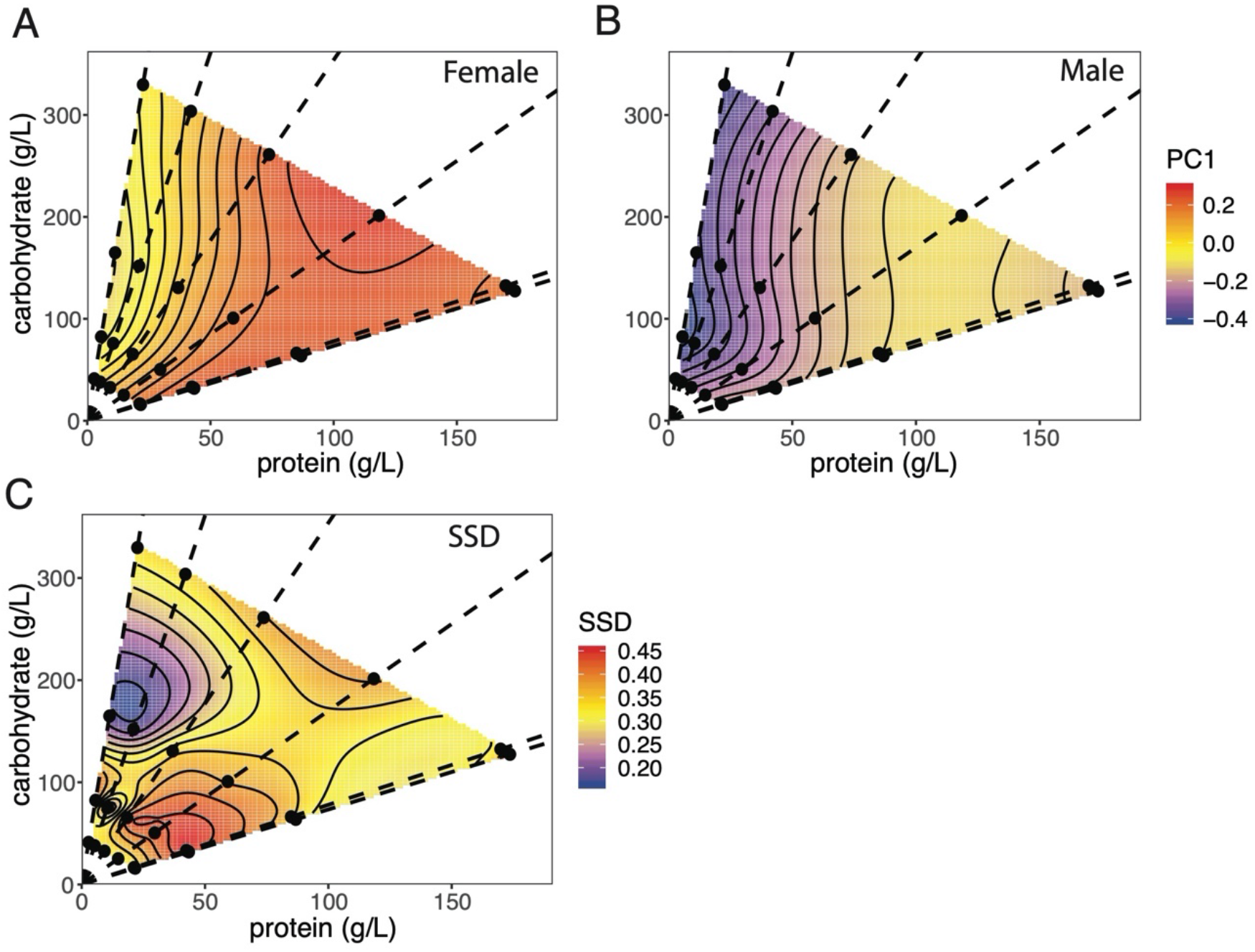
Thin plate spline of the effect of protein and carbohydrate concentration on female and male body size and sexual size dimorphism (SSD). (A, B) Surfaces shows the relationship between body size, carbohydrate level and protein level in female and male flies (C) Surface shows a thin plate spline of the difference in female and male body size (SSD) across the same nutritional landscape, using fitted values from A and B. Points indicate diets tested and dotted lines connect diets with equal protein-to-carbohydrate ratios (1:14.6, 1:7.2, 1:3.5, 1:1.7, 1.3:1, 1.4:1).

**Supplementary Figure 2:**
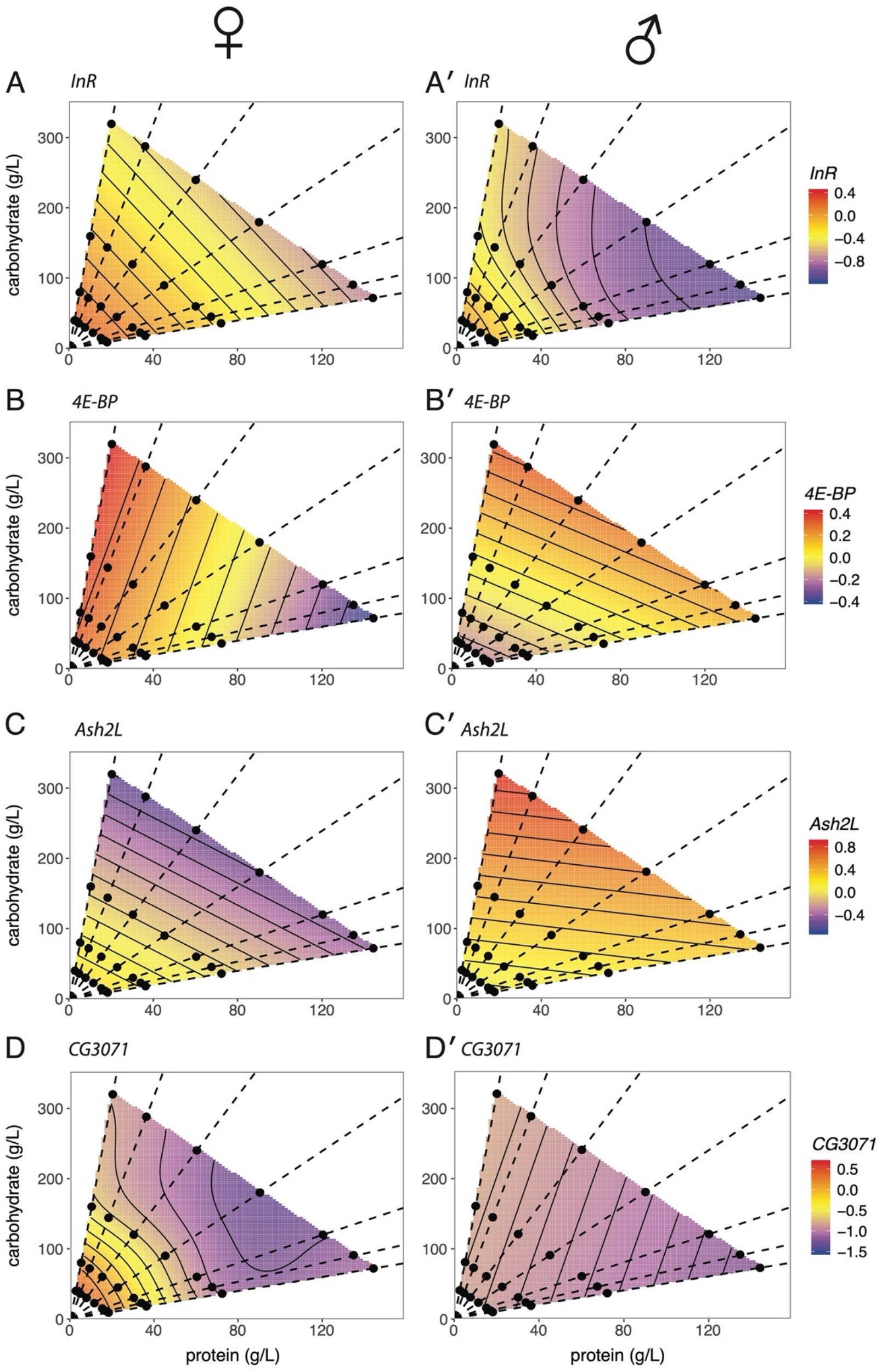
Thin plate spline of the effect of protein and carbohydrate concentration on the expression of IIS and TOR transcriptionally-regulated genes in females and males. Surfaces shows the relationship between gene expression, carbohydrate level and protein level in female and male flies. Expression of (A, A′) *InR*, and (B, B′) *4E-BP*, both negatively regulated by the activity of the IIS via the Forkhead transcription factor FOXO. (C, C′) Expression of *Ash2L*, ostensibly negatively regulated by the activity of TOR signaling. (D, D′) Expression of *CG3071*, ostensibly positively regulated by the activity of TOR signaling. Points indicate diets tested and dotted lines connect diets with equal protein-to-carbohydrate ratios (1:16, 1:8, 1:4, 1:2, 1:1, 2:1).

**Supplementary Figure 3:**
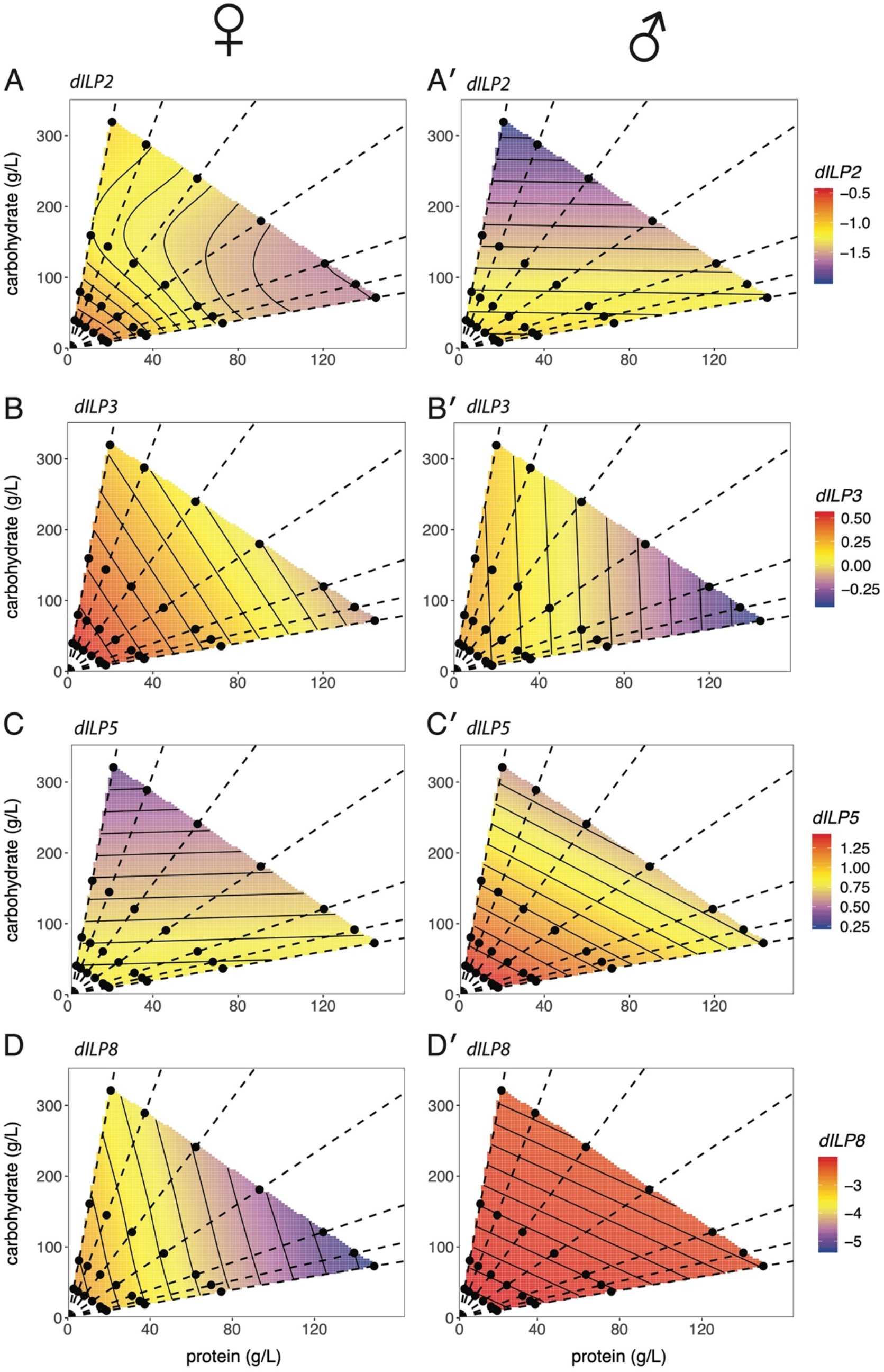
Thin plate spline of the effect of protein and carbohydrate concentration on the expression of dILPs in females and males. Surfaces shows the relationship between gene expression, carbohydrate level and protein level in female and male flies. (A) *dILP2*. (B) *dILP3.* (C) *dILP 5.* (D) *dILP8*. Points indicate diets tested and dotted lines connect diets with equal protein-to-carbohydrate ratios (1:16, 1:8, 1:4, 1:2, 1:1, 2:1). Corresponding thin-plate spline plots are shown in Supplementary Figure 1.

**Supplementary Table 1:**
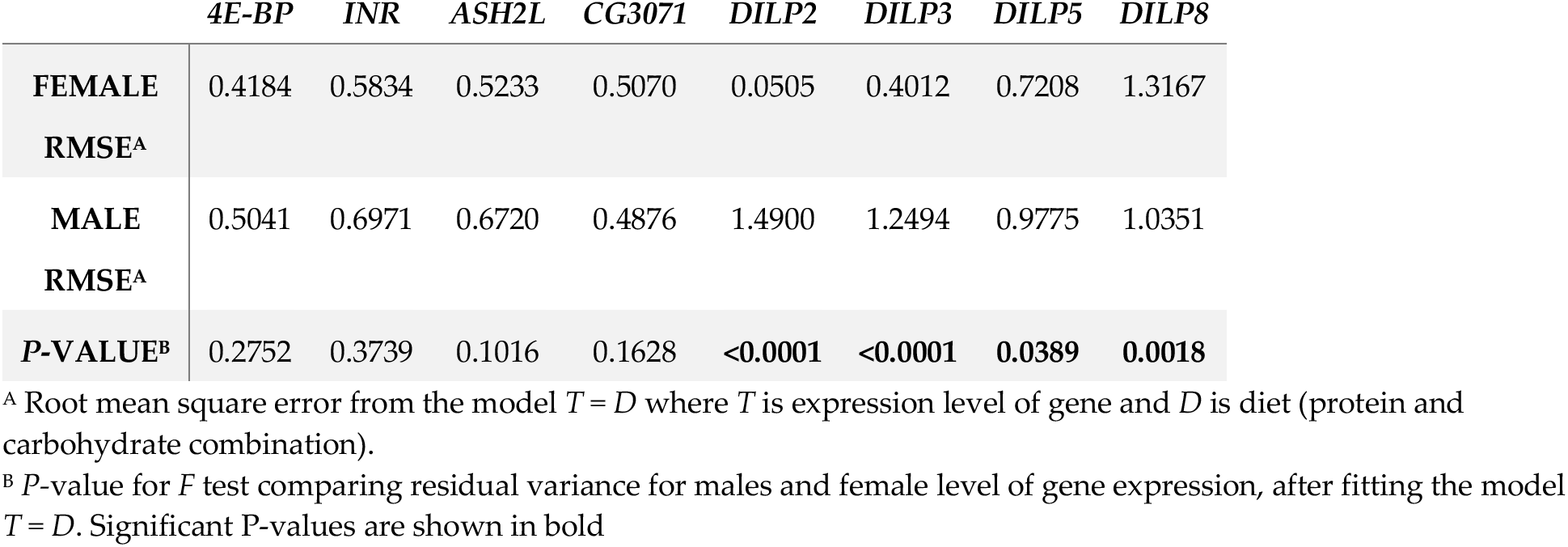
Variation in gene expression among samples within diets for male and female samples.

