## Supplementary material for "Sex-specific plasticity and the nutritional geometry of insulin-signaling gene expression in *Drosophila melanogaster*": R Markdown of Body Size Analysis

Alexander Shingleton

11/10/2020

### Analysis of the Nutritional Geometry of Gene Expression in Males and Females

Preamble

```
setwd("~/Documents/Data/Nutritional Geometry of Gene Expression/Final Analysis/Body Size")

suppressPackageStartupMessages({
    require("gdata")
    require("lmerTest")
    require("car")
    require("data.table")
    require("ggplot2")
    require("plyr")
    require("dplyr")
    require("lsmeans")
    require("mgcv")
    require("AICcmodavg")
    require("fields")
    require("lme4")
    require("piecewiseSEM")
    require("pbkrtest")
    require("car")
    require(MCMCglmm)  # To get posterior distributions
})

source("R/functions_body.R")
```

Import the data and add additional columns specifying nutritional details.

```
data <- read.csv("~/Documents/Data/Nutritional Geometry of Gene Expression/Final Analysis/Body Size/morp
diets <- read.csv("~/Documents/Data/Nutritional Geometry of Gene Expression/Final Analysis/Body Size/di

organsize <- within(data, {
    foodF <- factor(food)
    ratio <- factor(ratio, levels = paste("(", c("1:14.6", "1:7.2",
        "1:3.5", "1:1.7", "1.3:1", "1.4:1"), ")", sep = ""))
    inter = interaction(ratio, foodF)
    logprot = log(prot)
    logcarb = log(carb)
    logratio = log(ratio_num)
    logfood = log(food)
    repfull = paste(food, ratio, replicate, sep = "_")
})
```

```
organsize$cal <- data$food * 0.004
```

### Generate PCA1 from wing, thorax, leg and palp size

First we needvto calculate the value of the first principle component for each indvidual, as a measure of overall body size. To do this we need to eliminate all individuals where we do not have measurements of wing, thorax, leg and palp.

```
organsize <- organsize[!with(data, is.na(wing) | is.na(thorax) |
    is.na(leg) | is.na(palp)), ]

organsize$PCA1 <- prcomp(organsize[, c(11, 15, 16, 17)], center = T,
    scale. = F)$x[, 1]

write.csv(organsize, "organsize.csv")
```

#Female Analysis

Looking at the female data first. First we need to get rid of all missing values.

```
# for fem organ size
femsize <- (subset(organsize, sex == "F"))
femsize$genital <- NULL
femsize <- na.omit(femsize)
```

We will analyse the data using the model proposed by Lee et al., including replicate vials as a random factor in our analysis.

```
# The model used by lee et al., but with heterogeneity within
# replicates accounted for----
size.model.lee.replicates <- lmer(PCA1 ~ carb * prot + poly(carb,
    2) + poly(prot, 2) + (1 | repfull), data = femsize)
summary(size.model.lee.replicates)[10]
```

```
## $coefficients
##                       Estimate   Std. Error       df    t value     Pr(>|t|)
## (Intercept)     2.036105e-01 4.260685e-02 21.01247  4.7788217 1.010573e-04
## carb           -1.535366e-03 3.719818e-04 22.95910 -4.1275297 4.109142e-04
## prot            1.277453e-04 8.810182e-04 17.55215  0.1449974 8.863650e-01
## poly(carb, 2)2  1.202633e+00 2.596656e-01 21.97021  4.6314687 1.294529e-04
## poly(prot, 2)2 -1.548736e+00 2.572342e-01 17.82840 -6.0207246 1.124953e-05
## carb:prot       1.457952e-05 6.652392e-06 18.09549  2.1916204 4.172686e-02
```

```
Anova(size.model.lee.replicates, type = "III")
```

```
## Analysis of Deviance Table (Type III Wald chisquare tests)
##
## Response: PCA1
##                  Chisq Df Pr(>Chisq)
## (Intercept)    22.8371  1  1.763e-06 ***
## carb           17.0365  1  3.667e-05 ***
## prot            0.0210  1    0.88471
## poly(carb, 2) 21.4505  1  3.631e-06 ***
## poly(prot, 2) 36.2491  1  1.736e-09 ***
```

```
## carb:prot      4.8032  1    0.02841 *
## ---
## Signif. codes:  0 '***' 0.001 '**' 0.01 '*' 0.05 '.' 0.1 ' ' 1
```

```
# Get non-orthoganol paramters
summary(lmer(PCA1 ~ carb * prot + poly(carb, 2, raw = TRUE) +
    poly(prot, 2, raw = TRUE) + (1 | repfull), data = femsize))[10]
```

```
## $coefficients
##                                Estimate   Std. Error       df   t value
## (Intercept)                1.208015e-01 5.119537e-02 21.64087  2.359618
## carb                      -4.357400e-03 7.572064e-04 20.11517 -5.754574
## prot                       6.675500e-03 1.167252e-03 19.75656  5.718990
## poly(carb, 2, raw = TRUE)2  9.753910e-06 2.106008e-06 21.97021  4.631469
## poly(prot, 2, raw = TRUE)2 -3.467210e-05 5.758791e-06 17.82840 -6.020725
## carb:prot                  1.457952e-05 6.652392e-06 18.09549  2.191620
##                               Pr(>|t|)
## (Intercept)               2.774608e-02
## carb                      1.218702e-05
## prot                      1.412523e-05
## poly(carb, 2, raw = TRUE)2 1.294529e-04
## poly(prot, 2, raw = TRUE)2 1.124953e-05
## carb:prot                 4.172686e-02
```

All the parameters of the model are significant, although the carb:protein interaction is only marginally significiant.

We can now plot the model:

```
#+ Data preparation for plots, do not include in the output, include=F
# Generate Plot Space, Removing extrapolated points----
fembody.space <- genspace(femsize)
points <- genpoints(femsize)

## Generate predictions based on models.

fembody.predictions <- cbind(fembody.space, (predict(size.model.lee.replicates,
    fembody.space, re.form = NA)))
names(fembody.predictions)[6] <- "lee"

range(fembody.predictions$lee)
```

```
## [1] -0.2376796  0.2959486
```

```
body.range <- c(min(fembody.predictions$lee), max(fembody.predictions$lee))

#+Plots
### Level plots-----

fembody.plot.lee <- ggplot(data = fembody.predictions, aes(x = prot,
    y = carb, z = lee, fill = lee)) + geom_tile(alpha = 0.9) +
    theme_bw() + theme(panel.grid = element_blank(), axis.title.x = element_text(size = 16),
    axis.title.y = element_text(size = 16), axis.text.x = element_text(size = 14),
    axis.text.y = element_text(size = 14), legend.text = element_text(size = 14),
    legend.title = element_text(size = 16)) + scale_x_continuous(limits = c(0,
    max(diets$prot * 1.1)), expand = c(0, 0)) + scale_y_continuous(expand = c(0,
    0), limits = c(0, max(diets$carb * 1.1))) + geom_contour(color = "black",
```

```
    bins = 10) + xlab("protein (g/L)") + ylab("carbohydrate (g/L)") +
    scale_fill_gradientn(colours = c("blue", "yellow", "red"),
        name = "log body size") + geom_abline(data = ratios,
    aes(slope = 1/ratios, intercept = 0), linetype = 2, size = rel(0.8)) +
    geom_point(data = diets, aes(x = prot, y = carb, z = 1),
        size = 3, fill = "black") + guides(fill = guide_colorbar(order = 1))
fembody.plot.lee
```

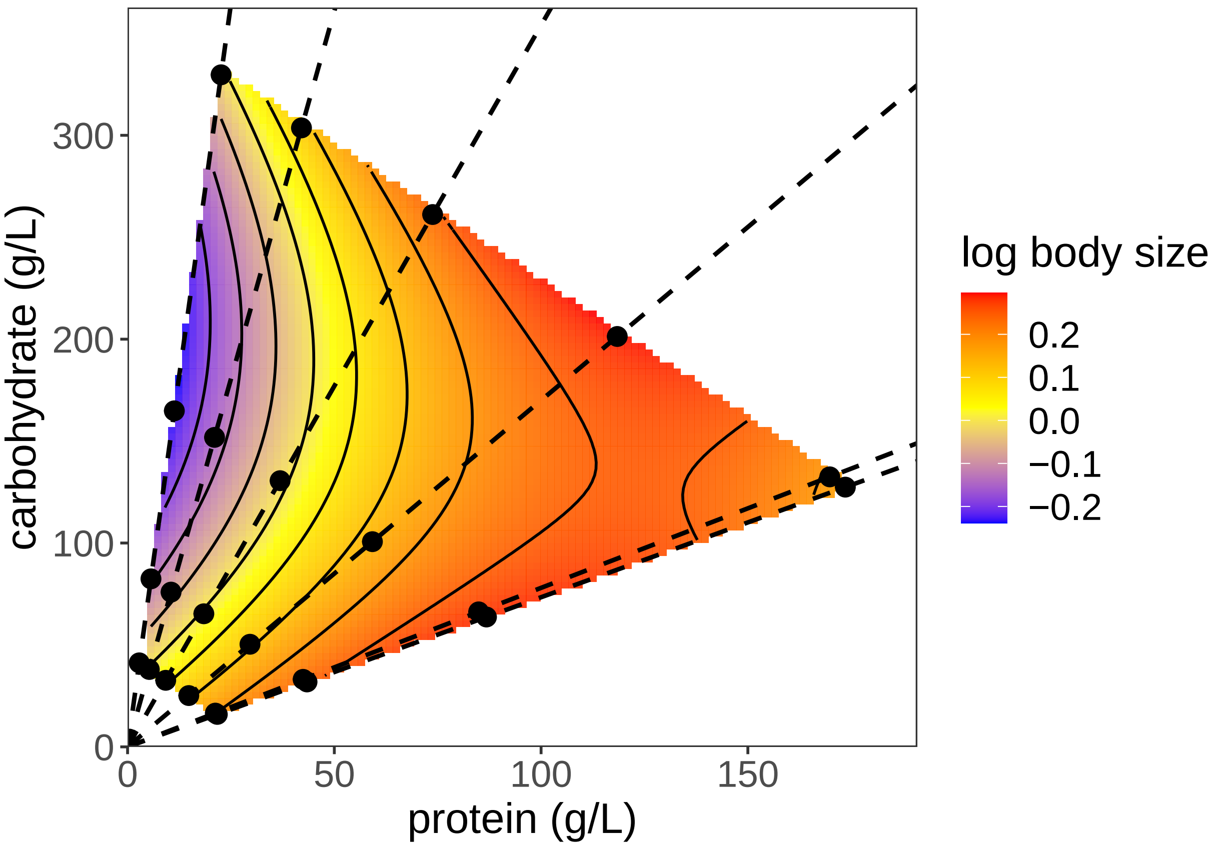


#Male Analysis

Now repeating for male body size

```
malesize <- (subset(organsize, sex == "M"))
malesize$genital <- NULL
malesize <- na.omit(malesize)

# The model used by lee et al., but with heterogeneity within
# replicates accounted for----
size.model.lee.replicates <- lmer(PCA1 ~ carb * prot + poly(carb,
    2) + poly(prot, 2) + (1 | repfull), data = malesize)

anova(size.model.lee.replicates)
```

```
## Type III Analysis of Variance Table with Satterthwaite's method
##                  Sum Sq  Mean Sq NumDF  DenDF F value   Pr(>F)
## carb           0.030274 0.030274     1 25.314  2.3477 0.137870
## prot           0.000439 0.000439     1 21.881  0.0340 0.855338
## poly(carb, 2) 0.007004 0.007004     1 23.926  0.5431 0.468301
```

```
## poly(prot, 2) 0.129617 0.129617     1 22.640 10.0518 0.004325 **
## carb:prot     0.014879 0.014879     1 22.406  1.1539 0.294172
## ---
## Signif. codes:  0 '***' 0.001 '**' 0.01 '*' 0.05 '.' 0.1 ' ' 1
```

```
summary(size.model.lee.replicates)
```

```
## Linear mixed model fit by REML. t-tests use Satterthwaite's method [
## lmerModLmerTest]
## Formula: PCA1 ~ carb * prot + poly(carb, 2) + poly(prot, 2) + (1 | repfull)
##    Data: malesize
##
## REML criterion at convergence: -387.6
##
## Scaled residuals:
##     Min      1Q  Median      3Q     Max
## -3.7138 -0.5329 -0.0198  0.5511  3.1962
##
## Random effects:
##  Groups   Name        Variance Std.Dev.
##  repfull  (Intercept) 0.01067  0.1033
##  Residual             0.01289  0.1136
## Number of obs: 347, groups:  repfull, 55
##
## Fixed effects:
##                  Estimate Std. Error         df t value Pr(>|t|)
## (Intercept)    -2.318e-01  6.296e-02  2.393e+01  -3.682  0.00118 **
## carb           -8.331e-04  5.437e-04  2.531e+01  -1.532  0.13787
## prot            2.367e-04  1.283e-03  2.188e+01   0.184  0.85534
## poly(carb, 2)2  2.623e-01  3.560e-01  2.393e+01   0.737  0.46830
## poly(prot, 2)2 -1.041e+00  3.284e-01  2.264e+01  -3.170  0.00432 **
## carb:prot       1.035e-05  9.637e-06  2.241e+01   1.074  0.29417
## ---
## Signif. codes:  0 '***' 0.001 '**' 0.01 '*' 0.05 '.' 0.1 ' ' 1
##
## Correlation of Fixed Effects:
##              (Intr) carb   prot   ply(c,2)2 ply(p,2)2
## carb         -0.882
## prot         -0.891  0.813
## ply(crb,2)2 -0.261  0.103  0.268
## ply(prt,2)2 -0.424  0.250  0.319  0.392
## carb:prot    0.850 -0.901 -0.946 -0.148    -0.238
## fit warnings:
## fixed-effect model matrix is rank deficient so dropping 2 columns / coefficients
## Some predictor variables are on very different scales: consider rescaling
```

Because only protein has an effect on body size, we can eliminate the effect of carbohydrates and re-test.

```
# Revised model used by lee et al., but with heterogeneity
# within replicates accounted for----
size.model.lee.replicates <- lmer(PCA1 ~ poly(prot, 2) + (1 |
    repfull), data = malesize)
summary(size.model.lee.replicates)[10]
```

```
## $coefficients
##                  Estimate Std. Error       df     t value     Pr(>|t|)
```

```
## (Intercept)    -0.2231327 0.01791467 27.34065 -12.455308 8.759641e-13
## poly(prot, 2)1  1.0696301 0.31668091 26.98528   3.377627 2.235273e-03
## poly(prot, 2)2 -1.0203677 0.30071978 30.75433  -3.393085 1.918506e-03
```

```
Anova(size.model.lee.replicates, type = "III")
```

```
## Analysis of Deviance Table (Type III Wald chisquare tests)
##
## Response: PCA1
##                  Chisq Df Pr(>Chisq)
## (Intercept)    155.135  1  < 2.2e-16 ***
## poly(prot, 2)   29.519  2   3.89e-07 ***
## ---
## Signif. codes:  0 '***' 0.001 '**' 0.01 '*' 0.05 '.' 0.1 ' ' 1
```

```
# Get non-orthoganol paramters
summary(lmer(PCA1 ~ poly(prot, 2, raw = TRUE) + (1 | repfull),
    data = malesize))[10]
```

```
## $coefficients
##                                 Estimate   Std. Error       df    t value
## (Intercept)                -4.454493e-01 4.036976e-02 43.05873 -11.034232
## poly(prot, 2, raw = TRUE)1  5.893507e-03 1.332743e-03 33.49204   4.422089
## poly(prot, 2, raw = TRUE)2 -2.509812e-05 7.396844e-06 30.75433  -3.393085
##                                Pr(>|t|)
## (Intercept)                3.932563e-14
## poly(prot, 2, raw = TRUE)1 9.760184e-05
## poly(prot, 2, raw = TRUE)2 1.918506e-03
```

We can now plot the model:

```
# Plot Model

#+ Data preparation for plots, do not include in the output, include=F
# Generate Plot Space that is the same as for females
malebody.space <- genspace(femsize)
points <- genpoints(femsize)

## Generate predictions based on models.


malebody.predictions <- cbind(malebody.space, (predict(size.model.lee.replicates,
    malebody.space, re.form = NA)))
names(malebody.predictions)[6] <- "lee"

range(malebody.predictions$lee)
```

```
## [1] -0.41293387 -0.09947388
```

```
body.range <- c(min(malebody.predictions$lee), max(malebody.predictions$lee))
body.range
```

```
## [1] -0.41293387 -0.09947388
```

```
#+Plots
### Level plots-----

malebody.plot.lee <- ggplot(data = malebody.predictions, aes(x = prot,
```

```
    y = carb, z = lee, fill = lee)) + geom_tile(alpha = 0.9) +
    theme_bw() + theme(panel.grid = element_blank(), axis.title.x = element_text(size = 16),
    axis.title.y = element_text(size = 16), axis.text.x = element_text(size = 14),
    axis.text.y = element_text(size = 14), legend.text = element_text(size = 14),
    legend.title = element_text(size = 16)) + scale_x_continuous(limits = c(0,
    max(diets$prot * 1.1)), expand = c(0, 0)) + scale_y_continuous(expand = c(0,
    0), limits = c(0, max(diets$carb * 1.1))) + geom_contour(color = "black",
    bins = 10) + xlab("protein (g/L)") + ylab("carbohydrate (g/L)") +
    scale_fill_gradientn(colours = c("blue", "yellow", "red"),
        name = "log body size") + geom_abline(data = ratios,
    aes(slope = 1/ratios, intercept = 0), linetype = 2, size = rel(0.8)) +
    geom_point(data = diets, aes(x = prot, y = carb, z = 1),
        size = 3, fill = "black") + guides(fill = guide_colorbar(order = 1))
malebody.plot.lee
```

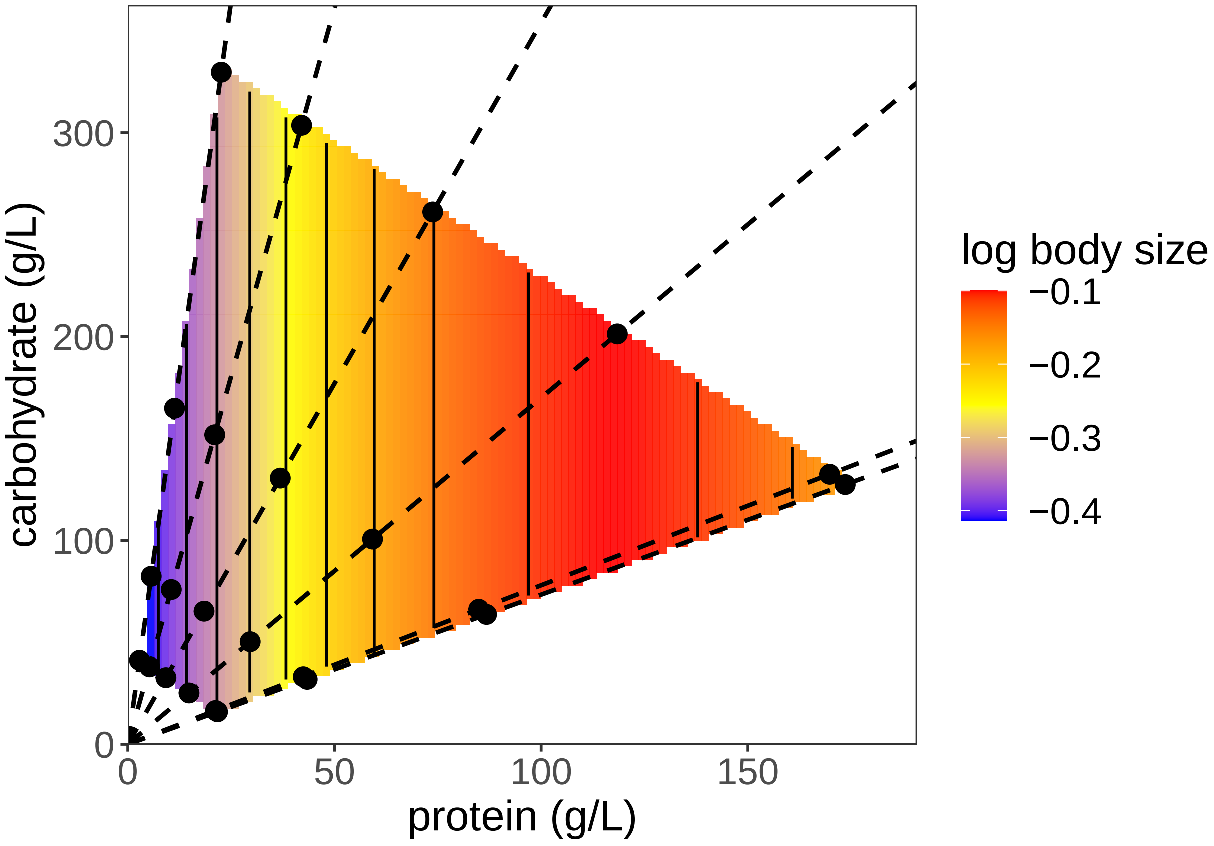


It is useful to replot the male and female data on the same scale, to allow for direct comparison:

```
# Replot using same scale for males and females

body.range <- c(min(malebody.predictions$lee), max(fembody.predictions$lee))

fembody.plot.lee <- ggplot(data = fembody.predictions, aes(x = prot,
    y = carb, z = lee, fill = lee)) + geom_tile(alpha = 0.9) +
    theme_bw() + theme(panel.grid = element_blank(), axis.title.x = element_text(size = 16),
    axis.title.y = element_text(size = 16), axis.text.x = element_text(size = 14),
    axis.text.y = element_text(size = 14), legend.text = element_text(size = 14),
    legend.title = element_text(size = 16)) + scale_x_continuous(limits = c(0,
```

```
    max(diets$prot * 1.1)), expand = c(0, 0)) + scale_y_continuous(expand = c(0,
    0), limits = c(0, max(diets$carb * 1.1))) + geom_contour(color = "black",
    bins = 10) + xlab("protein (g/L)") + ylab("carbohydrate (g/L)") +
    scale_fill_gradientn(colours = c("blue", "yellow", "red"),
        name = "PC1", limits = body.range) + geom_abline(data = ratios,
    aes(slope = 1/ratios, intercept = 0), linetype = 2, size = rel(0.8)) +
    geom_point(data = diets, aes(x = prot, y = carb, z = 1),
        size = 3, fill = "black") + guides(fill = guide_colorbar(order = 1))
fembody.plot.lee
```

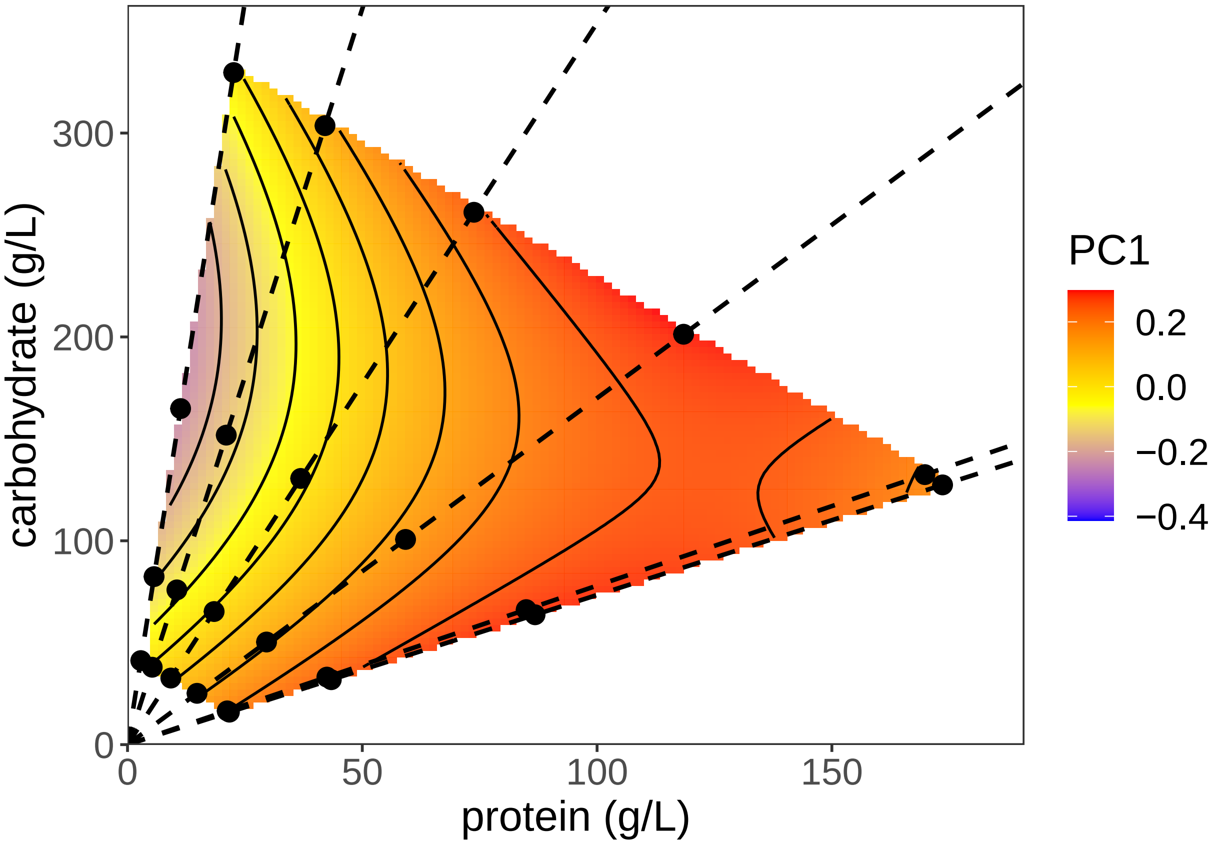


```
malebody.plot.lee <- ggplot(data = malebody.predictions, aes(x = prot,
    y = carb, z = lee, fill = lee)) + geom_tile(alpha = 0.9) +
    theme_bw() + theme(panel.grid = element_blank(), axis.title.x = element_text(size = 16),
    axis.title.y = element_text(size = 16), axis.text.x = element_text(size = 14),
    axis.text.y = element_text(size = 14), legend.text = element_text(size = 14),
    legend.title = element_text(size = 16)) + scale_x_continuous(limits = c(0,
    max(diets$prot * 1.1)), expand = c(0, 0)) + scale_y_continuous(expand = c(0,
    0), limits = c(0, max(diets$carb * 1.1))) + geom_contour(color = "black",
    bins = 10) + xlab("protein (g/L)") + ylab("carbohydrate (g/L)") +
    scale_fill_gradientn(colours = c("blue", "yellow", "red"),
        name = "PC1", limits = body.range) + geom_abline(data = ratios,
    aes(slope = 1/ratios, intercept = 0), linetype = 2, size = rel(0.8)) +
    geom_point(data = diets, aes(x = prot, y = carb, z = 1),
        size = 3, fill = "black") + guides(fill = guide_colorbar(order = 1))
malebody.plot.lee
```

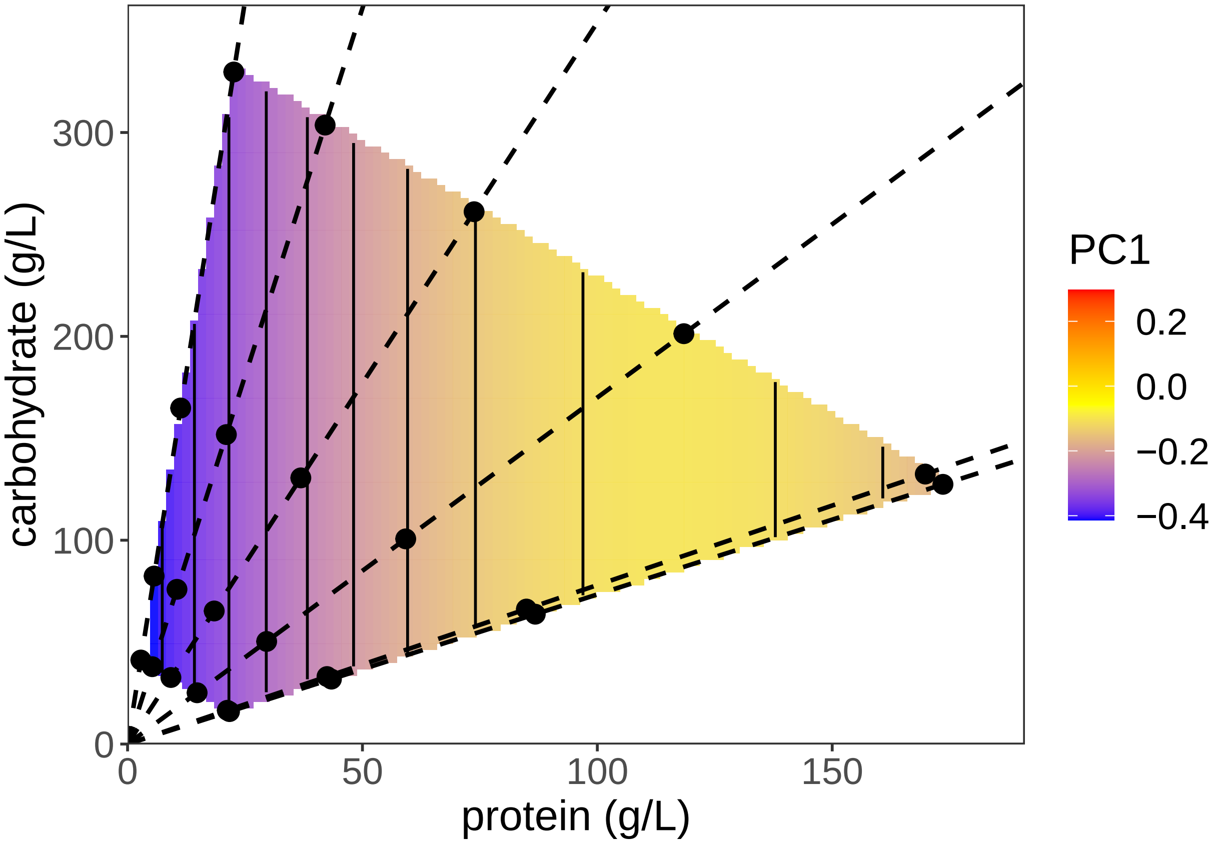


For completeness, we can also plot the thin plate splines. Females first, using the same scale as for the model.

```
tps.f <- with(femsize, Tps(cbind(prot, carb), PCA1))

gen.tps.f <- within(fembody.space, {
    tps <- predict(tps.f, data.frame(prot, carb))
})

gen.plot.tps.f <- ggplot(data = gen.tps.f, aes(x = prot, y = carb,
    z = tps, fill = tps)) + geom_tile(alpha = 0.9) + theme_bw() +
    theme(panel.grid = element_blank(), axis.title.x = element_text(size = 16),
        axis.title.y = element_text(size = 16), axis.text.x = element_text(size = 14),
        axis.text.y = element_text(size = 14), legend.text = element_text(size = 14),
        legend.title = element_text(size = 16)) + scale_x_continuous(limits = c(0,
    max(diets$prot * 1.1)), expand = c(0, 0)) + scale_y_continuous(expand = c(0,
    0), limits = c(0, max(diets$carb * 1.1))) + geom_contour(color = "black",
    bins = 10) + xlab("protein (g/L)") + ylab("carbohydrate (g/L)") +
    scale_fill_gradientn(colours = c("blue", "yellow", "red"),
        name = "PC1", limits = body.range) + geom_abline(data = ratios,
    aes(slope = 1/ratios, intercept = 0), linetype = 2, size = rel(0.8)) +
    geom_point(data = diets, aes(x = prot, y = carb, z = 1),
        size = 3, fill = "black") + guides(fill = guide_colorbar(order = 1)) +
    ggtitle("female.tps")
gen.plot.tps.f
```

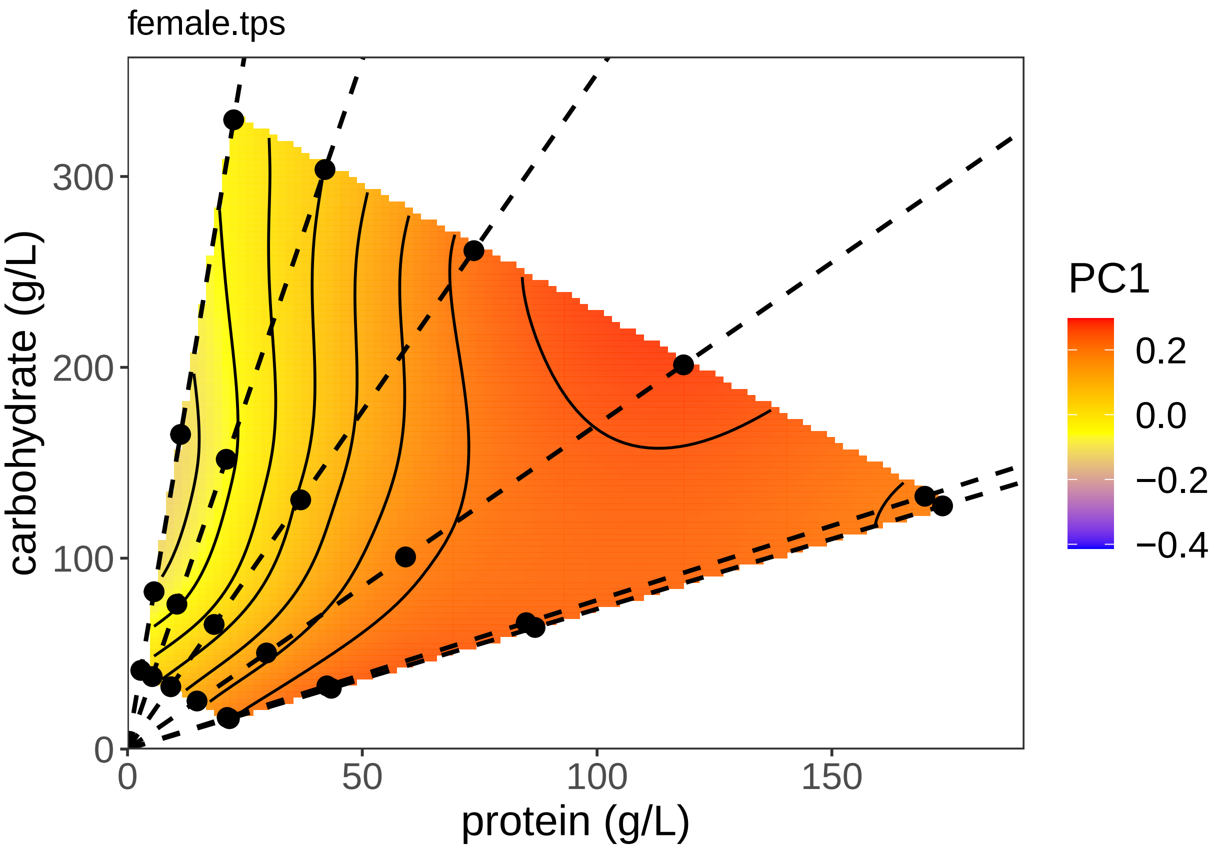


Now repeating for the males:

```
tps.m <- with(malesize, Tps(cbind(prot, carb), PCA1))

gen.tps.m <- within(malebody.space, {
    tps <- predict(tps.m, data.frame(prot, carb))
})

gen.plot.tps.m <- ggplot(data = gen.tps.m, aes(x = prot, y = carb,
    z = tps, fill = tps)) + geom_tile(alpha = 0.9) + theme_bw() +
    theme(panel.grid = element_blank(), axis.title.x = element_text(size = 16),
        axis.title.y = element_text(size = 16), axis.text.x = element_text(size = 14),
        axis.text.y = element_text(size = 14), legend.text = element_text(size = 14),
        legend.title = element_text(size = 16)) + scale_x_continuous(limits = c(0,
    max(diets$prot * 1.1)), expand = c(0, 0)) + scale_y_continuous(expand = c(0,
    0), limits = c(0, max(diets$carb * 1.1))) + geom_contour(color = "black",
    bins = 10) + xlab("protein (g/L)") + ylab("carbohydrate (g/L)") +
    scale_fill_gradientn(colours = c("blue", "yellow", "red"),
        name = "PC1", limits = body.range) + geom_abline(data = ratios,
    aes(slope = 1/ratios, intercept = 0), linetype = 2, size = rel(0.8)) +
    geom_point(data = diets, aes(x = prot, y = carb, z = 1),
        size = 3, fill = "black") + guides(fill = guide_colorbar(order = 1)) +
    ggtitle("male.tps")
gen.plot.tps.m
```

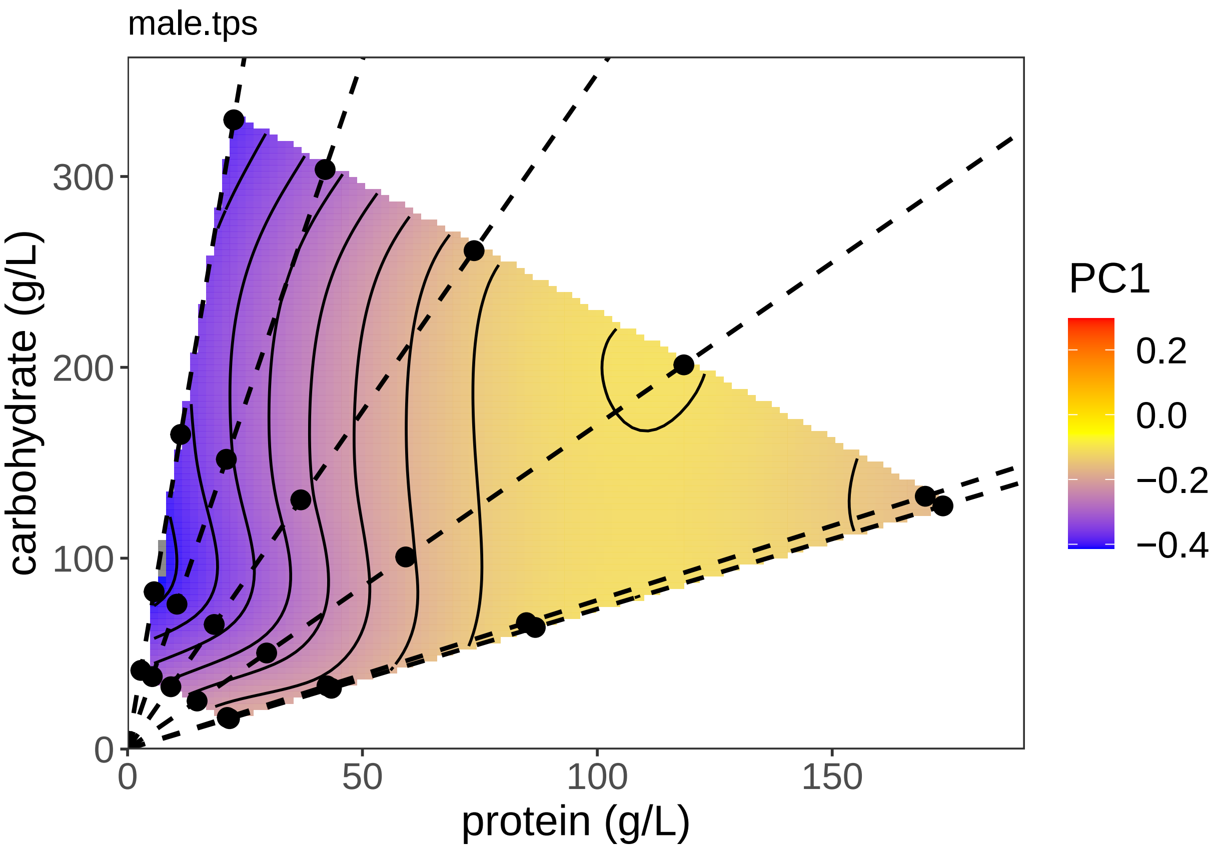


#Plotting the SSD

It is interesting to see how SSD varies across the nutritional landscape. We can use the fitted values from the above analysis to do this, using the difference in (fitted) male and female body size as a measure of SSD.

```
# Subtract Male From Female
all.predictions <- malebody.predictions %>% right_join(fembody.predictions,
    by = c("carb", "prot"))
all.predictions$SSD <- all.predictions$lee.y - all.predictions$lee.x

SSD.plot.lee <- ggplot(data = all.predictions, aes(x = prot,
    y = carb, z = SSD, fill = SSD)) + geom_tile(alpha = 0.9) +
    theme_bw() + theme(panel.grid = element_blank(), axis.title.x = element_text(size = 16),
    axis.title.y = element_text(size = 16), axis.text.x = element_text(size = 14),
    axis.text.y = element_text(size = 14), legend.text = element_text(size = 14),
    legend.title = element_text(size = 16)) + scale_x_continuous(limits = c(0,
    max(diets$prot * 1.1)), expand = c(0, 0)) + scale_y_continuous(expand = c(0,
    0), limits = c(0, max(diets$carb * 1.1))) + geom_contour(color = "black",
    bins = 10) + xlab("protein (g/L)") + ylab("carbohydrate (g/L)") +
    scale_fill_gradientn(colours = c("blue", "yellow", "red"),
        name = "SSD") + geom_abline(data = ratios, aes(slope = 1/ratios,
    intercept = 0), linetype = 2, size = rel(0.8)) + geom_point(data = diets,
    aes(x = prot, y = carb, z = 1), size = 3, fill = "black") +
    guides(fill = guide_colorbar(order = 1))

SSD.plot.lee
```

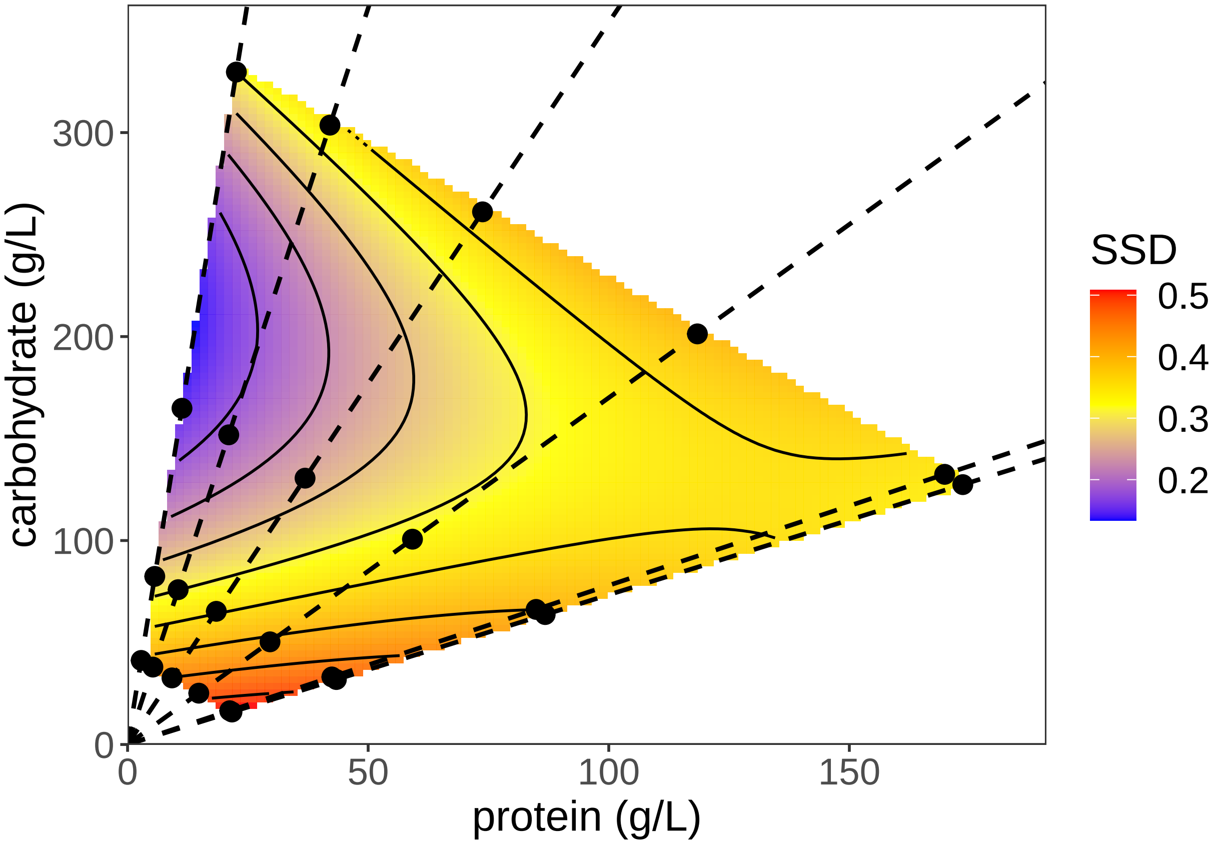


We can also plot mean SSD at each diet.

```
mean_m <- aggregate(PCA1 ~ prot + carb, data = malesize, FUN = "mean")
mean_f <- aggregate(PCA1 ~ prot + carb, data = femsize, FUN = "mean")
all.means <- mean_m %>% right_join(mean_f, by = c("carb", "prot"))
all.means$SSD <- all.means$PCA1.y - all.means$PCA1.x

tps.SSD <- with(all.means, Tps(cbind(prot, carb), SSD))
```

```
## Warning:
## Grid searches over lambda (nugget and sill variances) with  minima at the endpoints:
##   (GCV) Generalized Cross-Validation
##    minimum at  right endpoint  lambda  =  2.286841e-06 (eff. df= 17.10001 )
```

```
gen.tps.SSD <- within(fembody.space, {
    tps <- predict(tps.SSD, data.frame(prot, carb))
})

SSD.range <- c(min(gen.tps.SSD$tps), max(gen.tps.SSD$tps))

gen.plot.tps.SSD <- ggplot(data = gen.tps.SSD, aes(x = prot,
    y = carb, z = tps, fill = tps)) + geom_tile(alpha = 0.9) +
    theme_bw() + theme(panel.grid = element_blank(), axis.title.x = element_text(size = 16),
    axis.title.y = element_text(size = 16), axis.text.x = element_text(size = 14),
    axis.text.y = element_text(size = 14), legend.text = element_text(size = 14),
    legend.title = element_text(size = 16)) + scale_x_continuous(limits = c(0,
    max(diets$prot * 1.1)), expand = c(0, 0)) + scale_y_continuous(expand = c(0,
    0), limits = c(0, max(diets$carb * 1.1))) + geom_contour(color = "black",
```

```
    bins = 10) + xlab("protein (g/L)") + ylab("carbohydrate (g/L)") +
    scale_fill_gradientn(colours = c("blue", "yellow", "red"),
        name = "SSD", limits = SSD.range) + geom_abline(data = ratios,
    aes(slope = 1/ratios, intercept = 0), linetype = 2, size = rel(0.8)) +
    geom_point(data = diets, aes(x = prot, y = carb, z = 1),
        size = 3, fill = "black") + guides(fill = guide_colorbar(order = 1)) +
    ggtitle("SSD.tps")
gen.plot.tps.SSD
```

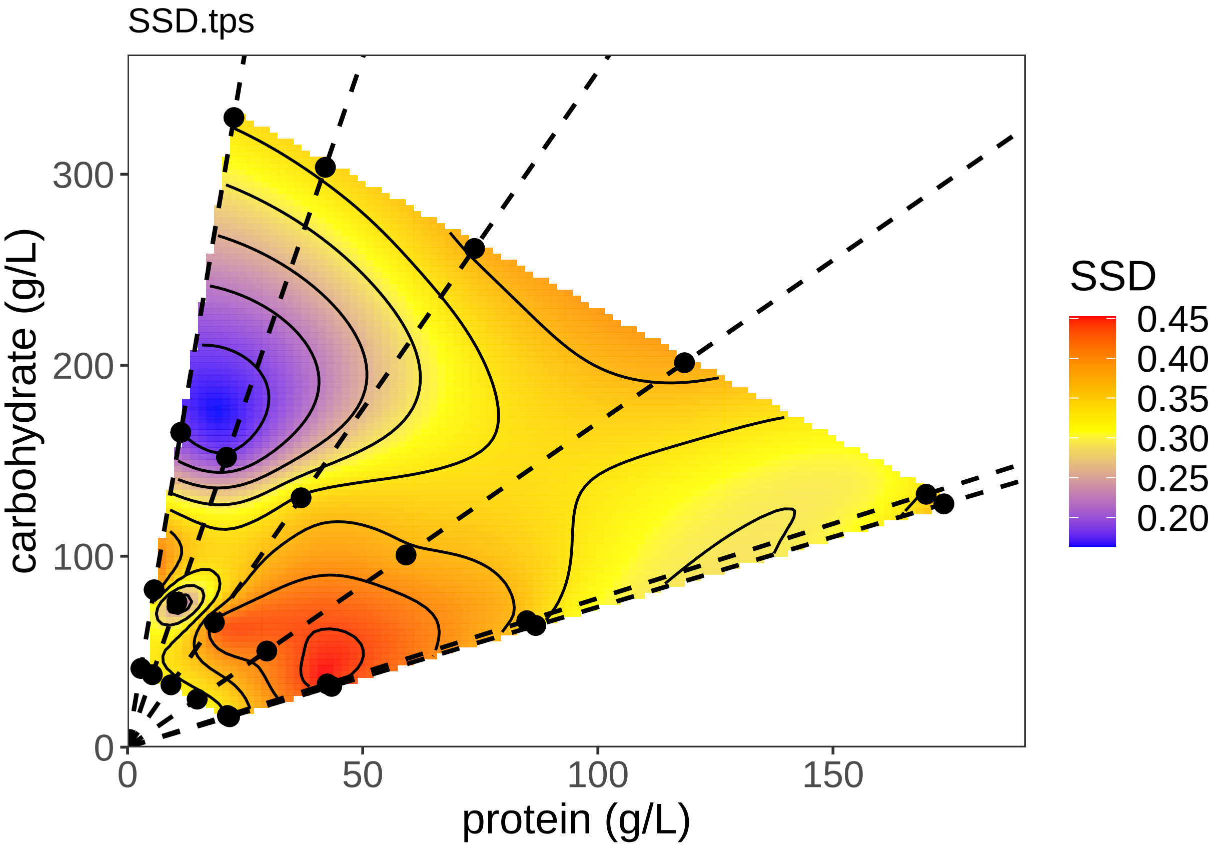


It looks like SSD varies with diet, suggesting that males and females differ in their response to diet. We can formally test this using lmer:

```
sex.test.0 <- lmer(PCA1 ~ sex + carb:prot + poly(carb, 2) + poly(prot,
    2) + (1 | repfull), data = organsize, REML = F)
summary(sex.test.0)
```

```
## Linear mixed model fit by maximum likelihood . t-tests use Satterthwaite's
##   method [lmerModLmerTest]
## Formula: PCA1 ~ sex + carb:prot + poly(carb, 2) + poly(prot, 2) + (1 |
##     repfull)
##    Data: organsize
##
##      AIC      BIC   logLik deviance df.resid
##  -1059.0  -1017.3    538.5  -1077.0      748
##
## Scaled residuals:
##     Min      1Q  Median      3Q     Max
```

```
## -6.7795 -0.5678  0.0476  0.5815  3.3174
##
## Random effects:
##  Groups   Name        Variance Std.Dev.
##  repfull  (Intercept) 0.003787 0.06154
##  Residual             0.012683 0.11262
## Number of obs: 757, groups:  repfull, 61
##
## Fixed effects:
##                   Estimate Std. Error         df t value Pr(>|t|)
## (Intercept)      7.330e-02  4.464e-02  3.072e+01   1.642 0.110754
## sexM            -3.783e-01  8.611e-03  7.199e+02 -43.929  < 2e-16 ***
## poly(carb, 2)1 -2.676e+00  6.562e-01  3.887e+01  -4.078 0.000217 ***
## poly(carb, 2)2  1.113e+00  2.926e-01  3.559e+01   3.805 0.000537 ***
## poly(prot, 2)1  2.135e-01  9.383e-01  2.923e+01   0.228 0.821570
## poly(prot, 2)2 -1.800e+00  2.849e-01  2.971e+01  -6.320 5.97e-07 ***
## carb:prot       1.244e-05  5.497e-06  3.051e+01   2.263 0.030920 *
## ---
## Signif. codes:  0 '***' 0.001 '**' 0.01 '*' 0.05 '.' 0.1 ' ' 1
##
## Correlation of Fixed Effects:
##              (Intr) sexM   ply(c,2)1 ply(c,2)2 ply(p,2)1 ply(p,2)2
## sexM         -0.089
## ply(crb,2)1  0.860 -0.016
## ply(crb,2)2  0.033  0.033 -0.040
## ply(prt,2)1  0.921  0.005  0.784     0.193
## ply(prt,2)2  0.154 -0.006  0.224     0.313     0.315
## carb:prot   -0.968  0.005 -0.892    -0.039    -0.936    -0.219
## fit warnings:
## Some predictor variables are on very different scales: consider rescaling
```

```r
sex.test.1 <- lmer(PCA1 ~ carb:prot + sex * poly(prot, 2) + poly(carb,
    2) + (1 | repfull), data = organsize, REML = F)
summary(sex.test.1)
```

```
## Linear mixed model fit by maximum likelihood . t-tests use Satterthwaite's
##   method [lmerModLmerTest]
## Formula: PCA1 ~ carb:prot + sex * poly(prot, 2) + poly(carb, 2) + (1 |
##     repfull)
##    Data: organsize
##
##      AIC      BIC   logLik deviance df.resid
##  -1057.7  -1006.8    539.8  -1079.7      746
##
## Scaled residuals:
##     Min      1Q  Median      3Q     Max
## -6.7548 -0.5674  0.0524  0.5758  3.2844
##
## Random effects:
##  Groups   Name        Variance Std.Dev.
##  repfull  (Intercept) 0.003904 0.06248
##  Residual             0.012613 0.11231
## Number of obs: 757, groups:  repfull, 61
##
## Fixed effects:
```

```
##                        Estimate Std. Error         df t value Pr(>|t|)
## (Intercept)          7.349e-02  4.513e-02  3.081e+01   1.628 0.113613
## sexM                -3.784e-01  8.598e-03  7.217e+02 -44.013  < 2e-16 ***
## poly(prot, 2)1       1.596e-01  9.538e-01  3.001e+01   0.167 0.868193
## poly(prot, 2)2      -1.952e+00  3.068e-01  3.849e+01  -6.362 1.72e-07 ***
## poly(carb, 2)1      -2.676e+00  6.625e-01  3.892e+01  -4.039 0.000244 ***
## poly(carb, 2)2       1.126e+00  2.958e-01  3.553e+01   3.805 0.000538 ***
## carb:prot            1.242e-05  5.558e-06  3.061e+01   2.234 0.032918 *
## sexM:poly(prot, 2)1  1.666e-01  2.326e-01  7.109e+02   0.716 0.474023
## sexM:poly(prot, 2)2  3.453e-01  2.345e-01  7.340e+02   1.473 0.141289
## ---
## Signif. codes:  0 '***' 0.001 '**' 0.01 '*' 0.05 '.' 0.1 ' ' 1
##
## Correlation of Fixed Effects:
##             (Intr) sexM   ply(p,2)1 ply(p,2)2 ply(c,2)1 ply(c,2)2 crb:pr
## sexM        -0.088
## ply(prt,2)1  0.914  0.009
## ply(prt,2)2  0.149 -0.010  0.291
## ply(crb,2)1  0.860 -0.017  0.777     0.216
## ply(crb,2)2  0.032  0.034  0.194     0.282    -0.042
## carb:prot   -0.968  0.006 -0.930    -0.210    -0.891    -0.037
## sxM:pl(,2)1  0.015 -0.038 -0.102     0.026     0.018    -0.030    -0.011
## sxM:pl(,2)2 -0.014  0.011  0.005    -0.344    -0.019     0.035     0.012
##             sM:(,2)1
## sexM
## ply(prt,2)1
## ply(prt,2)2
## ply(crb,2)1
## ply(crb,2)2
## carb:prot
## sxM:pl(,2)1
## sxM:pl(,2)2 -0.029
## fit warnings:
## Some predictor variables are on very different scales: consider rescaling
```

```r
sex.test.2 <- lmer(PCA1 ~ carb:prot + sex * poly(carb, 2) + poly(prot,
    2) + (1 | repfull), data = organsize, REML = F)
summary(sex.test.2)
```

```
## Linear mixed model fit by maximum likelihood . t-tests use Satterthwaite's
##   method [lmerModLmerTest]
## Formula: PCA1 ~ carb:prot + sex * poly(carb, 2) + poly(prot, 2) + (1 |
##     repfull)
##    Data: organsize
##
##      AIC      BIC   logLik deviance df.resid
##  -1066.7  -1015.8    544.3  -1088.7      746
##
## Scaled residuals:
##     Min      1Q  Median      3Q     Max
## -6.5824 -0.5609  0.0417  0.5944  3.6066
##
## Random effects:
##  Groups   Name        Variance Std.Dev.
##  repfull  (Intercept) 0.003882 0.06231
```

```
##  Residual             0.012460 0.11162
## Number of obs: 757, groups:  repfull, 61
##
## Fixed effects:
##                       Estimate Std. Error         df t value Pr(>|t|)
## (Intercept)          6.940e-02  4.499e-02  3.104e+01   1.543   0.1331
## sexM                -3.800e-01  8.554e-03  7.223e+02 -44.427  < 2e-16 ***
## poly(carb, 2)1      -2.931e+00  6.681e-01  4.170e+01  -4.387 7.66e-05 ***
## poly(carb, 2)2       1.438e+00  3.130e-01  4.781e+01   4.593 3.20e-05 ***
## poly(prot, 2)1       1.656e-01  9.459e-01  2.955e+01   0.175   0.8622
## poly(prot, 2)2      -1.808e+00  2.871e-01  2.999e+01  -6.297 6.11e-07 ***
## carb:prot            1.295e-05  5.542e-06  3.085e+01   2.337   0.0261 *
## sexM:poly(carb, 2)1  4.184e-01  2.372e-01  7.262e+02   1.764   0.0782 .
## sexM:poly(carb, 2)2 -6.966e-01  2.371e-01  7.373e+02  -2.938   0.0034 **
## ---
## Signif. codes:  0 '***' 0.001 '**' 0.01 '*' 0.05 '.' 0.1 ' ' 1
##
## Correlation of Fixed Effects:
##             (Intr) sexM   ply(c,2)1 ply(c,2)2 ply(p,2)1 ply(p,2)2 crb:pr
## sexM        -0.085
## ply(crb,2)1  0.850 -0.009
## ply(crb,2)2  0.019  0.013 -0.055
## ply(prt,2)1  0.921  0.006  0.774     0.172
## ply(prt,2)2  0.154 -0.006  0.219     0.290     0.316
## carb:prot   -0.968  0.003 -0.881    -0.023    -0.936    -0.220
## sxM:pl(,2)1  0.002 -0.029 -0.151     0.012     0.006     0.012    -0.003
## sxM:pl(,2)2  0.036  0.052  0.044    -0.339     0.026     0.016    -0.038
##             sM:(,2)1
## sexM
## ply(crb,2)1
## ply(crb,2)2
## ply(prt,2)1
## ply(prt,2)2
## carb:prot
## sxM:pl(,2)1
## sxM:pl(,2)2  0.006
## fit warnings:
## Some predictor variables are on very different scales: consider rescaling
```

```r
sex.test.3 <- lmer(PCA1 ~ sex * (carb:prot + poly(carb, 2) +
    poly(prot, 2)) + (1 | repfull), data = organsize, REML = F)
summary(sex.test.3)
```

```
## Linear mixed model fit by maximum likelihood . t-tests use Satterthwaite's
##   method [lmerModLmerTest]
## Formula: PCA1 ~ sex * (carb:prot + poly(carb, 2) + poly(prot, 2)) + (1 |
##     repfull)
##    Data: organsize
##
##      AIC      BIC   logLik deviance df.resid
##  -1067.7  -1002.9    547.9  -1095.7      743
##
## Scaled residuals:
##     Min      1Q  Median      3Q     Max
## -6.5636 -0.5882  0.0370  0.5871  3.6443
```

```
##
## Random effects:
##  Groups   Name        Variance Std.Dev.
##  repfull  (Intercept) 0.004162 0.06452
##  Residual             0.012289 0.11085
## Number of obs: 757, groups:  repfull, 61
##
## Fixed effects:
##                        Estimate Std. Error         df t value Pr(>|t|)
## (Intercept)           4.237e-02  4.866e-02  3.941e+01   0.871  0.38923
## sexM                 -3.198e-01  3.808e-02  7.299e+02  -8.399 2.34e-16 ***
## poly(carb, 2)1       -3.478e+00  7.310e-01  5.592e+01  -4.758 1.42e-05 ***
## poly(carb, 2)2        1.571e+00  3.288e-01  5.138e+01   4.779 1.51e-05 ***
## poly(prot, 2)1       -9.772e-02  1.020e+00  3.732e+01  -0.096  0.92420
## poly(prot, 2)2       -1.993e+00  3.138e-01  3.893e+01  -6.350 1.70e-07 ***
## carb:prot             1.631e-05  6.011e-06  3.975e+01   2.713  0.00981 **
## sexM:poly(carb, 2)1   1.611e+00  6.499e-01  7.491e+02   2.479  0.01340 *
## sexM:poly(carb, 2)2  -8.393e-01  2.827e-01  7.403e+02  -2.969  0.00308 **
## sexM:poly(prot, 2)1   7.212e-01  7.932e-01  7.286e+02   0.909  0.36358
## sexM:poly(prot, 2)2   4.068e-01  2.469e-01  7.346e+02   1.648  0.09986 .
## sexM:carb:prot       -7.631e-06  4.720e-06  7.313e+02  -1.617  0.10639
## ---
## Signif. codes:  0 '***' 0.001 '**' 0.01 '*' 0.05 '.' 0.1 ' ' 1
##
## Correlation of Fixed Effects:
##                (Intr) sexM   ply(c,2)1 ply(c,2)2 ply(p,2)1 ply(p,2)2 crb:pr
## sexM           -0.326
## ply(crb,2)1     0.863 -0.336
## ply(crb,2)2     0.001  0.022 -0.082
## ply(prt,2)1     0.919 -0.281  0.775     0.177
## ply(prt,2)2     0.157 -0.059  0.225     0.271     0.295
## carb:prot      -0.972  0.317 -0.892    -0.006    -0.932    -0.215
## sxM:ply(c,2)1 -0.295  0.891 -0.381     0.057    -0.246    -0.078     0.302
## sxM:ply(c,2)2  0.025  0.081  0.058    -0.394    -0.052    -0.038    -0.024
## sxM:ply(p,2)1 -0.281  0.908 -0.281    -0.068    -0.308    -0.046     0.290
## sxM:ply(p,2)2 -0.072  0.209 -0.108    -0.026    -0.053    -0.345     0.071
## sxM:crb:prt    0.316 -0.975  0.342    -0.020     0.290     0.058    -0.325
##                sxM:ply(c,2)1 sxM:ply(c,2)2 sxM:ply(p,2)1 sxM:ply(p,2)2
## sexM
## ply(crb,2)1
## ply(crb,2)2
## ply(prt,2)1
## ply(prt,2)2
## carb:prot
## sxM:ply(c,2)1
## sxM:ply(c,2)2  0.002
## sxM:ply(p,2)1  0.798         0.256
## sxM:ply(p,2)2  0.265         0.187         0.197
## sxM:crb:prt   -0.916        -0.073        -0.932        -0.210
## fit warnings:
## Some predictor variables are on very different scales: consider rescaling
```

```
anova(sex.test.0, sex.test.1, sex.test.2, sex.test.3)
```

```
## Data: organsize
```

```
## Models:
## sex.test.0: PCA1 ~ sex + carb:prot + poly(carb, 2) + poly(prot, 2) + (1 |
## sex.test.0:     repfull)
## sex.test.1: PCA1 ~ carb:prot + sex * poly(prot, 2) + poly(carb, 2) + (1 |
## sex.test.1:     repfull)
## sex.test.2: PCA1 ~ carb:prot + sex * poly(carb, 2) + poly(prot, 2) + (1 |
## sex.test.2:     repfull)
## sex.test.3: PCA1 ~ sex * (carb:prot + poly(carb, 2) + poly(prot, 2)) + (1 |
## sex.test.3:     repfull)
##            npar     AIC     BIC logLik deviance  Chisq Df Pr(>Chisq)
## sex.test.0    9 -1059.0 -1017.3 538.48  -1077.0
## sex.test.1   11 -1057.7 -1006.8 539.85  -1079.7 2.7298  2    0.25540
## sex.test.2   11 -1066.7 -1015.8 544.34  -1088.7 8.9777  0    < 2e-16 ***
## sex.test.3   14 -1067.7 -1002.9 547.85  -1095.7 7.0366  3    0.07074 .
## ---
## Signif. codes:  0 '***' 0.001 '**' 0.01 '*' 0.05 '.' 0.1 ' ' 1
```

Re-testing but this time using paramteric bootstrapping:

```
PBmodcomp(sex.test.1, sex.test.0, nsim = 1000, details = 1)
```

```
## Reference distribution with  1000 samples; computing time: 20.85 secs.

## Bootstrap test; time: 20.85 sec;samples: 1000; extremes: 276;
## large : PCA1 ~ carb:prot + sex * poly(prot, 2) + poly(carb, 2) + (1 |
##     repfull)
## small : PCA1 ~ sex + carb:prot + poly(carb, 2) + poly(prot, 2) + (1 |
##     repfull)
##          stat df p.value
## LRT    2.7298  2  0.2554
## PBtest 2.7298     0.2767
```

```
PBmodcomp(sex.test.2, sex.test.0, nsim = 1000, details = 1)
```

```
## Reference distribution with  1000 samples; computing time: 19.88 secs.

## Bootstrap test; time: 19.88 sec;samples: 1000; extremes: 0;
## large : PCA1 ~ carb:prot + sex * poly(carb, 2) + poly(prot, 2) + (1 |
##     repfull)
## small : PCA1 ~ sex + carb:prot + poly(carb, 2) + poly(prot, 2) + (1 |
##     repfull)
##          stat df  p.value
## LRT    11.707  2 0.002869 **
## PBtest 11.707    0.000999 ***
## ---
## Signif. codes:  0 '***' 0.001 '**' 0.01 '*' 0.05 '.' 0.1 ' ' 1
```

```
PBmodcomp(sex.test.3, sex.test.2, nsim = 1000, details = 1)
```

```
## Reference distribution with  1000 samples; computing time: 20.05 secs.

## Bootstrap test; time: 20.05 sec;samples: 1000; extremes: 74;
## large : PCA1 ~ sex * (carb:prot + poly(carb, 2) + poly(prot, 2)) + (1 |
##     repfull)
## small : PCA1 ~ carb:prot + sex * poly(carb, 2) + poly(prot, 2) + (1 |
##     repfull)
##          stat df p.value
## LRT    7.0366  3 0.07074 .
```

```
## PBtest 7.0366    0.07493 .
## ---
## Signif. codes:  0 '***' 0.001 '**' 0.01 '*' 0.05 '.' 0.1 ' ' 1
```

Finally, we can use Baysian analysis to do the same thing:

```
model1M.MCMC <- MCMCglmm(PCA1 ~ sex * (carb + I(carb^2) + prot +
    I(prot^2) + carb:prot), random = ~repfull, burnin = 5000,
    nitt = 20000, thin = 10, verbose = F, pr = T, data = organsize)
summary(model1M.MCMC)
```

```
##
##  Iterations = 5001:19991
##  Thinning interval  = 10
##  Sample size  = 1500
##
##  DIC: -1125.141
##
##  G-structure:  ~repfull
##
##         post.mean l-95% CI u-95% CI eff.samp
## repfull  0.005628 0.002395 0.009309     1313
##
##  R-structure:  ~units
##
##       post.mean l-95% CI u-95% CI eff.samp
## units   0.01238  0.01106  0.01369     1500
##
##  Location effects: PCA1 ~ sex * (carb + I(carb^2) + prot + I(prot^2) + carb:prot)
##
##                  post.mean   l-95% CI   u-95% CI eff.samp   pMCMC
## (Intercept)      1.345e-01  3.159e-02  2.343e-01     1500 0.00800 **
## sexM            -4.541e-01 -5.364e-01 -3.771e-01     1500 < 7e-04 ***
## carb            -4.341e-03 -5.970e-03 -3.010e-03     1500 < 7e-04 ***
## I(carb^2)        9.459e-06  5.095e-06  1.368e-05     1500 < 7e-04 ***
## prot             6.185e-03  3.660e-03  8.251e-03     1500 < 7e-04 ***
## I(prot^2)       -3.310e-05 -4.470e-05 -2.171e-05     1500 < 7e-04 ***
## carb:prot        1.585e-05  2.559e-06  3.003e-05     1373 0.03600 *
## sexM:carb        2.174e-03  1.063e-03  3.283e-03     1418 < 7e-04 ***
## sexM:I(carb^2) -4.934e-06 -8.178e-06 -1.625e-06     1500 0.00267 **
## sexM:prot       -7.390e-04 -2.499e-03  9.733e-04     1500 0.40800
## sexM:I(prot^2)   6.888e-06 -1.336e-06  1.510e-05     1500 0.10667
## sexM:carb:prot -7.552e-06 -1.662e-05  2.566e-06     1277 0.13200
## ---
## Signif. codes:  0 '***' 0.001 '**' 0.01 '*' 0.05 '.' 0.1 ' ' 1
```

Again, the analysis does not support inclusion of a sex*protein interaction.
