## Supplementary material for "Sex-specific plasticity and the nutritional geometry of insulin-signaling gene expression in *Drosophila melanogaster*": R Markdown of Gene Expression Analysis

### Gene Expression Analysis without Outliers

Alexander Shingleton

11/11/2020

#### Analysis of the Nutritional Geometry of Gene Expression in Males and Females

Preamble

```
setwd("~/Documents/Data/Nutritional Geometry of Gene Expression/Final Analysis/Expression Analysis")


suppressPackageStartupMessages({
    require("gdata")
    require("car")
    require("data.table")
    require("ggplot2")
    require("plyr")
    require("lsmeans")
    require("mgcv")
    require("nlme")
    require("AICcmodavg")
    require("fields")
    require("lme4")
    require("piecewiseSEM")
    require("gridExtra")
    require("lmSupport")
    require("WebPower")
    require("pwr")
})

source("R/functions.R")
```

Import the data and add additional columns specifying nutritional details.

```
data <- read.csv("gene expression data log.csv")

diets <- read.csv("diets gene expression.csv")

all.data <- within(data, {
    foodF <- factor(food)
    inter = interaction(ratio, foodF)
    logprot = log(prot)
    logcarb = log(carb)
    logratio = log(ratio_num)
    logfood = log(food)
```

```
})

all.data$cal <- data$food * 0.004
m.data <- subset(all.data, sex == "M")
f.data <- subset(all.data, sex == "F")
```

#### Multivariate Analysis of Gene Expression

Using Lee's model of nutritional geometry.

```
gexp <- as.matrix(cbind(all.data[, c(9:16)]))

all.manova0 <- lm(cbind(X4eBP, InR, dILP2, dILP3, dILP5, dILP8,
    Ash2L, CG3071) ~ sex * (prot:carb + poly(carb, 2) + poly(prot,
    2)), data = all.data)
all.manova1 <- lm(cbind(X4eBP, InR, dILP2, dILP3, dILP5, dILP8,
    Ash2L, CG3071) ~ sex + (prot:carb + poly(carb, 2) + poly(prot,
    2)), data = all.data)
anova(all.manova0, all.manova1)
```

```
## Analysis of Variance Table
##
## Model 1: cbind(X4eBP, InR, dILP2, dILP3, dILP5, dILP8, Ash2L, CG3071) ~
##     sex * (prot:carb + poly(carb, 2) + poly(prot, 2))
## Model 2: cbind(X4eBP, InR, dILP2, dILP3, dILP5, dILP8, Ash2L, CG3071) ~
##     sex + (prot:carb + poly(carb, 2) + poly(prot, 2))
##   Res.Df Df Gen.var.  Pillai approx F num Df den Df Pr(>F)
## 1     85     0.51583
## 2     90  5  0.53614 0.65981   1.5582     40    410 0.0191 *
## ---
## Signif. codes:  0 '***' 0.001 '**' 0.01 '*' 0.05 '.' 0.1 ' ' 1
```

```
Man.all.manova0 <- Manova(all.manova0, multivariate = TRUE, type = "III")
Man.all.manova0
```

```
##
## Type III MANOVA Tests: Pillai test statistic
##                    Df test stat approx F num Df den Df    Pr(>F)
## (Intercept)         1   0.67473  20.2254      8     78 3.471e-16 ***
## sex                 1   0.17797   2.1108      8     78 0.0444554 *
## poly(carb, 2)       2   0.26745   1.5244     16    158 0.0970869 .
## poly(prot, 2)       2   0.47337   3.0620     16    158 0.0001653 ***
## prot:carb           1   0.10890   1.1915      8     78 0.3150340
## sex:poly(carb, 2)  2   0.16264   0.8741     16    158 0.5998437
## sex:poly(prot, 2)  2   0.28750   1.6578     16    158 0.0601867 .
## sex:prot:carb       1   0.09032   0.9681      8     78 0.4670743
## ---
## Signif. codes:  0 '***' 0.001 '**' 0.01 '*' 0.05 '.' 0.1 ' ' 1
```

It looks like there is a significant sex:gene expression interaction, but inspecting the factors, the prot:carb and sex:carb^2 terms are not significant. Try a simpler model with a linear carbohydrate term:

```
all.manova0 <- lm(cbind(X4eBP, InR, dILP2, dILP3, dILP5, dILP8,
    Ash2L, CG3071) ~ sex * (prot:carb + poly(carb, 2) + poly(prot,
    2)), data = all.data)
```

```r
all.manova1 <- lm(cbind(X4eBP, InR, dILP2, dILP3, dILP5, dILP8,
    Ash2L, CG3071) ~ sex * (poly(carb, 1) + poly(prot, 2)), data = all.data)
all.manova2 <- lm(cbind(X4eBP, InR, dILP2, dILP3, dILP5, dILP8,
    Ash2L, CG3071) ~ sex + (poly(carb, 1) + poly(prot, 2)), data = all.data)
anova(all.manova2, all.manova1, all.manova0)
```

```
## Analysis of Variance Table
##
## Model 1: cbind(X4eBP, InR, dILP2, dILP3, dILP5, dILP8, Ash2L, CG3071) ~
##     sex + (poly(carb, 1) + poly(prot, 2))
## Model 2: cbind(X4eBP, InR, dILP2, dILP3, dILP5, dILP8, Ash2L, CG3071) ~
##     sex * (poly(carb, 1) + poly(prot, 2))
## Model 3: cbind(X4eBP, InR, dILP2, dILP3, dILP5, dILP8, Ash2L, CG3071) ~
##     sex * (prot:carb + poly(carb, 2) + poly(prot, 2))
##   Res.Df Df Gen.var.  Pillai approx F num Df den Df   Pr(>F)
## 1     92     0.53979
## 2     89 -3  0.51856 0.52584   2.1254     24    240 0.002304 **
## 3     85 -4  0.51583 0.37906   1.0599     32    324 0.383939
## ---
## Signif. codes:  0 '***' 0.001 '**' 0.01 '*' 0.05 '.' 0.1 ' ' 1
```

```r
Man.all.manova1 <- Manova(all.manova1, multivariate = TRUE, type = "III")
Man.all.manova1
```

```
##
## Type III MANOVA Tests: Pillai test statistic
##                    Df test stat approx F num Df den Df    Pr(>F)
## (Intercept)         1   0.90826  101.475      8     82 < 2.2e-16 ***
## sex                 1   0.49122    9.896      8     82 1.571e-09 ***
## poly(carb, 1)       1   0.11512    1.333      8     82   0.23883
## poly(prot, 2)       2   0.60333    4.482     16    166 2.382e-07 ***
## sex:poly(carb, 1)  1   0.19038    2.410      8     82   0.02173 *
## sex:poly(prot, 2)  2   0.30278    1.851     16    166   0.02852 *
## ---
## Signif. codes:  0 '***' 0.001 '**' 0.01 '*' 0.05 '.' 0.1 ' ' 1
```

It looks like there is a significant quadratic protein and linear carbohydrate interaction with sex

#### Gene Expression and Body Size

We can also explore how body gene expression of each gene change changes with body size. Because we used different diets for the body size and analysis and gene expression analysis, we will use fitted values to test the relationship.

First looking at gene expression and fitted values for overall body size.

We can also get the fitted PCA1 for the diets used in the gene expression analysis. First load the body size data and fit the PCA values predicted by the body size model to the expression data diets.

Load the body size data

```r
diets.gene.expression <- read.csv("~/Documents/Data/Nutritional Geometry of Gene Expression/Final Analy
organsize <- read.csv("~/Documents/Data/Nutritional Geometry of Gene Expression/Final Analysis/Body Siz
```

Now extract the females data, fit the complete model and add the predicted PCA values to the expression data.

```r
femsize <- (subset(organsize, sex == "F"))
femsize$genital <- NULL
femsize <- na.omit(femsize)

size.model.lee.replicates <- lmer(PCA1 ~ carb * prot + poly(carb,
    2) + poly(prot, 2) + (1 | repfull), data = femsize)
GE_fem_size <- cbind(diets.gene.expression, (predict(size.model.lee.replicates,
    diets.gene.expression, re.form = NA)))
names(GE_fem_size)[5] <- "lee"

GE_fem_size[, c(3, 4)] <- NULL
f.data <- merge(GE_fem_size, f.data, by = c("carb", "prot"))
```

Now do the same with the thin plate spline

```r
tps.f <- with(femsize, Tps(cbind(prot, carb), PCA1))
gen.tps.f <- within(diets.gene.expression, {
    tps <- predict(tps.f, data.frame(prot, carb))
})
gen.tps.f[, c(3, 4)] <- NULL
f.data <- merge(gen.tps.f, f.data, by = c("carb", "prot"))
```

Now males:

```r
malsize <- (subset(organsize, sex == "M"))
malsize <- na.omit(malsize)

size.model.lee.replicates <- lmer(PCA1 ~ poly(prot, 2) + (1 |
    repfull), data = malsize)
GE_mal_size <- cbind(diets.gene.expression, (predict(size.model.lee.replicates,
    diets.gene.expression, re.form = NA)))
names(GE_mal_size)[5] <- "lee"

GE_mal_size[, c(3, 4)] <- NULL
m.data <- merge(GE_mal_size, m.data, by = c("carb", "prot"))

tps.m <- with(malsize, Tps(cbind(prot, carb), PCA1))
gen.tps.m <- within(diets.gene.expression, {
    tps <- predict(tps.m, data.frame(prot, carb))
})
gen.tps.m[, c(3, 4)] <- NULL
m.data <- merge(gen.tps.m, m.data, by = c("carb", "prot"))
```

The expression of which gene correlates most strongly with body size?

```r
fem_lm_lee <- lm(lee ~ X4eBP + InR + dILP2 + dILP3 + dILP5 +
    dILP8 + Ash2L + CG3071, data = f.data)
Anova(fem_lm_lee)
```

```
## Anova Table (Type II tests)
##
## Response: lee
##             Sum Sq Df F value   Pr(>F)
## X4eBP      0.16378  1  9.7805 0.002777 **
## InR        0.00375  1  0.2241 0.637754
## dILP2      0.01772  1  1.0581 0.308001
```

```
## dILP3     0.00048  1  0.0288 0.865893
## dILP5     0.07280  1  4.3477 0.041548 *
## dILP8     0.07847  1  4.6859 0.034612 *
## Ash2L     0.00327  1  0.1951 0.660389
## CG3071    0.03812  1  2.2762 0.136893
## Residuals 0.95449 57
## ---
## Signif. codes:  0 '***' 0.001 '**' 0.01 '*' 0.05 '.' 0.1 ' ' 1
```

```
summary(fem_lm_lee)
```

```
##
## Call:
## lm(formula = lee ~ X4eBP + InR + dILP2 + dILP3 + dILP5 + dILP8 +
##     Ash2L + CG3071, data = f.data)
##
## Residuals:
##      Min       1Q   Median       3Q      Max
## -0.30673 -0.08027  0.01124  0.08127  0.20740
##
## Coefficients:
##               Estimate Std. Error t value Pr(>|t|)
## (Intercept)  0.0003867  0.0724318   0.005  0.99576
## X4eBP       -0.1225625  0.0391901  -3.127  0.00278 **
## InR         -0.0135075  0.0285345  -0.473  0.63775
## dILP2        0.0390298  0.0379437   1.029  0.30800
## dILP3       -0.0104325  0.0614972  -0.170  0.86589
## dILP5        0.0544697  0.0261230   2.085  0.04155 *
## dILP8       -0.0252488  0.0116639  -2.165  0.03461 *
## Ash2L        0.0141105  0.0319471   0.442  0.66039
## CG3071      -0.0527754  0.0349801  -1.509  0.13689
## ---
## Signif. codes:  0 '***' 0.001 '**' 0.01 '*' 0.05 '.' 0.1 ' ' 1
##
## Residual standard error: 0.1294 on 57 degrees of freedom
##   (18 observations deleted due to missingness)
## Multiple R-squared:  0.3759, Adjusted R-squared:  0.2883
## F-statistic: 4.291 on 8 and 57 DF,  p-value: 0.0004363
```

```
modelEffectSizes(fem_lm_lee)
```

```
## lm(formula = lee ~ X4eBP + InR + dILP2 + dILP3 + dILP5 + dILP8 +
##     Ash2L + CG3071, data = f.data)
##
## Coefficients
##                SSR df pEta-sqr dR-sqr
## (Intercept) 0.0000  1   0.0000     NA
## X4eBP       0.1638  1   0.1465 0.1071
## InR         0.0038  1   0.0039 0.0025
## dILP2       0.0177  1   0.0182 0.0116
## dILP3       0.0005  1   0.0005 0.0003
## dILP5       0.0728  1   0.0709 0.0476
## dILP8       0.0785  1   0.0760 0.0513
## Ash2L       0.0033  1   0.0034 0.0021
## CG3071      0.0381  1   0.0384 0.0249
```

```
##
## Sum of squared errors (SSE): 1.0
## Sum of squared total  (SST): 1.5
```

```r
fem_lm_tps <- lm(tps ~ X4eBP + InR + dILP2 + dILP3 + dILP5 +
    dILP8 + Ash2L + CG3071, data = f.data)
Anova(fem_lm_tps)
```

```
## Anova Table (Type II tests)
##
## Response: tps
##            Sum Sq Df F value   Pr(>F)
## X4eBP     0.09389  1  9.2536 0.003549 **
## InR       0.00230  1  0.2263 0.636104
## dILP2     0.01412  1  1.3919 0.242984
## dILP3     0.00169  1  0.1663 0.684909
## dILP5     0.05663  1  5.5816 0.021586 *
## dILP8     0.04274  1  4.2124 0.044732 *
## Ash2L     0.00338  1  0.3332 0.566068
## CG3071    0.03101  1  3.0569 0.085780 .
## Residuals 0.57832 57
## ---
## Signif. codes:  0 '***' 0.001 '**' 0.01 '*' 0.05 '.' 0.1 ' ' 1
```

```r
modelEffectSizes(fem_lm_tps)
```

```
## lm(formula = tps ~ X4eBP + InR + dILP2 + dILP3 + dILP5 + dILP8 +
##     Ash2L + CG3071, data = f.data)
##
## Coefficients
##                SSR df pEta-sqr dR-sqr
## (Intercept) 0.0054  1   0.0092     NA
## X4eBP       0.0939  1   0.1397 0.0987
## InR         0.0023  1   0.0040 0.0024
## dILP2       0.0141  1   0.0238 0.0148
## dILP3       0.0017  1   0.0029 0.0018
## dILP5       0.0566  1   0.0892 0.0595
## dILP8       0.0427  1   0.0688 0.0449
## Ash2L       0.0034  1   0.0058 0.0036
## CG3071      0.0310  1   0.0509 0.0326
##
## Sum of squared errors (SSE): 0.6
## Sum of squared total  (SST): 1.0
```

```r
mal_lm_lee <- lm(lee ~ X4eBP + InR + dILP2 + dILP3 + dILP5 +
    dILP8 + Ash2L + CG3071, data = m.data)
Anova(mal_lm_lee)
```

```
## Anova Table (Type II tests)
##
## Response: lee
##             Sum Sq Df F value  Pr(>F)
## X4eBP     0.002047  1  0.2941 0.59303
## InR       0.030943  1  4.4467 0.04658 *
## dILP2     0.011795  1  1.6950 0.20640
## dILP3     0.013289  1  1.9097 0.18087
```

```
## dILP5     0.026073  1  3.7469 0.06586 .
## dILP8     0.000010  1  0.0015 0.96977
## Ash2L     0.005330  1  0.7659 0.39094
## CG3071    0.002626  1  0.3774 0.54530
## Residuals 0.153089 22
## ---
## Signif. codes:  0 '***' 0.001 '**' 0.01 '*' 0.05 '.' 0.1 ' ' 1
```

```r
summary(mal_lm_lee)
```

```
##
## Call:
## lm(formula = lee ~ X4eBP + InR + dILP2 + dILP3 + dILP5 + dILP8 +
##     Ash2L + CG3071, data = m.data)
##
## Residuals:
##       Min        1Q    Median        3Q       Max
## -0.129064 -0.051501 -0.008355  0.030318  0.222267
##
## Coefficients:
##              Estimate Std. Error t value Pr(>|t|)
## (Intercept) -0.450970   0.058608  -7.695 1.12e-07 ***
## X4eBP        0.018454   0.034027   0.542   0.5930
## InR         -0.038244   0.018136  -2.109   0.0466 *
## dILP2       -0.018943   0.014550  -1.302   0.2064
## dILP3       -0.018031   0.013048  -1.382   0.1809
## dILP5        0.047342   0.024457   1.936   0.0659 .
## dILP8        0.000499   0.013019   0.038   0.9698
## Ash2L        0.019158   0.021892   0.875   0.3909
## CG3071      -0.020244   0.032952  -0.614   0.5453
## ---
## Signif. codes:  0 '***' 0.001 '**' 0.01 '*' 0.05 '.' 0.1 ' ' 1
##
## Residual standard error: 0.08342 on 22 degrees of freedom
##   (44 observations deleted due to missingness)
## Multiple R-squared:  0.3817, Adjusted R-squared:  0.1569
## F-statistic: 1.698 on 8 and 22 DF,  p-value: 0.155
```

```r
modelEffectSizes(mal_lm_lee)
```

```
## lm(formula = lee ~ X4eBP + InR + dILP2 + dILP3 + dILP5 + dILP8 +
##     Ash2L + CG3071, data = m.data)
##
## Coefficients
##                SSR df pEta-sqr dR-sqr
## (Intercept) 0.4120  1   0.7291     NA
## X4eBP       0.0020  1   0.0132 0.0083
## InR         0.0309  1   0.1681 0.1250
## dILP2       0.0118  1   0.0715 0.0476
## dILP3       0.0133  1   0.0799 0.0537
## dILP5       0.0261  1   0.1455 0.1053
## dILP8       0.0000  1   0.0001 0.0000
## Ash2L       0.0053  1   0.0336 0.0215
## CG3071      0.0026  1   0.0169 0.0106
##
```

```
## Sum of squared errors (SSE): 0.2
## Sum of squared total  (SST): 0.2
```

```
mal_lm_tps <- lm(tps ~ X4eBP + InR + dILP2 + dILP3 + dILP5 +
    dILP8 + Ash2L + CG3071, data = m.data)
Anova(mal_lm_tps)
```

```
## Anova Table (Type II tests)
##
## Response: tps
##             Sum Sq Df F value  Pr(>F)
## X4eBP     0.000942  1  0.1486 0.70354
## InR       0.004025  1  0.6350 0.43406
## dILP2     0.000821  1  0.1296 0.72229
## dILP3     0.004038  1  0.6371 0.43329
## dILP5     0.019218  1  3.0318 0.09561 .
## dILP8     0.002093  1  0.3301 0.57140
## Ash2L     0.000115  1  0.0182 0.89396
## CG3071    0.000130  1  0.0206 0.88727
## Residuals 0.139449 22
## ---
## Signif. codes:  0 '***' 0.001 '**' 0.01 '*' 0.05 '.' 0.1 ' ' 1
```

```
modelEffectSizes(mal_lm_tps)
```

```
## lm(formula = tps ~ X4eBP + InR + dILP2 + dILP3 + dILP5 + dILP8 +
##     Ash2L + CG3071, data = m.data)
##
## Coefficients
##                SSR df pEta-sqr dR-sqr
## (Intercept) 0.2494  1   0.6414     NA
## X4eBP       0.0009  1   0.0067 0.0050
## InR         0.0040  1   0.0281 0.0212
## dILP2       0.0008  1   0.0059 0.0043
## dILP3       0.0040  1   0.0281 0.0212
## dILP5       0.0192  1   0.1211 0.1011
## dILP8       0.0021  1   0.0148 0.0110
## Ash2L       0.0001  1   0.0008 0.0006
## CG3071      0.0001  1   0.0009 0.0007
##
## Sum of squared errors (SSE): 0.1
## Sum of squared total  (SST): 0.2
```

Now lets examine the nutritional geometry of each gene's expression individually.

#### Analysis of InR

Test whether there is a difference between the sexes

```
InRModel1 <- lm(InR ~ sex + (carb:prot + poly(carb, 2) + poly(prot,
    2)), data = all.data)
InRModel2 <- lm(InR ~ sex * (carb:prot + poly(carb, 2) + poly(prot,
    2)), data = all.data)
anova(InRModel1, InRModel2)
```

```
## Analysis of Variance Table
```

```
##
## Model 1: InR ~ sex + (carb:prot + poly(carb, 2) + poly(prot, 2))
## Model 2: InR ~ sex * (carb:prot + poly(carb, 2) + poly(prot, 2))
##   Res.Df    RSS Df Sum of Sq      F Pr(>F)
## 1    144 92.907
## 2    139 89.474  5    3.4329 1.0666 0.3816
```

No difference.

Test the effect of C and P on InR expression in males and females

```r
InR.m <- lm(InR ~ (carb:prot + poly(carb, 2) + poly(prot, 2)),
    data = m.data)
InR.f <- lm(InR ~ (carb:prot + poly(carb, 2) + poly(prot, 2)),
    data = f.data)
summary(InR.m)
```

```
##
## Call:
## lm(formula = InR ~ (carb:prot + poly(carb, 2) + poly(prot, 2)),
##     data = m.data)
##
## Residuals:
##     Min      1Q  Median      3Q     Max
## -2.1976 -0.4168  0.1617  0.6228  2.5751
##
## Coefficients:
##                   Estimate Std. Error t value Pr(>|t|)
## (Intercept)     -1.025e+00  3.094e-01  -3.314  0.00153 **
## poly(carb, 2)1 -3.546e+00  1.842e+00  -1.926  0.05868 .
## poly(carb, 2)2  1.800e+00  1.068e+00   1.686  0.09682 .
## poly(prot, 2)1 -6.030e+00  2.376e+00  -2.538  0.01363 *
## poly(prot, 2)2  2.782e+00  1.068e+00   2.605  0.01145 *
## carb:prot       1.328e-04  6.824e-05   1.945  0.05621 .
## ---
## Signif. codes:  0 '***' 0.001 '**' 0.01 '*' 0.05 '.' 0.1 ' ' 1
##
## Residual standard error: 0.9518 on 63 degrees of freedom
##   (6 observations deleted due to missingness)
## Multiple R-squared:  0.228,  Adjusted R-squared:  0.1668
## F-statistic: 3.722 on 5 and 63 DF,  p-value: 0.005145
```

```r
summary(InR.f)
```

```
##
## Call:
## lm(formula = InR ~ (carb:prot + poly(carb, 2) + poly(prot, 2)),
##     data = f.data)
##
## Residuals:
##     Min      1Q  Median      3Q     Max
## -2.9826 -0.2982  0.1193  0.3427  1.1351
##
## Coefficients:
##                   Estimate Std. Error t value Pr(>|t|)
## (Intercept)     -1.898e-01  1.874e-01  -1.013   0.3142
```

```
## poly(carb, 2)1 -1.591e+00  1.315e+00  -1.209   0.2303
## poly(carb, 2)2 -3.863e-01  7.467e-01  -0.517   0.6064
## poly(prot, 2)1 -3.020e+00  1.517e+00  -1.990   0.0502 .
## poly(prot, 2)2  1.659e+00  6.770e-01   2.451   0.0166 *
## carb:prot       1.620e-05  4.296e-05   0.377   0.7071
## ---
## Signif. codes:  0 '***' 0.001 '**' 0.01 '*' 0.05 '.' 0.1 ' ' 1
##
## Residual standard error: 0.653 on 76 degrees of freedom
##   (2 observations deleted due to missingness)
## Multiple R-squared:  0.2577, Adjusted R-squared:  0.2088
## F-statistic: 5.276 on 5 and 76 DF,  p-value: 0.0003264
```

There is a quadratic protein effect in males and females

Reanalyse the data using the simplest model (the carb effect in males disappears when the carb*protein interaction is dropped, suggesting that it is not well supported)

```
# Non-orthogoganol coef
summary(lm(InR ~ (poly(prot, 2, raw = TRUE)), data = m.data))[4]
```

```
## $coefficients
##                                Estimate   Std. Error   t value    Pr(>|t|)
## (Intercept)                0.3232356028 2.552972e-01  1.266115 0.209922632
## poly(prot, 2, raw = TRUE)1 -0.0333977270 1.126691e-02 -2.964232 0.004218233
## poly(prot, 2, raw = TRUE)2  0.0001874908 7.988717e-05  2.346945 0.021938116
```

```
summary(lm(InR ~ (poly(prot, 2, raw = TRUE)), data = f.data))[4]
```

```
## $coefficients
##                                Estimate   Std. Error   t value     Pr(>|t|)
## (Intercept)                0.4805635111 1.505834e-01  3.191344 0.0020326380
## poly(prot, 2, raw = TRUE)1 -0.0254449192 6.581802e-03 -3.865950 0.0002258192
## poly(prot, 2, raw = TRUE)2  0.0001361743 4.688718e-05  2.904297 0.0047711815
```

```
# Orthoganol test
InR.m <- lm(InR ~ (poly(prot, 2)), data = m.data)
InR.f <- lm(InR ~ (poly(prot, 2)), data = f.data)
summary(InR.m)[4]
```

```
## $coefficients
##                  Estimate Std. Error   t value     Pr(>|t|)
## (Intercept)    -0.4650434  0.1174132 -3.960742 0.0001857595
## poly(prot, 2)1 -2.6342643  1.0683063 -2.465832 0.0162763298
## poly(prot, 2)2  2.4686947  1.0518756  2.346945 0.0219381160
```

```
summary(InR.f)[4]
```

```
## $coefficients
##                  Estimate Std. Error   t value     Pr(>|t|)
## (Intercept)    -0.1191192 0.07214818 -1.651035 0.1027015008
## poly(prot, 2)1 -2.5520932 0.65720556 -3.883250 0.0002127483
## poly(prot, 2)2  1.9058965 0.65623331  2.904297 0.0047711815
```

Re-test whether there is a difference between the sexes

```
InRModel0 <- lm(InR ~ (poly(prot, 2)), data = all.data)
InRModel1 <- lm(InR ~ sex + (poly(prot, 2)), data = all.data)
InRModel2 <- lm(InR ~ sex * (poly(prot, 2)), data = all.data)
```

```r
anova(InRModel0, InRModel1, InRModel2)
```

```
## Analysis of Variance Table
##
## Model 1: InR ~ (poly(prot, 2))
## Model 2: InR ~ sex + (poly(prot, 2))
## Model 3: InR ~ sex * (poly(prot, 2))
##   Res.Df     RSS Df Sum of Sq      F  Pr(>F)
## 1    148 100.209
## 2    147  96.398  1    3.8116 5.7500 0.01776 *
## 3    145  96.120  2    0.2781 0.2098 0.81103
## ---
## Signif. codes:  0 '***' 0.001 '**' 0.01 '*' 0.05 '.' 0.1 ' ' 1
```

```r
summary(InRModel1)
```

```
##
## Call:
## lm(formula = InR ~ sex + (poly(prot, 2)), data = all.data)
##
## Residuals:
##     Min      1Q  Median      3Q     Max
## -2.9285 -0.4677  0.1439  0.4885  2.8730
##
## Coefficients:
##                 Estimate Std. Error t value Pr(>|t|)
## (Intercept)     -0.13071    0.08943  -1.462 0.145983
## sexM            -0.31900    0.13232  -2.411 0.017148 *
## poly(prot, 2)1 -3.66252    0.84858  -4.316  2.9e-05 ***
## poly(prot, 2)2  3.04381    0.84187   3.616 0.000411 ***
## ---
## Signif. codes:  0 '***' 0.001 '**' 0.01 '*' 0.05 '.' 0.1 ' ' 1
##
## Residual standard error: 0.8098 on 147 degrees of freedom
##   (8 observations deleted due to missingness)
## Multiple R-squared:  0.2054, Adjusted R-squared:  0.1892
## F-statistic: 12.67 on 3 and 147 DF,  p-value: 2.061e-07
```

There is still no effect of sex on InR expression althought it is slightly lower in males than females.

Plot the model and the TPS

```
## Warning:
## Grid searches over lambda (nugget and sill variances) with  minima at the endpoints:
##   (GCV) Generalized Cross-Validation
##    minimum at  right endpoint  lambda  =  275.416 (eff. df= 3.000995 )
```

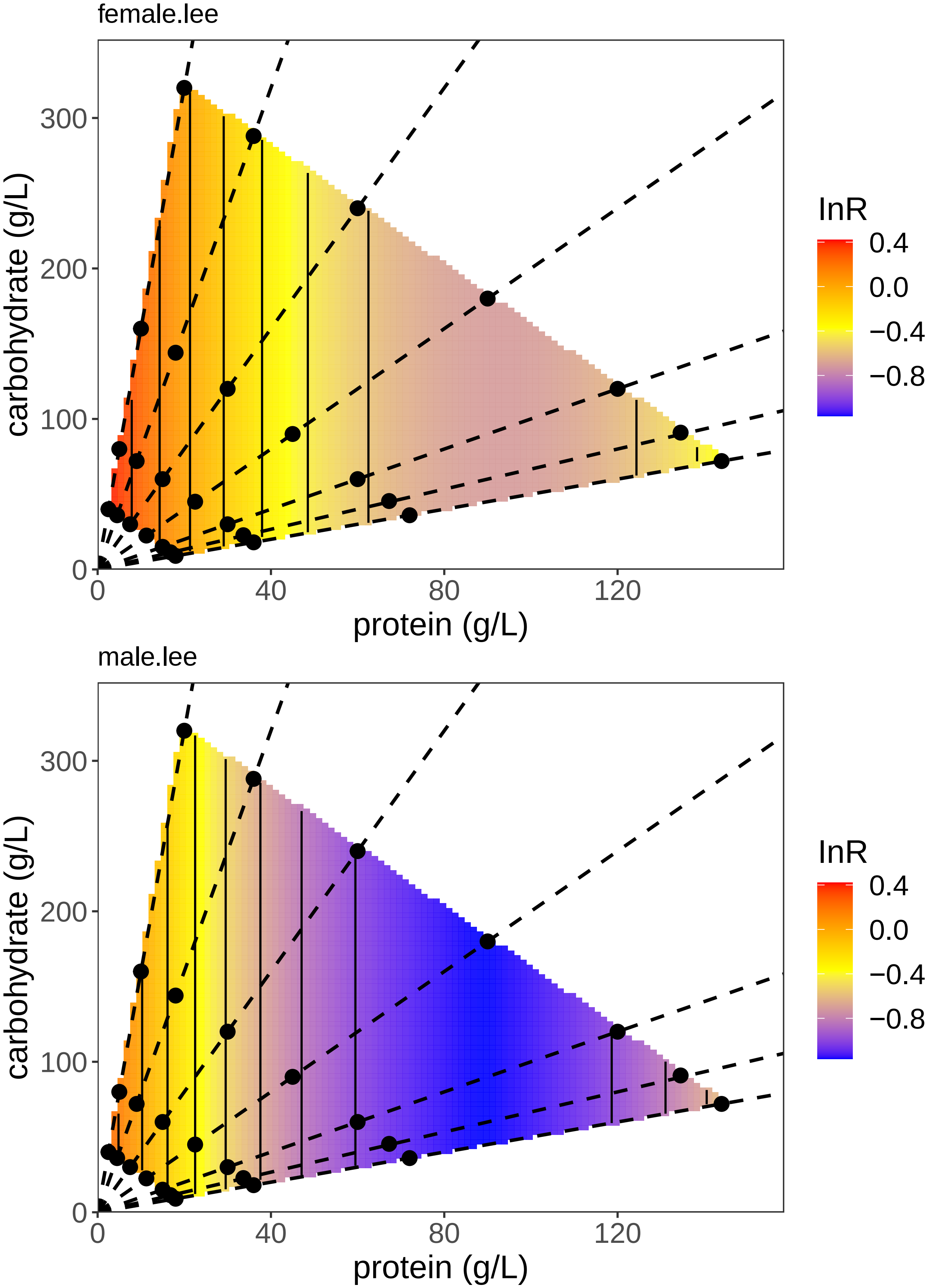
female.lee
carbohydrate (g/L)
300
200
100
0
0
40
80
120
protein (g/L)
lnR
0.4
0.0
−0.4
−0.8
male.lee
carbohydrate (g/L)
300
200
100
0
0
40
80
120
protein (g/L)
lnR
0.4
0.0
−0.4
−0.8

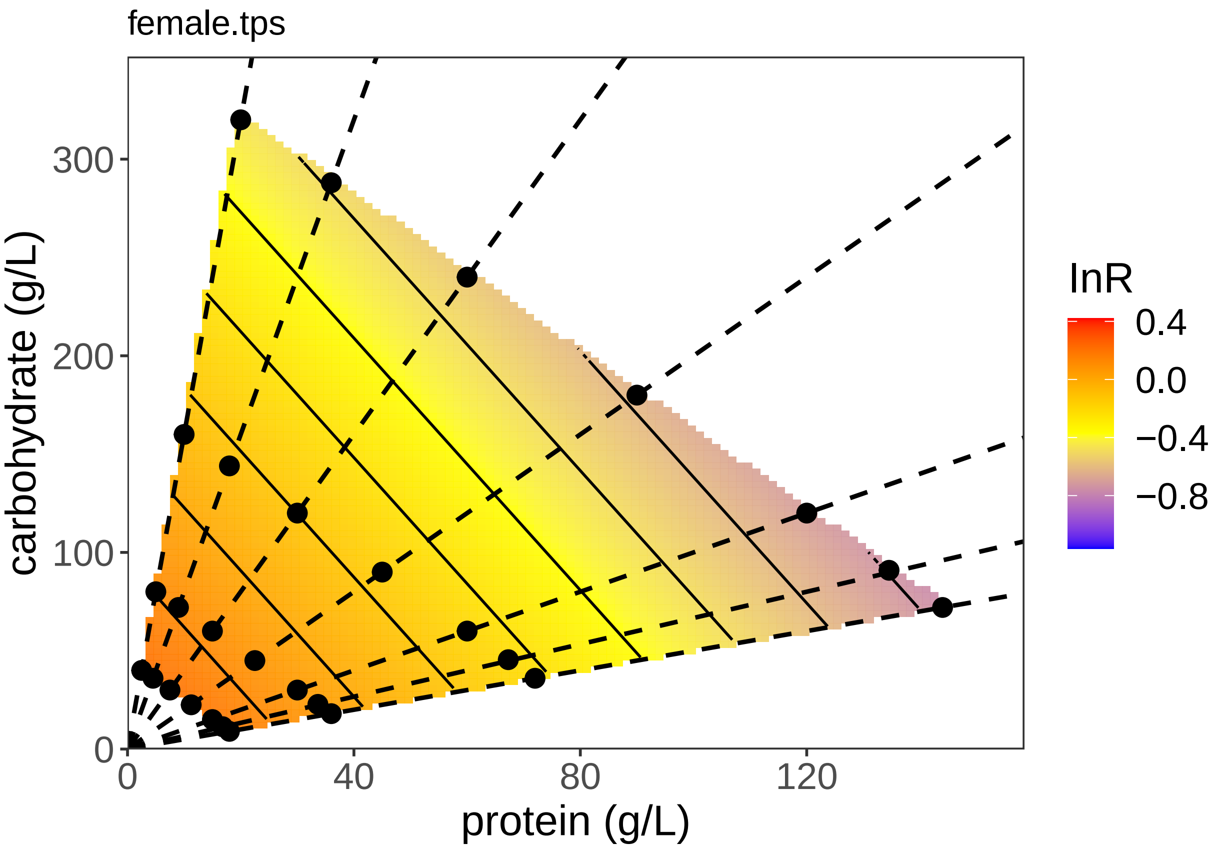


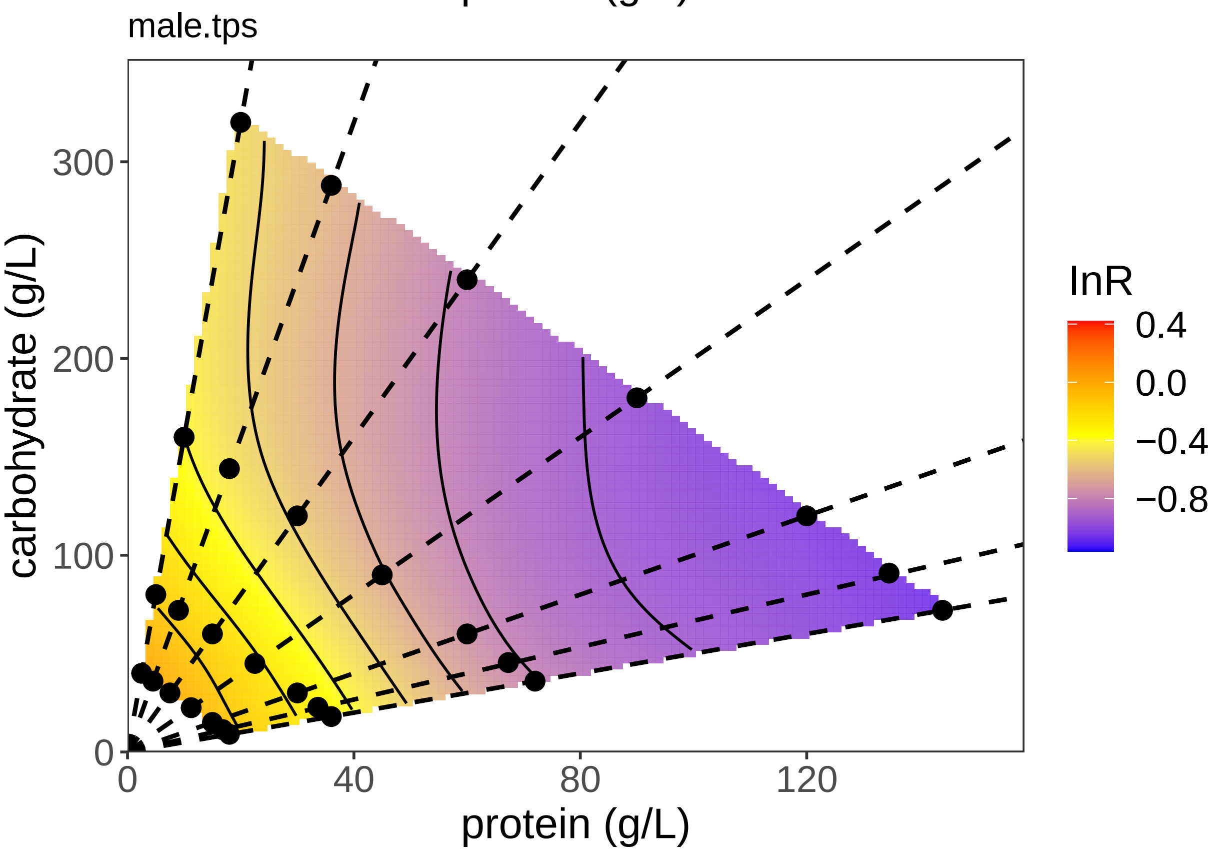


##Analysis of 4eBP

Test whether there is a difference between the sexes

```r
X4eBPModel1 <- lm(X4eBP ~ sex + (carb:prot + poly(carb, 2) +
    poly(prot, 2)), data = all.data)
X4eBPModel2 <- lm(X4eBP ~ sex * (carb:prot + poly(carb, 2) +
    poly(prot, 2)), data = all.data)
anova(X4eBPModel1, X4eBPModel2)
```

```
## Analysis of Variance Table
##
## Model 1: X4eBP ~ sex + (carb:prot + poly(carb, 2) + poly(prot, 2))
## Model 2: X4eBP ~ sex * (carb:prot + poly(carb, 2) + poly(prot, 2))
##   Res.Df    RSS Df Sum of Sq      F  Pr(>F)
## 1    149 50.578
## 2    144 46.530  5    4.0481 2.5056 0.03296 *
## ---
## Signif. codes:  0 '***' 0.001 '**' 0.01 '*' 0.05 '.' 0.1 ' ' 1
```

```r
summary(X4eBPModel2)
```

```
##
## Call:
## lm(formula = X4eBP ~ sex * (carb:prot + poly(carb, 2) + poly(prot,
##     2)), data = all.data)
##
## Residuals:
##      Min       1Q   Median       3Q      Max
## -1.58347 -0.24844  0.05244  0.35367  1.26846
##
## Coefficients:
##                         Estimate Std. Error t value Pr(>|t|)
## (Intercept)            4.303e-02  1.652e-01   0.260   0.7949
## sexM                  -2.106e-01  2.394e-01  -0.880   0.3805
## poly(carb, 2)1        -2.877e-01  1.551e+00  -0.186   0.8531
## poly(carb, 2)2        -1.201e-01  8.889e-01  -0.135   0.8927
## poly(prot, 2)1        -3.791e+00  1.837e+00  -2.063   0.0409 *
## poly(prot, 2)2        -2.216e-01  8.098e-01  -0.274   0.7847
## carb:prot              2.521e-05  3.740e-05   0.674   0.5014
## sexM:poly(carb, 2)1  5.980e-01  2.240e+00   0.267   0.7899
## sexM:poly(carb, 2)2  6.977e-01  1.284e+00   0.543   0.5877
## sexM:poly(prot, 2)1  3.383e+00  2.662e+00   1.271   0.2058
## sexM:poly(prot, 2)2  1.238e-02  1.180e+00   0.010   0.9916
## sexM:carb:prot         1.314e-05  5.440e-05   0.241   0.8095
## ---
## Signif. codes:  0 '***' 0.001 '**' 0.01 '*' 0.05 '.' 0.1 ' ' 1
##
## Residual standard error: 0.5684 on 144 degrees of freedom
##   (3 observations deleted due to missingness)
## Multiple R-squared:  0.1325, Adjusted R-squared:  0.06626
## F-statistic:     2 on 11 and 144 DF,  p-value: 0.03235
```

There is a sex*diet interaction, but none of the indiviual parameters in the full model are significant apart from prot^2.

Test the effect of C and P on 4EBP expression in males and females

```r
X4eBP.m <- lm(X4eBP ~ (carb:prot + poly(carb, 2) + poly(prot,
    2)), data = m.data)
X4eBP.f <- lm(X4eBP ~ (carb:prot + poly(carb, 2) + poly(prot,
    2)), data = f.data)
summary(X4eBP.m)
```

```
##
## Call:
## lm(formula = X4eBP ~ (carb:prot + poly(carb, 2) + poly(prot,
##     2)), data = m.data)
##
## Residuals:
##      Min       1Q   Median       3Q      Max
## -1.58347 -0.23334  0.09196  0.38112  1.26846
##
## Coefficients:
##                   Estimate Std. Error t value Pr(>|t|)
## (Intercept)     -1.693e-01  1.946e-01  -0.870    0.387
## poly(carb, 2)1  1.934e-01  1.209e+00   0.160    0.873
## poly(carb, 2)2  3.934e-01  6.990e-01   0.563    0.575
## poly(prot, 2)1 -2.890e-01  1.481e+00  -0.195    0.846
## poly(prot, 2)2 -1.433e-01  6.509e-01  -0.220    0.826
## carb:prot       3.834e-05  4.375e-05   0.876    0.384
##
## Residual standard error: 0.6295 on 68 degrees of freedom
##   (1 observation deleted due to missingness)
## Multiple R-squared:  0.08459,    Adjusted R-squared:  0.01728
## F-statistic: 1.257 on 5 and 68 DF,  p-value: 0.2928
```

```r
summary(X4eBP.f)
```

```
##
## Call:
## lm(formula = X4eBP ~ (carb:prot + poly(carb, 2) + poly(prot,
##     2)), data = f.data)
##
## Residuals:
##      Min       1Q   Median       3Q      Max
## -1.55728 -0.28149  0.03724  0.34331  1.08779
##
## Coefficients:
##                   Estimate Std. Error t value Pr(>|t|)
## (Intercept)      5.089e-02  1.457e-01   0.349    0.728
## poly(carb, 2)1 -2.157e-01  1.022e+00  -0.211    0.834
## poly(carb, 2)2 -8.783e-02  5.804e-01  -0.151    0.880
## poly(prot, 2)1 -2.717e+00  1.180e+00  -2.304    0.024 *
## poly(prot, 2)2 -1.613e-01  5.262e-01  -0.306    0.760
## carb:prot        2.521e-05  3.340e-05   0.755    0.453
## ---
## Signif. codes:  0 '***' 0.001 '**' 0.01 '*' 0.05 '.' 0.1 ' ' 1
##
## Residual standard error: 0.5076 on 76 degrees of freedom
##   (2 observations deleted due to missingness)
## Multiple R-squared:  0.1572, Adjusted R-squared:  0.1017
```

```
## F-statistic: 2.835 on 5 and 76 DF,  p-value: 0.02116
```

There is no effect of diet on x4eBP expression except prot in females.

Try a simpler model.

```
X4eBP.m <- lm(X4eBP ~ prot, data = m.data)
X4eBP.f <- lm(X4eBP ~ prot, data = f.data)
summary(X4eBP.m)
```

```
##
## Call:
## lm(formula = X4eBP ~ prot, data = m.data)
##
## Residuals:
##     Min      1Q  Median      3Q     Max
## -1.6630 -0.2768  0.1314  0.4481  1.2299
##
## Coefficients:
##              Estimate Std. Error t value Pr(>|t|)
## (Intercept) -0.122570   0.108210  -1.133    0.261
## prot         0.002613   0.001834   1.424    0.159
##
## Residual standard error: 0.6306 on 72 degrees of freedom
##   (1 observation deleted due to missingness)
## Multiple R-squared:  0.02741,    Adjusted R-squared:  0.0139
## F-statistic: 2.029 on 1 and 72 DF,  p-value: 0.1586
```

```
summary(X4eBP.f)
```

```
##
## Call:
## lm(formula = X4eBP ~ prot, data = f.data)
##
## Residuals:
##      Min       1Q   Median       3Q      Max
## -1.59712 -0.29330  0.07305  0.34584  1.06245
##
## Coefficients:
##              Estimate Std. Error t value Pr(>|t|)
## (Intercept)  0.357451   0.079935   4.472 2.53e-05 ***
## prot        -0.005082   0.001406  -3.614 0.000524 ***
## ---
## Signif. codes:  0 '***' 0.001 '**' 0.01 '*' 0.05 '.' 0.1 ' ' 1
##
## Residual standard error: 0.4997 on 80 degrees of freedom
##   (2 observations deleted due to missingness)
## Multiple R-squared:  0.1404, Adjusted R-squared:  0.1296
## F-statistic: 13.06 on 1 and 80 DF,  p-value: 0.0005244
```

There is an effect in females but not in males.

Compare sex effect again.

```
X4eBPModel1 <- lm(X4eBP ~ sex + prot, data = all.data)
X4eBPModel2 <- lm(X4eBP ~ sex * prot, data = all.data)
anova(X4eBPModel1, X4eBPModel2)
```

```
## Analysis of Variance Table
##
## Model 1: X4eBP ~ sex + prot
## Model 2: X4eBP ~ sex * prot
##   Res.Df    RSS Df Sum of Sq      F    Pr(>F)
## 1    153 52.219
## 2    152 48.604  1    3.6148 11.305 0.0009782 ***
## ---
## Signif. codes:  0 '***' 0.001 '**' 0.01 '*' 0.05 '.' 0.1 ' ' 1
```

```
summary(X4eBPModel2)
```

```
##
## Call:
## lm(formula = X4eBP ~ sex * prot, data = all.data)
##
## Residuals:
##      Min       1Q   Median       3Q      Max
## -1.66303 -0.28606  0.07854  0.39899  1.22990
##
## Coefficients:
##              Estimate Std. Error t value Pr(>|t|)
## (Intercept)  0.357451   0.090465   3.951 0.000119 ***
## sexM        -0.480021   0.132663  -3.618 0.000403 ***
## prot        -0.005082   0.001591  -3.194 0.001707 **
## sexM:prot    0.007695   0.002289   3.362 0.000978 ***
## ---
## Signif. codes:  0 '***' 0.001 '**' 0.01 '*' 0.05 '.' 0.1 ' ' 1
##
## Residual standard error: 0.5655 on 152 degrees of freedom
##   (3 observations deleted due to missingness)
## Multiple R-squared:  0.09386,    Adjusted R-squared:  0.07598
## F-statistic: 5.248 on 3 and 152 DF,  p-value: 0.001791
```

There is a significant difference between sexes in the response to protein. Males appear to have lower expression in general, but there is an interaction with protein, such that there is no effect on expression in males.

Plot the model and the TPS

```
## Warning:
## Grid searches over lambda (nugget and sill variances) with  minima at the endpoints:
##   (GCV) Generalized Cross-Validation
##    minimum at  right endpoint  lambda  =  275.416 (eff. df= 3.000995 )
```

```
## Warning:
## Grid searches over lambda (nugget and sill variances) with  minima at the endpoints:
##   (GCV) Generalized Cross-Validation
##    minimum at  right endpoint  lambda  =  246.4776 (eff. df= 3.001005 )
```

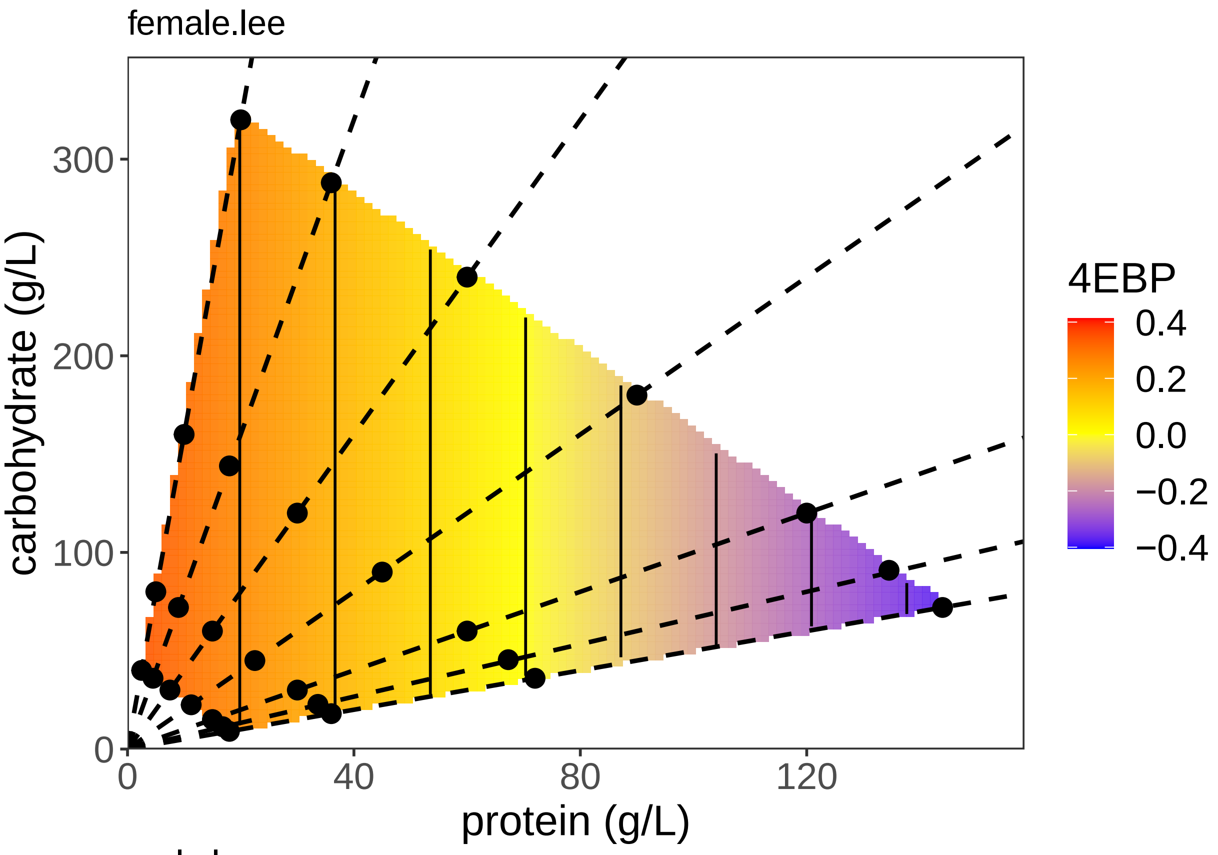
female.lee
carbohydrate (g/L)
protein (g/L)
0
100
200
300
0
40
80
120
4EBP
0.4
0.2
0.0
−0.2
−0.4

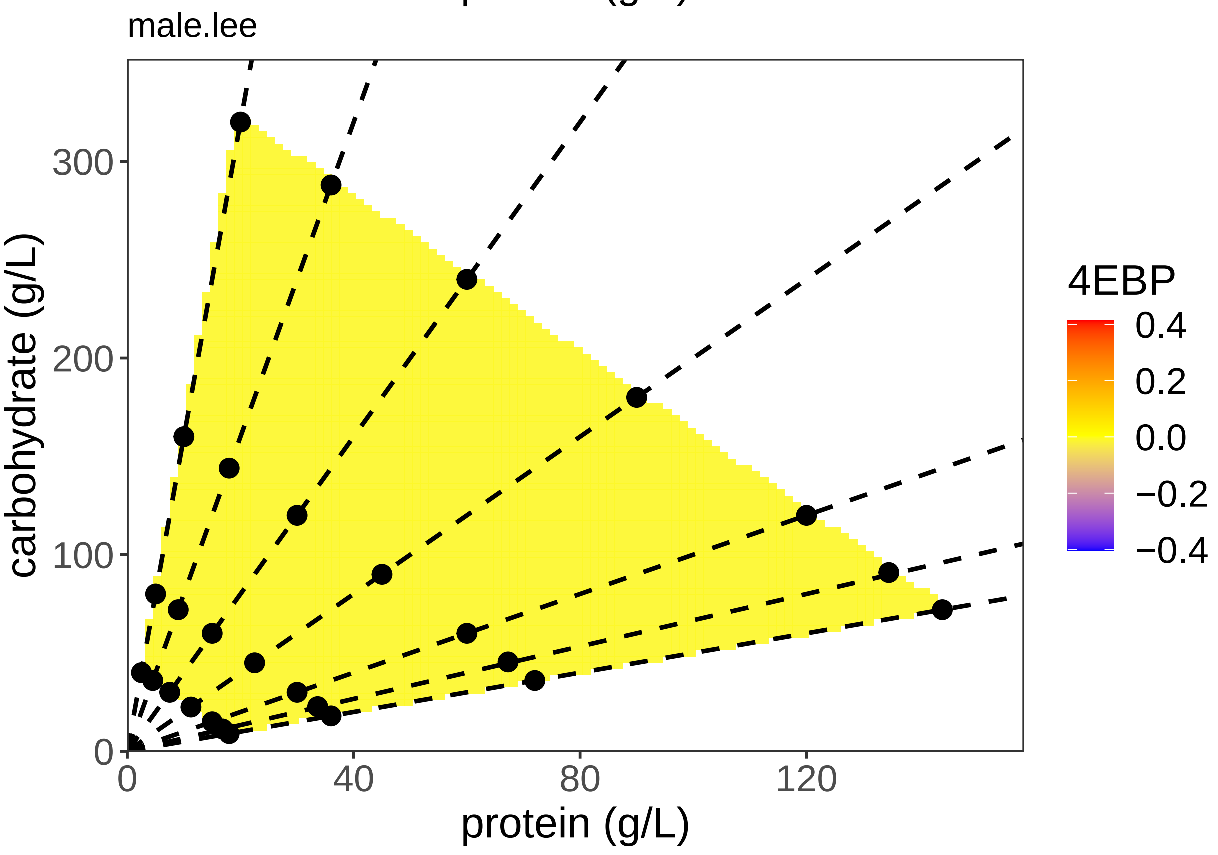
male.lee
carbohydrate (g/L)
protein (g/L)
0
100
200
300
0
40
80
120
4EBP
0.4
0.2
0.0
−0.2
−0.4

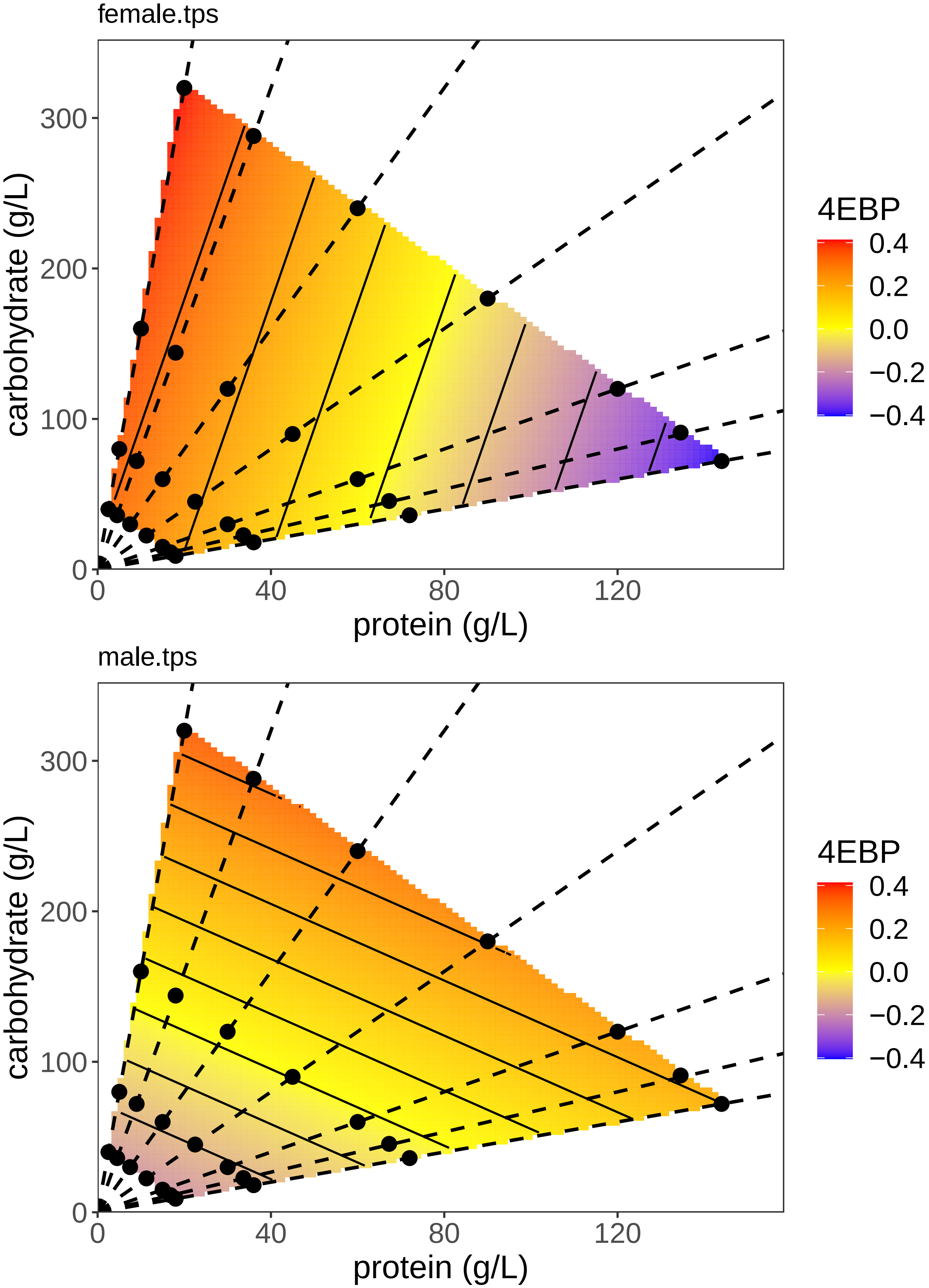


##Analysis of Ash2L

Test whether there is a difference between the sexes

```
Ash2LModel1 <- lm(Ash2L ~ sex + (carb:prot + poly(carb, 2) +
    poly(prot, 2)), data = all.data)
Ash2LModel2 <- lm(Ash2L ~ sex * (carb:prot + poly(carb, 2) +
    poly(prot, 2)), data = all.data)
anova(Ash2LModel1, Ash2LModel2)
```

```
## Analysis of Variance Table
##
## Model 1: Ash2L ~ sex + (carb:prot + poly(carb, 2) + poly(prot, 2))
## Model 2: Ash2L ~ sex * (carb:prot + poly(carb, 2) + poly(prot, 2))
##   Res.Df    RSS Df Sum of Sq      F   Pr(>F)
## 1    148 77.290
## 2    143 67.329  5    9.9602 4.2309 0.001287 **
## ---
## Signif. codes:  0 '***' 0.001 '**' 0.01 '*' 0.05 '.' 0.1 ' ' 1
```

There is a significant sex by diet interaction

Test the effect of C and P on Ash2L expression in males and females

```
Ash2L.m <- lm(Ash2L ~ (carb:prot + poly(carb, 2) + poly(prot,
    2)), data = m.data)
Ash2L.f <- lm(Ash2L ~ (carb:prot + poly(carb, 2) + poly(prot,
    2)), data = f.data)
summary(Ash2L.m)
```

```
##
## Call:
## lm(formula = Ash2L ~ (carb:prot + poly(carb, 2) + poly(prot,
##     2)), data = m.data)
##
## Residuals:
##      Min       1Q   Median       3Q      Max
## -3.07398 -0.32739  0.03009  0.32073  1.71666
##
## Coefficients:
##                   Estimate Std. Error t value Pr(>|t|)
## (Intercept)      3.515e-01  2.108e-01   1.668   0.0999 .
## poly(carb, 2)1   2.225e+00  1.312e+00   1.696   0.0944 .
## poly(carb, 2)2  -5.638e-01  7.523e-01  -0.749   0.4561
## poly(prot, 2)1   6.928e-01  1.606e+00   0.431   0.6675
## poly(prot, 2)2   4.282e-02  7.010e-01   0.061   0.9515
## carb:prot       -1.957e-05  4.748e-05  -0.412   0.6815
## ---
## Signif. codes:  0 '***' 0.001 '**' 0.01 '*' 0.05 '.' 0.1 ' ' 1
##
## Residual standard error: 0.6834 on 69 degrees of freedom
## Multiple R-squared:  0.09914,    Adjusted R-squared:  0.03386
## F-statistic: 1.519 on 5 and 69 DF,  p-value: 0.1954
```

```
summary(Ash2L.f)
```

```
##
## Call:
## lm(formula = Ash2L ~ (carb:prot + poly(carb, 2) + poly(prot,
```

```
##     2)), data = f.data)
##
## Residuals:
##      Min       1Q   Median       3Q      Max
## -2.43341 -0.29747  0.04111  0.35310  2.20723
##
## Coefficients:
##                   Estimate Std. Error t value Pr(>|t|)
## (Intercept)     -2.343e-01  1.982e-01  -1.182   0.2409
## poly(carb, 2)1 -2.857e+00  1.388e+00  -2.057   0.0432 *
## poly(carb, 2)2 -5.736e-01  7.920e-01  -0.724   0.4712
## poly(prot, 2)1 -3.175e+00  1.603e+00  -1.981   0.0513 .
## poly(prot, 2)2  1.451e+00  7.177e-01   2.022   0.0468 *
## carb:prot       4.564e-05  4.532e-05   1.007   0.3172
## ---
## Signif. codes:  0 '***' 0.001 '**' 0.01 '*' 0.05 '.' 0.1 ' ' 1
##
## Residual standard error: 0.6888 on 74 degrees of freedom
##   (4 observations deleted due to missingness)
## Multiple R-squared:  0.2077, Adjusted R-squared:  0.1542
## F-statistic: 3.881 on 5 and 74 DF,  p-value: 0.003533
```

There is an effect of protein in females but not in males. The quadratic is marginally significant in females.

Re-test the effect of P on expression in males and females using the simpler models

```r
# Non-Orthoganol Coef
summary(lm(Ash2L ~ (poly(carb, 1) + poly(prot, 2, raw = TRUE)),
    data = f.data))[4]
```

```
## $coefficients
##                               Estimate   Std. Error   t value   Pr(>|t|)
## (Intercept)                  0.3642920416 1.652260e-01  2.204810 0.03049249
## poly(carb, 1)               -1.6425786832 7.536358e-01 -2.179539 0.03238919
## poly(prot, 2, raw = TRUE)1 -0.0184075237 7.186085e-03 -2.561551 0.01240092
## poly(prot, 2, raw = TRUE)2  0.0001052562 5.070288e-05  2.075942 0.04128150
```

```r
# Orthoganol Test
Ash2L.m <- lm(Ash2L ~ (poly(carb, 1)), data = m.data)
Ash2L.f <- lm(Ash2L ~ (poly(carb, 1) + poly(prot, 2)), data = f.data)
summary(Ash2L.m)
```

```
##
## Call:
## lm(formula = Ash2L ~ (poly(carb, 1)), data = m.data)
##
## Residuals:
##      Min       1Q   Median       3Q      Max
## -3.16227 -0.34539  0.03132  0.34251  1.65940
##
## Coefficients:
##               Estimate Std. Error t value Pr(>|t|)
## (Intercept)     0.2710     0.0772   3.510 0.000772 ***
## poly(carb, 1)   1.7724     0.6686   2.651 0.009833 **
## ---
## Signif. codes:  0 '***' 0.001 '**' 0.01 '*' 0.05 '.' 0.1 ' ' 1
##
```

```
## Residual standard error: 0.6686 on 73 degrees of freedom
## Multiple R-squared:  0.08782,    Adjusted R-squared:  0.07533
## F-statistic: 7.028 on 1 and 73 DF,  p-value: 0.009833
```

```r
summary(Ash2L.f)
```

```
##
## Call:
## lm(formula = Ash2L ~ (poly(carb, 1) + poly(prot, 2)), data = f.data)
##
## Residuals:
##      Min       1Q   Median       3Q      Max
## -2.39590 -0.25368  0.02828  0.38388  2.19680
##
## Coefficients:
##                 Estimate Std. Error t value Pr(>|t|)
## (Intercept)     -0.04822    0.07694  -0.627   0.5327
## poly(carb, 1)   -1.64258    0.75364  -2.180   0.0324 *
## poly(prot, 2)1 -1.52209    0.70469  -2.160   0.0339 *
## poly(prot, 2)2  1.47317    0.70964   2.076   0.0413 *
## ---
## Signif. codes:  0 '***' 0.001 '**' 0.01 '*' 0.05 '.' 0.1 ' ' 1
##
## Residual standard error: 0.6876 on 76 degrees of freedom
##   (4 observations deleted due to missingness)
## Multiple R-squared:  0.1891, Adjusted R-squared:  0.1571
## F-statistic: 5.907 on 3 and 76 DF,  p-value: 0.001118
```

There is a carb effect in both males and female and a quadratic protein effect in females

Re-test the effect of sex using the simplest model

```r
Ash2LModel1 <- lm(Ash2L ~ sex * (poly(carb, 1) + poly(prot, 2)),
    data = all.data)
Ash2LModel1_5 <- lm(Ash2L ~ sex * (carb), data = all.data)
Ash2LModel2 <- lm(Ash2L ~ sex + (carb), data = all.data)
anova(Ash2LModel2, Ash2LModel1_5, Ash2LModel1)
```

```
## Analysis of Variance Table
##
## Model 1: Ash2L ~ sex + (carb)
## Model 2: Ash2L ~ sex * (carb)
## Model 3: Ash2L ~ sex * (poly(carb, 1) + poly(prot, 2))
##   Res.Df    RSS Df Sum of Sq       F    Pr(>F)
## 1    152 80.009
## 2    151 72.610  1    7.3989 15.8875 0.0001055 ***
## 3    147 68.459  4    4.1514  2.2286 0.0686905 .
## ---
## Signif. codes:  0 '***' 0.001 '**' 0.01 '*' 0.05 '.' 0.1 ' ' 1
```

```r
summary(Ash2LModel1_5)
```

```
##
## Call:
## lm(formula = Ash2L ~ sex * (carb), data = all.data)
##
## Residuals:
```

```
##      Min       1Q   Median       3Q      Max
## -3.16227 -0.36458  0.04368  0.39592  2.04856
##
## Coefficients:
##               Estimate Std. Error t value Pr(>|t|)
## (Intercept)  0.2019914  0.1149528   1.757 0.080916 .
## sexM        -0.1527463  0.1647749  -0.927 0.355406
## carb        -0.0029432  0.0009809  -3.000 0.003154 **
## sexM:carb    0.0055443  0.0014134   3.923 0.000133 ***
## ---
## Signif. codes:  0 '***' 0.001 '**' 0.01 '*' 0.05 '.' 0.1 ' ' 1
##
## Residual standard error: 0.6934 on 151 degrees of freedom
##   (4 observations deleted due to missingness)
## Multiple R-squared:  0.137,  Adjusted R-squared:  0.1198
## F-statistic: 7.989 on 3 and 151 DF,  p-value: 5.6e-05
```

There is a significiant difference in response between males and females, but it lies in the response to carbohydrates

Plot the model and the TPS

```
## Warning:
## Grid searches over lambda (nugget and sill variances) with  minima at the endpoints:
##   (GCV) Generalized Cross-Validation
##    minimum at  right endpoint  lambda  =  268.0589 (eff. df= 3.00101 )
```

```
## Warning:
## Grid searches over lambda (nugget and sill variances) with  minima at the endpoints:
##   (GCV) Generalized Cross-Validation
##    minimum at  right endpoint  lambda  =  253.2424 (eff. df= 3.000991 )
```

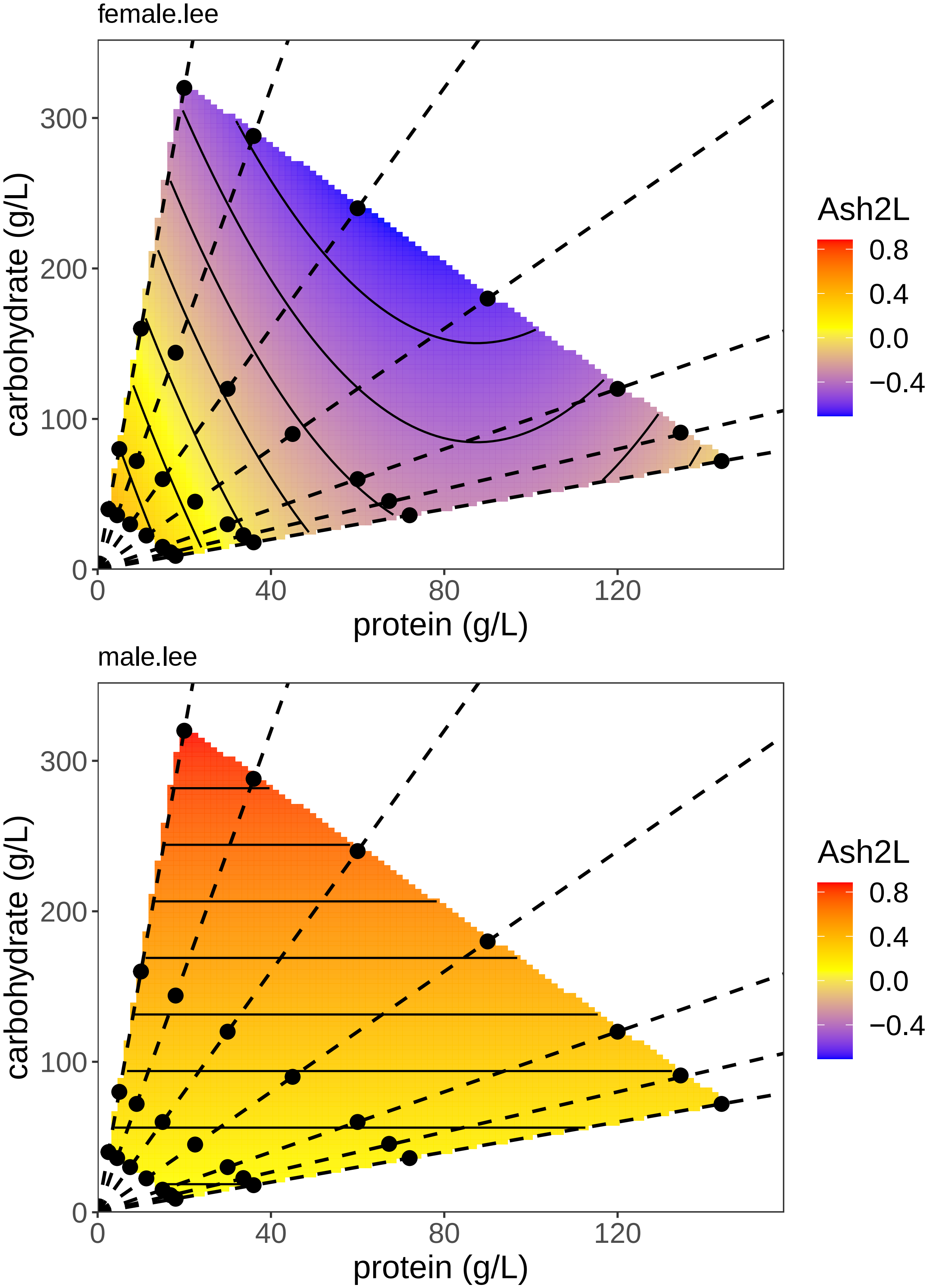
female.lee
carbohydrate (g/L)
protein (g/L)
0
100
200
300
0
40
80
120
Ash2L
0.8
0.4
0.0
−0.4
male.lee
carbohydrate (g/L)
protein (g/L)
0
100
200
300
0
40
80
120
Ash2L
0.8
0.4
0.0
−0.4

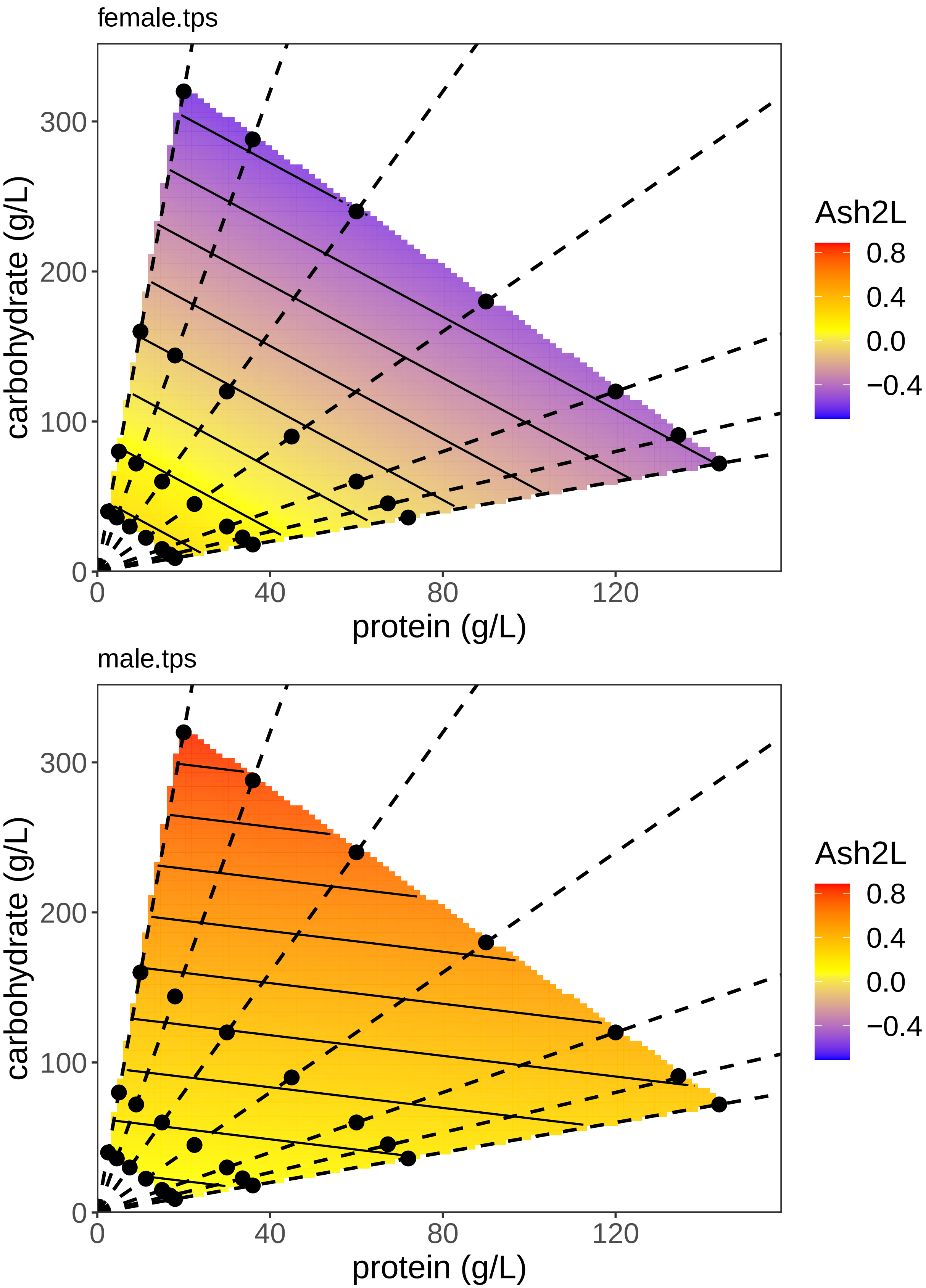

female.tps
carbohydrate (g/L)
protein (g/L)
Ash2L
0.8
0.4
0.0
−0.4
300
200
100
0
0
40
80
120
male.tps
carbohydrate (g/L)
protein (g/L)
Ash2L
0.8
0.4
0.0
−0.4
300
200
100
0
0
40
80
120

#### Analysis of CG3071

Test whether there is a difference between the sexes

```
CG3071Model1 <- lm(CG3071 ~ sex + (carb:prot + poly(carb, 2) +
    poly(prot, 2)), data = all.data)
CG3071Model2 <- lm(CG3071 ~ sex * (carb:prot + poly(carb, 2) +
    poly(prot, 2)), data = all.data)
anova(CG3071Model1, CG3071Model2)
```

```
## Analysis of Variance Table
##
## Model 1: CG3071 ~ sex + (carb:prot + poly(carb, 2) + poly(prot, 2))
## Model 2: CG3071 ~ sex * (carb:prot + poly(carb, 2) + poly(prot, 2))
##   Res.Df    RSS Df Sum of Sq     F    Pr(>F)
## 1    123 52.497
## 2    118 43.322  5    9.1747 4.998 0.0003452 ***
## ---
## Signif. codes:  0 '***' 0.001 '**' 0.01 '*' 0.05 '.' 0.1 ' ' 1
```

```
summary(CG3071Model2)
```

```
##
## Call:
## lm(formula = CG3071 ~ sex * (carb:prot + poly(carb, 2) + poly(prot,
##     2)), data = all.data)
##
## Residuals:
##      Min       1Q   Median       3Q      Max
## -1.75495 -0.34503  0.04439  0.33764  1.68731
##
## Coefficients:
##                        Estimate Std. Error t value Pr(>|t|)
## (Intercept)          -9.634e-01  1.794e-01  -5.370 4.00e-07 ***
## sexM                 -5.207e-02  2.827e-01  -0.184  0.85420
## poly(carb, 2)1       -7.502e+00  1.729e+00  -4.338 3.05e-05 ***
## poly(carb, 2)2        2.090e+00  9.909e-01   2.109  0.03702 *
## poly(prot, 2)1       -1.053e+01  2.049e+00  -5.137 1.12e-06 ***
## poly(prot, 2)2        3.880e+00  8.885e-01   4.367 2.72e-05 ***
## carb:prot             1.339e-04  4.165e-05   3.215  0.00169 **
## sexM:poly(carb, 2)1  7.328e+00  2.590e+00   2.830  0.00547 **
## sexM:poly(carb, 2)2 -7.139e-01  1.445e+00  -0.494  0.62223
## sexM:poly(prot, 2)1  8.925e+00  3.179e+00   2.808  0.00584 **
## sexM:poly(prot, 2)2 -2.691e+00  1.410e+00  -1.909  0.05875 .
## sexM:carb:prot      -1.162e-04  6.628e-05  -1.753  0.08228 .
## ---
## Signif. codes:  0 '***' 0.001 '**' 0.01 '*' 0.05 '.' 0.1 ' ' 1
##
## Residual standard error: 0.6059 on 118 degrees of freedom
##   (29 observations deleted due to missingness)
## Multiple R-squared:  0.4652, Adjusted R-squared:  0.4153
## F-statistic:  9.33 on 11 and 118 DF,  p-value: 7.009e-12
```

There is a sex*diet interaction. It's complex, but males have (generally) a lower expression level, although you can really interpret this because of the higher order interactions.

Test the effect of C and P on CG3071 expression in males and females

```r
# Non-orthoganol Coef
summary(lm(CG3071 ~ (carb:prot + poly(carb, 2, raw = TRUE) +
    poly(prot, 2, raw = TRUE)), data = f.data))[4]
```

```
## $coefficients
##                                 Estimate   Std. Error   t value     Pr(>|t|)
## (Intercept)                 1.296450e+00 2.432483e-01  5.329737 1.242480e-06
## poly(carb, 2, raw = TRUE)1 -1.413151e-02 3.809595e-03 -3.709452 4.240074e-04
## poly(carb, 2, raw = TRUE)2  2.303095e-05 1.105597e-05  2.083124 4.105921e-02
## poly(prot, 2, raw = TRUE)1 -4.856657e-02 7.824247e-03 -6.207188 3.857029e-08
## poly(prot, 2, raw = TRUE)2  2.017339e-04 4.677728e-05  4.312648 5.423138e-05
## carb:prot                   1.338788e-04 4.217265e-05  3.174542 2.267692e-03
```

```r
# Orthoganol Test
CG3071.m <- lm(CG3071 ~ (carb:prot + poly(carb, 2) + poly(prot,
    2)), data = m.data)
CG3071.f <- lm(CG3071 ~ (carb:prot + poly(carb, 2) + poly(prot,
    2)), data = f.data)
summary(CG3071.m)
```

```
##
## Call:
## lm(formula = CG3071 ~ (carb:prot + poly(carb, 2) + poly(prot,
##     2)), data = m.data)
##
## Residuals:
##      Min       1Q   Median       3Q      Max
## -1.57284 -0.34292  0.02048  0.35816  1.29044
##
## Coefficients:
##                   Estimate Std. Error t value Pr(>|t|)
## (Intercept)     -1.022e+00  2.182e-01  -4.685 2.11e-05 ***
## poly(carb, 2)1 -1.539e-01  1.278e+00  -0.120    0.905
## poly(carb, 2)2  9.373e-01  7.044e-01   1.331    0.189
## poly(prot, 2)1 -1.078e+00  1.660e+00  -0.650    0.519
## poly(prot, 2)2  8.143e-01  7.367e-01   1.105    0.274
## carb:prot       1.772e-05  5.069e-05   0.350    0.728
## ---
## Signif. codes:  0 '***' 0.001 '**' 0.01 '*' 0.05 '.' 0.1 ' ' 1
##
## Residual standard error: 0.5957 on 51 degrees of freedom
##   (18 observations deleted due to missingness)
## Multiple R-squared:  0.07303,    Adjusted R-squared:  -0.01785
## F-statistic: 0.8036 on 5 and 51 DF,  p-value: 0.5524
```

```r
summary(CG3071.f)
```

```
##
## Call:
## lm(formula = CG3071 ~ (carb:prot + poly(carb, 2) + poly(prot,
##     2)), data = f.data)
##
## Residuals:
##      Min       1Q   Median       3Q      Max
```

```
## -1.75495 -0.35676  0.04507  0.29742  1.68731
##
## Coefficients:
##                   Estimate Std. Error t value Pr(>|t|)
## (Intercept)     -9.414e-01  1.793e-01  -5.251 1.68e-06 ***
## poly(carb, 2)1 -5.498e+00  1.293e+00  -4.252 6.72e-05 ***
## poly(carb, 2)2  1.528e+00  7.337e-01   2.083  0.04106 *
## poly(prot, 2)1 -7.668e+00  1.493e+00  -5.137 2.60e-06 ***
## poly(prot, 2)2  2.823e+00  6.547e-01   4.313 5.42e-05 ***
## carb:prot       1.339e-04  4.217e-05   3.175  0.00227 **
## ---
## Signif. codes:  0 '***' 0.001 '**' 0.01 '*' 0.05 '.' 0.1 ' ' 1
##
## Residual standard error: 0.6136 on 67 degrees of freedom
##   (11 observations deleted due to missingness)
## Multiple R-squared:  0.5406, Adjusted R-squared:  0.5064
## F-statistic: 15.77 on 5 and 67 DF,  p-value: 3.055e-10
```

There is an effect of protein and carbohydrates in females but not in males

Plot the model and the TPS

```
## Warning:
## Grid searches over lambda (nugget and sill variances) with  minima at the endpoints:
##   (GCV) Generalized Cross-Validation
##    minimum at  right endpoint  lambda  =  182.9905 (eff. df= 3.001008 )
```

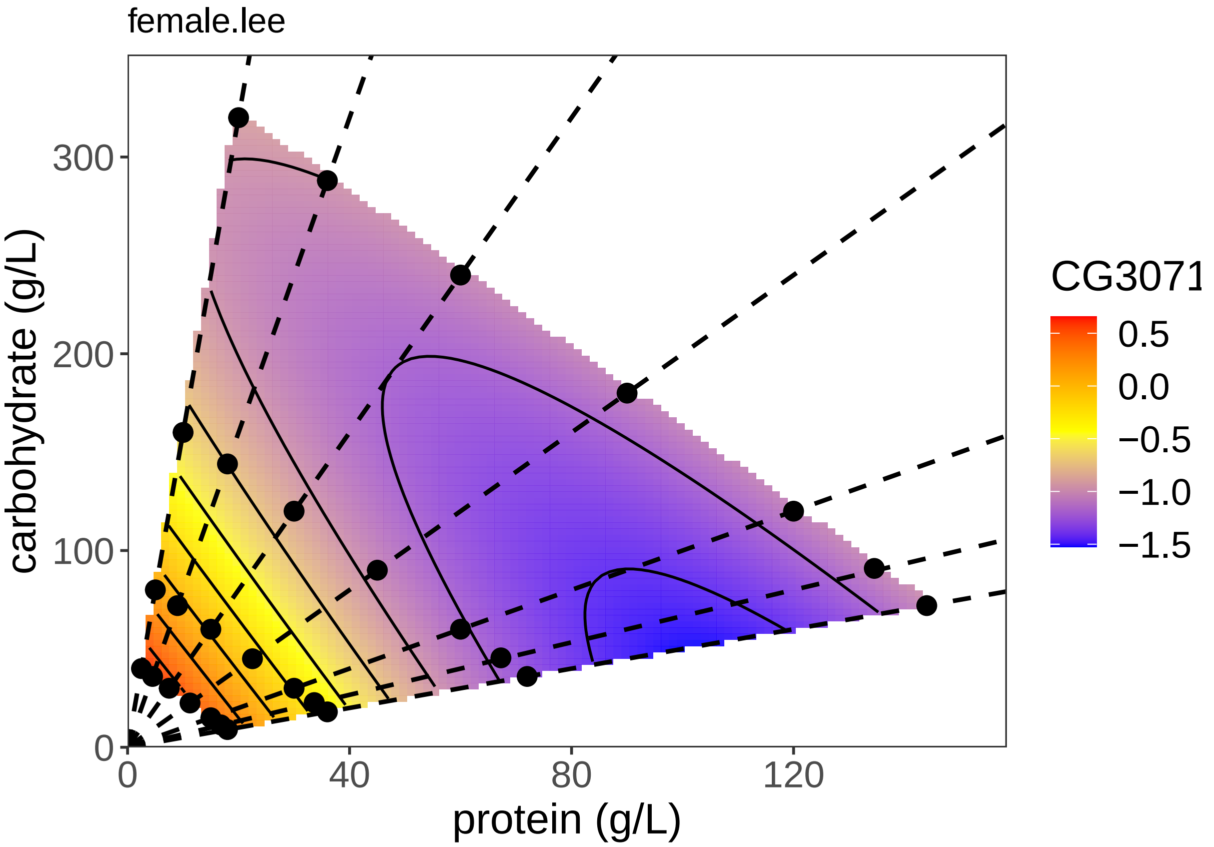

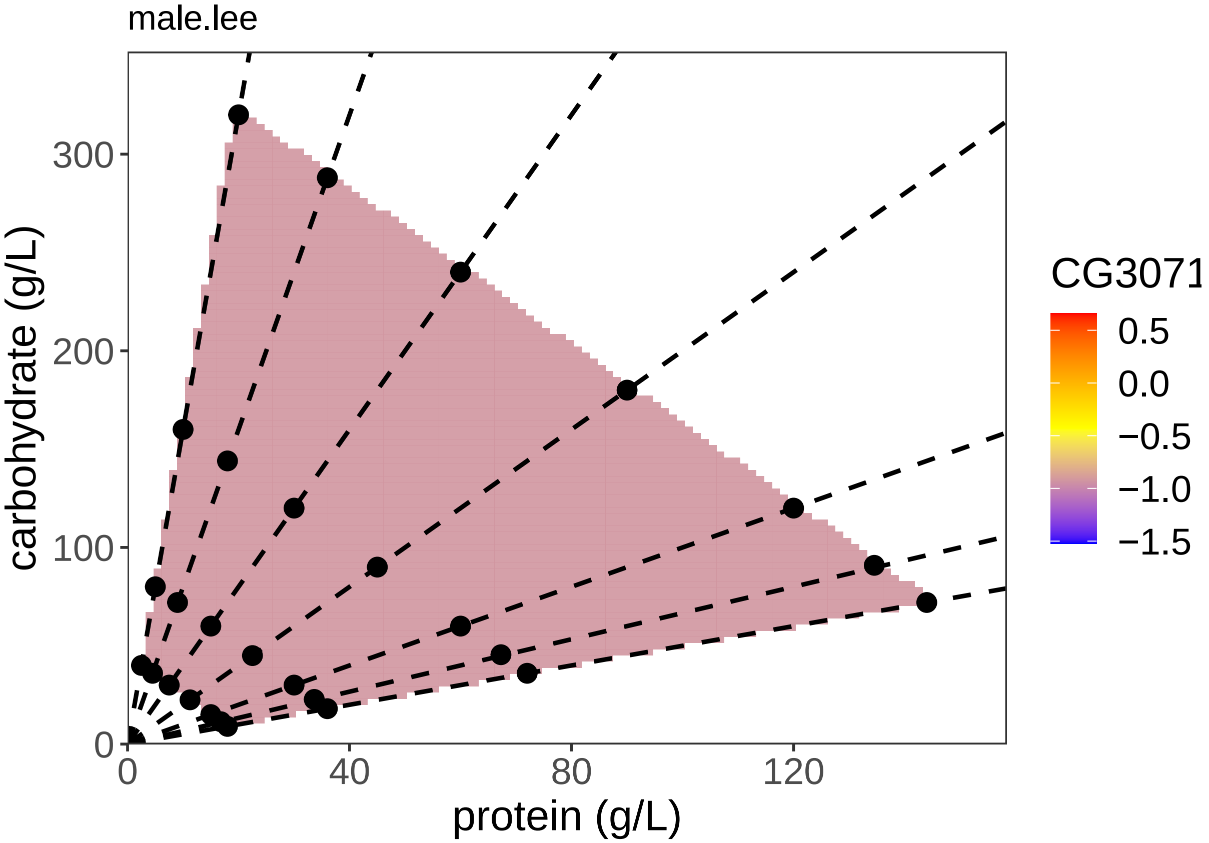
male.lee
carbohydrate (g/L)
protein (g/L)
0
100
200
300
0
40
80
120
CG3071
0.5
0.0
−0.5
−1.0
−1.5

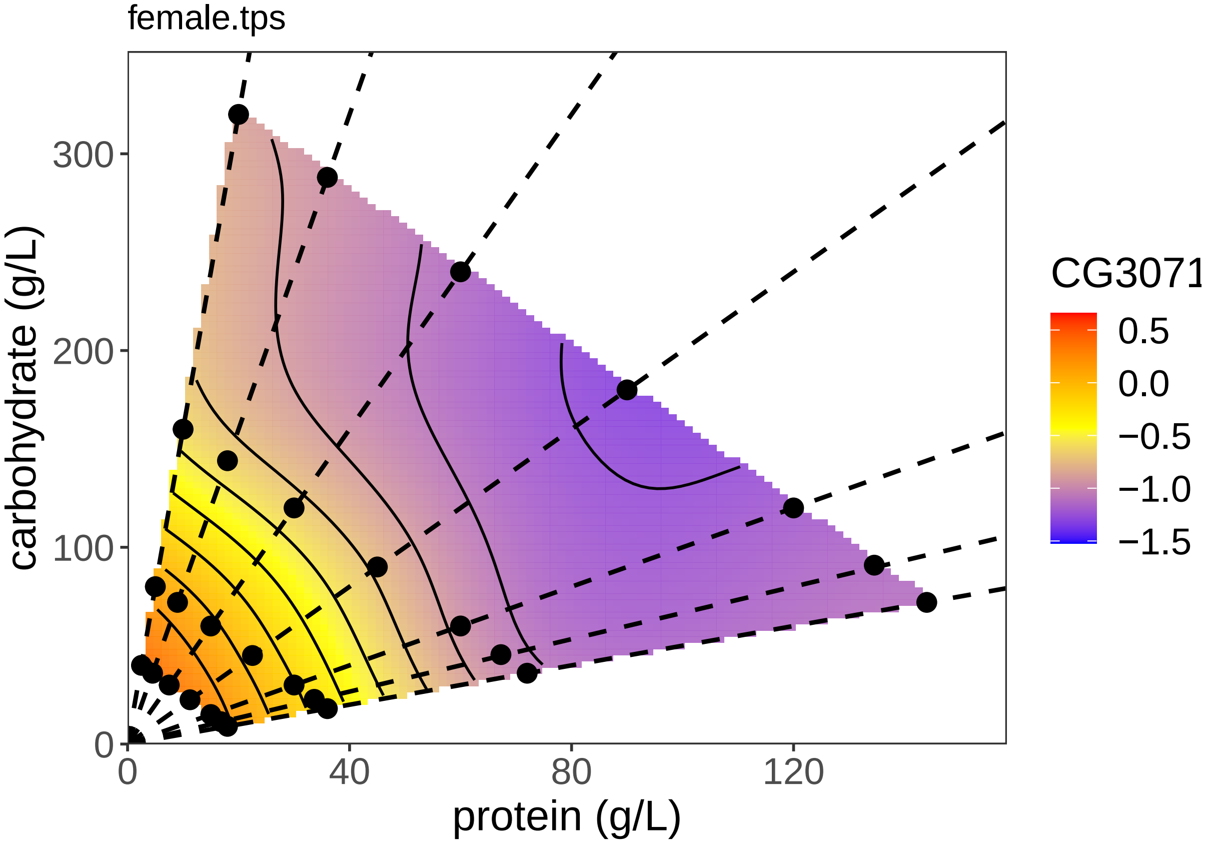
female.tps
carbohydrate (g/L)
protein (g/L)
0
100
200
300
0
40
80
120
CG3071
0.5
0.0
−0.5
−1.0
−1.5

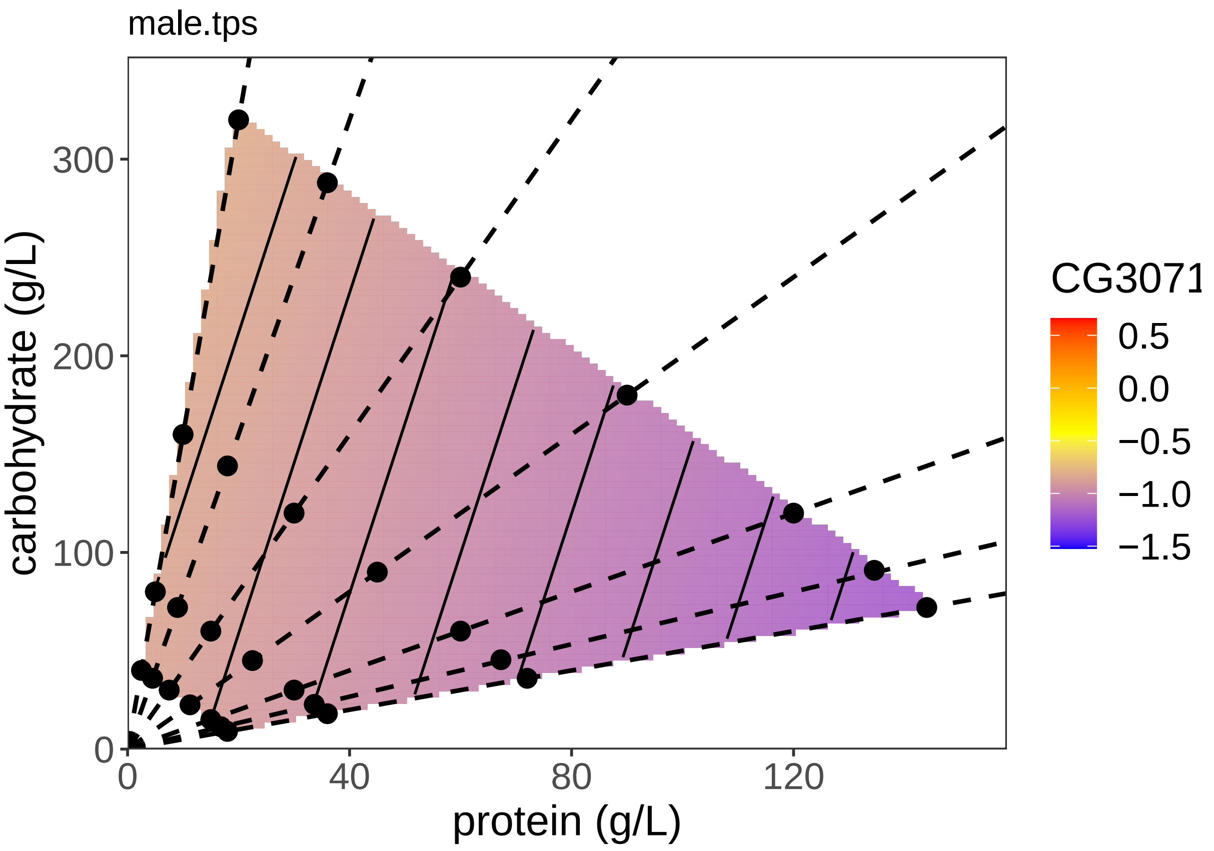


##### Analysis of dILP2

Test whether there is a difference between the sexes

```
dILP2Model1 <- lm(dILP2 ~ sex + (carb:prot + poly(carb, 2) +
    poly(prot, 1)), data = all.data)
dILP2Model2 <- lm(dILP2 ~ sex * (carb:prot + poly(carb, 2) +
    poly(prot, 1)), data = all.data)
anova(dILP2Model1, dILP2Model2)
```

```
## Analysis of Variance Table
##
## Model 1: dILP2 ~ sex + (carb:prot + poly(carb, 2) + poly(prot, 1))
## Model 2: dILP2 ~ sex * (carb:prot + poly(carb, 2) + poly(prot, 1))
##   Res.Df    RSS Df Sum of Sq      F Pr(>F)
## 1    121 150.84
## 2    117 147.46  4    3.3861 0.6717 0.6129
```

```
summary(dILP2Model1)
```

```
##
## Call:
## lm(formula = dILP2 ~ sex + (carb:prot + poly(carb, 2) + poly(prot,
##     1)), data = all.data)
##
## Residuals:
##     Min      1Q  Median      3Q     Max
```

```
## -2.7006 -0.5581  0.0247  0.5956  3.5333
##
## Coefficients:
##                   Estimate Std. Error t value Pr(>|t|)
## (Intercept)     -1.581e+00  2.679e-01  -5.902 3.36e-08 ***
## sexM            -2.227e-01  2.020e-01  -1.103   0.2724
## poly(carb, 2)1 -6.285e+00  2.417e+00  -2.600   0.0105 *
## poly(carb, 2)2  2.196e+00  1.378e+00   1.593   0.1137
## poly(prot, 1)  -6.729e+00  2.934e+00  -2.293   0.0236 *
## carb:prot       1.237e-04  5.829e-05   2.123   0.0358 *
## ---
## Signif. codes:  0 '***' 0.001 '**' 0.01 '*' 0.05 '.' 0.1 ' ' 1
##
## Residual standard error: 1.117 on 121 degrees of freedom
##   (32 observations deleted due to missingness)
## Multiple R-squared:  0.09108,    Adjusted R-squared:  0.05353
## F-statistic: 2.425 on 5 and 121 DF,  p-value: 0.03918
```

There is no sex*diet interaction, and males and females have the same expression level

Test the effect of C and P on dILP2 expression in males and females.

```
dILP2.m <- lm(dILP2 ~ (carb:prot + poly(carb, 2) + poly(prot,
    2)), data = m.data)
dILP2.f <- lm(dILP2 ~ (carb:prot + poly(carb, 2) + poly(prot,
    2)), data = f.data)
summary(dILP2.m)
```

```
##
## Call:
## lm(formula = dILP2 ~ (carb:prot + poly(carb, 2) + poly(prot,
##     2)), data = m.data)
##
## Residuals:
##     Min      1Q  Median      3Q     Max
## -2.7089 -1.1599 -0.0823  0.9700  3.6180
##
## Coefficients:
##                   Estimate Std. Error t value Pr(>|t|)
## (Intercept)     -1.8138671  0.6003913  -3.021  0.00407 **
## poly(carb, 2)1 -5.0772801  3.5113745  -1.446  0.15483
## poly(carb, 2)2  0.3515902  2.0640360   0.170  0.86547
## poly(prot, 2)1 -4.0002752  4.5763875  -0.874  0.38650
## poly(prot, 2)2 -0.0239460  2.0582487  -0.012  0.99077
## carb:prot       0.0001261  0.0001286   0.981  0.33161
## ---
## Signif. codes:  0 '***' 0.001 '**' 0.01 '*' 0.05 '.' 0.1 ' ' 1
##
## Residual standard error: 1.552 on 47 degrees of freedom
##   (22 observations deleted due to missingness)
## Multiple R-squared:  0.04861,    Adjusted R-squared:  -0.0526
## F-statistic: 0.4803 on 5 and 47 DF,  p-value: 0.7891
```

```
summary(dILP2.f)
```

```
##
```

```
## Call:
## lm(formula = dILP2 ~ (carb:prot + poly(carb, 2) + poly(prot,
##     2)), data = f.data)
##
## Residuals:
##     Min      1Q  Median      3Q     Max
## -2.3927 -0.3349  0.0405  0.3937  1.5137
##
## Coefficients:
##                   Estimate Std. Error t value Pr(>|t|)
## (Intercept)    -1.576e+00  2.032e-01  -7.759 5.99e-11 ***
## poly(carb, 2)1 -3.983e+00  1.457e+00  -2.734  0.00796 **
## poly(carb, 2)2  2.708e+00  8.286e-01   3.268  0.00170 **
## poly(prot, 2)1 -5.649e+00  1.690e+00  -3.343  0.00135 **
## poly(prot, 2)2  1.297e+00  7.537e-01   1.721  0.08976 .
## carb:prot       1.302e-04  4.725e-05   2.756  0.00750 **
## ---
## Signif. codes:  0 '***' 0.001 '**' 0.01 '*' 0.05 '.' 0.1 ' ' 1
##
## Residual standard error: 0.6946 on 68 degrees of freedom
##   (10 observations deleted due to missingness)
## Multiple R-squared:  0.2901, Adjusted R-squared:  0.2379
## F-statistic: 5.557 on 5 and 68 DF,  p-value: 0.0002381
```

There is no effect in males but there is an effect in females, although the quadratic effect in females is not significant.

Retest with simpler

```r
# Non-orthoganol Ceof
summary(lm(dILP2 ~ (carb:prot + poly(carb, 2, raw = TRUE) + poly(prot,
    1)), data = f.data))[4]
```

```
## $coefficients
##                                Estimate   Std. Error   t value     Pr(>|t|)
## (Intercept)                -6.434777e-01 1.957381e-01 -3.287442 0.0015924495
## poly(carb, 2, raw = TRUE)1 -1.644825e-02 4.411233e-03 -3.728719 0.0003906888
## poly(carb, 2, raw = TRUE)2  3.777346e-05 1.253535e-05  3.013355 0.0036112792
## poly(prot, 1)              -5.254696e+00 1.697643e+00 -3.095288 0.0028402027
## carb:prot                   1.199810e-04 4.753105e-05  2.524266 0.0138986254
```

```r
# Orthoganol Test
dILP2.m <- lm(dILP2 ~ (carb:prot + poly(carb, 2) + poly(prot,
    1)), data = m.data)
dILP2.f <- lm(dILP2 ~ (carb:prot + poly(carb, 2) + poly(prot,
    1)), data = f.data)
summary(dILP2.m)
```

```
##
## Call:
## lm(formula = dILP2 ~ (carb:prot + poly(carb, 2) + poly(prot,
##     1)), data = m.data)
##
## Residuals:
##     Min      1Q  Median      3Q     Max
## -2.7113 -1.1590 -0.0851  0.9701  3.6165
##
```

```
## Coefficients:
##                   Estimate Std. Error t value Pr(>|t|)
## (Intercept)     -1.8145327  0.5914019  -3.068  0.00354 **
## poly(carb, 2)1 -5.0747714  3.4680525  -1.463  0.14991
## poly(carb, 2)2  0.3558625  2.0098399   0.177  0.86021
## poly(prot, 1)  -4.0010091  4.5280422  -0.884  0.38131
## carb:prot       0.0001262  0.0001268   0.995  0.32462
## ---
## Signif. codes:  0 '***' 0.001 '**' 0.01 '*' 0.05 '.' 0.1 ' ' 1
##
## Residual standard error: 1.536 on 48 degrees of freedom
##   (22 observations deleted due to missingness)
## Multiple R-squared:  0.04861,    Adjusted R-squared:  -0.03067
## F-statistic: 0.6131 on 4 and 48 DF,  p-value: 0.6552
```

```r
summary(dILP2.f)
```

```
##
## Call:
## lm(formula = dILP2 ~ (carb:prot + poly(carb, 2) + poly(prot,
##     1)), data = f.data)
##
## Residuals:
##      Min       1Q   Median       3Q      Max
## -2.19688 -0.34517  0.00475  0.44653  1.57994
##
## Coefficients:
##                   Estimate Std. Error t value Pr(>|t|)
## (Intercept)     -1.544e+00  2.052e-01  -7.527 1.46e-10 ***
## poly(carb, 2)1 -3.978e+00  1.477e+00  -2.692  0.00889 **
## poly(carb, 2)2  2.507e+00  8.319e-01   3.013  0.00361 **
## poly(prot, 1)  -5.255e+00  1.698e+00  -3.095  0.00284 **
## carb:prot       1.200e-04  4.753e-05   2.524  0.01390 *
## ---
## Signif. codes:  0 '***' 0.001 '**' 0.01 '*' 0.05 '.' 0.1 ' ' 1
##
## Residual standard error: 0.7044 on 69 degrees of freedom
##   (10 observations deleted due to missingness)
## Multiple R-squared:  0.2591, Adjusted R-squared:  0.2162
## F-statistic: 6.034 on 4 and 69 DF,  p-value: 0.0003186
```

Re-testing sex interaction with the simpler model

```r
dILP2Model0 <- lm(dILP2 ~ (carb:prot + poly(carb, 2) + poly(prot,
    1)), data = all.data)
dILP2Model1 <- lm(dILP2 ~ sex + (carb:prot + poly(carb, 2) +
    poly(prot, 1)), data = all.data)
dILP2Model2 <- lm(dILP2 ~ sex * (carb:prot + poly(carb, 2) +
    poly(prot, 1)), data = all.data)
anova(dILP2Model0, dILP2Model1, dILP2Model2)
```

```
## Analysis of Variance Table
##
## Model 1: dILP2 ~ (carb:prot + poly(carb, 2) + poly(prot, 1))
## Model 2: dILP2 ~ sex + (carb:prot + poly(carb, 2) + poly(prot, 1))
## Model 3: dILP2 ~ sex * (carb:prot + poly(carb, 2) + poly(prot, 1))
```

```
##   Res.Df    RSS Df Sum of Sq      F Pr(>F)
## 1    122 152.36
## 2    121 150.84  1    1.5157 1.2026 0.2750
## 3    117 147.46  4    3.3861 0.6717 0.6129
```

```r
summary(anova(dILP2Model1, dILP2Model2))
```

```
##      Res.Df          RSS               Df      Sum of Sq           F
##  Min.   :117   Min.   :147.5   Min.   :4   Min.   :3.386   Min.   :0.6717
##  1st Qu.:118   1st Qu.:148.3   1st Qu.:4   1st Qu.:3.386   1st Qu.:0.6717
##  Median :119   Median :149.2   Median :4   Median :3.386   Median :0.6717
##  Mean   :119   Mean   :149.2   Mean   :4   Mean   :3.386   Mean   :0.6717
##  3rd Qu.:120   3rd Qu.:150.0   3rd Qu.:4   3rd Qu.:3.386   3rd Qu.:0.6717
##  Max.   :121   Max.   :150.8   Max.   :4   Max.   :3.386   Max.   :0.6717
##                                NA's   :1   NA's   :1       NA's   :1
##      Pr(>F)
##  Min.   :0.6129
##  1st Qu.:0.6129
##  Median :0.6129
##  Mean   :0.6129
##  3rd Qu.:0.6129
##  Max.   :0.6129
##  NA's   :1
```

```r
summary(dILP2Model1)
```

```
##
## Call:
## lm(formula = dILP2 ~ sex + (carb:prot + poly(carb, 2) + poly(prot,
##     1)), data = all.data)
##
## Residuals:
##     Min      1Q  Median      3Q     Max
## -2.7006 -0.5581  0.0247  0.5956  3.5333
##
## Coefficients:
##                   Estimate Std. Error t value Pr(>|t|)
## (Intercept)    -1.581e+00  2.679e-01  -5.902 3.36e-08 ***
## sexM           -2.227e-01  2.020e-01  -1.103   0.2724
## poly(carb, 2)1 -6.285e+00  2.417e+00  -2.600   0.0105 *
## poly(carb, 2)2  2.196e+00  1.378e+00   1.593   0.1137
## poly(prot, 1)  -6.729e+00  2.934e+00  -2.293   0.0236 *
## carb:prot       1.237e-04  5.829e-05   2.123   0.0358 *
## ---
## Signif. codes:  0 '***' 0.001 '**' 0.01 '*' 0.05 '.' 0.1 ' ' 1
##
## Residual standard error: 1.117 on 121 degrees of freedom
##   (32 observations deleted due to missingness)
## Multiple R-squared:  0.09108,    Adjusted R-squared:  0.05353
## F-statistic: 2.425 on 5 and 121 DF,  p-value: 0.03918
```

We can test for the power to detect a sex-by-diet interaction

```r
RsqFull <- summary(dILP2Model2)$r.squared
RsqRed <- summary(dILP2Model1)$r.squared
```

```r
fsq <- (RsqFull - RsqRed)/(1 - RsqFull)

predFull <- length(dILP2Model2$coefficients)
predRed <- length(dILP2Model1$coefficients)


wp.regression(n = nobs(dILP2Model2), p1 = predFull, p2 = predRed,
    f2 = fsq)
```

```
## Power for multiple regression
##
##       n p1 p2         f2 alpha     power
##     127 10  6 0.02296285  0.05 0.2195252
##
## URL: http://psychstat.org/regression
```

Plot the model and the TPS

```
## Warning:
## Grid searches over lambda (nugget and sill variances) with  minima at the endpoints:
##   (GCV) Generalized Cross-Validation
##    minimum at  right endpoint  lambda  =  158.9578 (eff. df= 3.001043 )
```

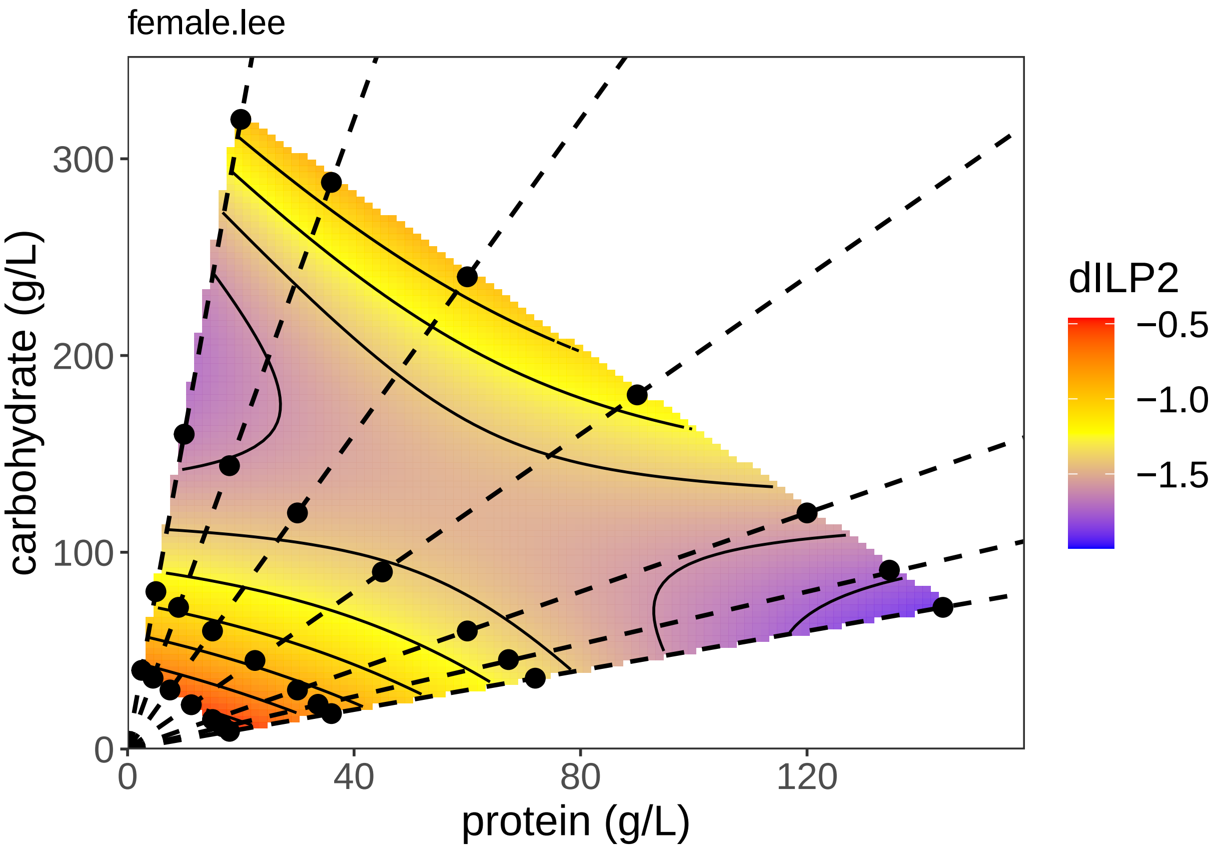

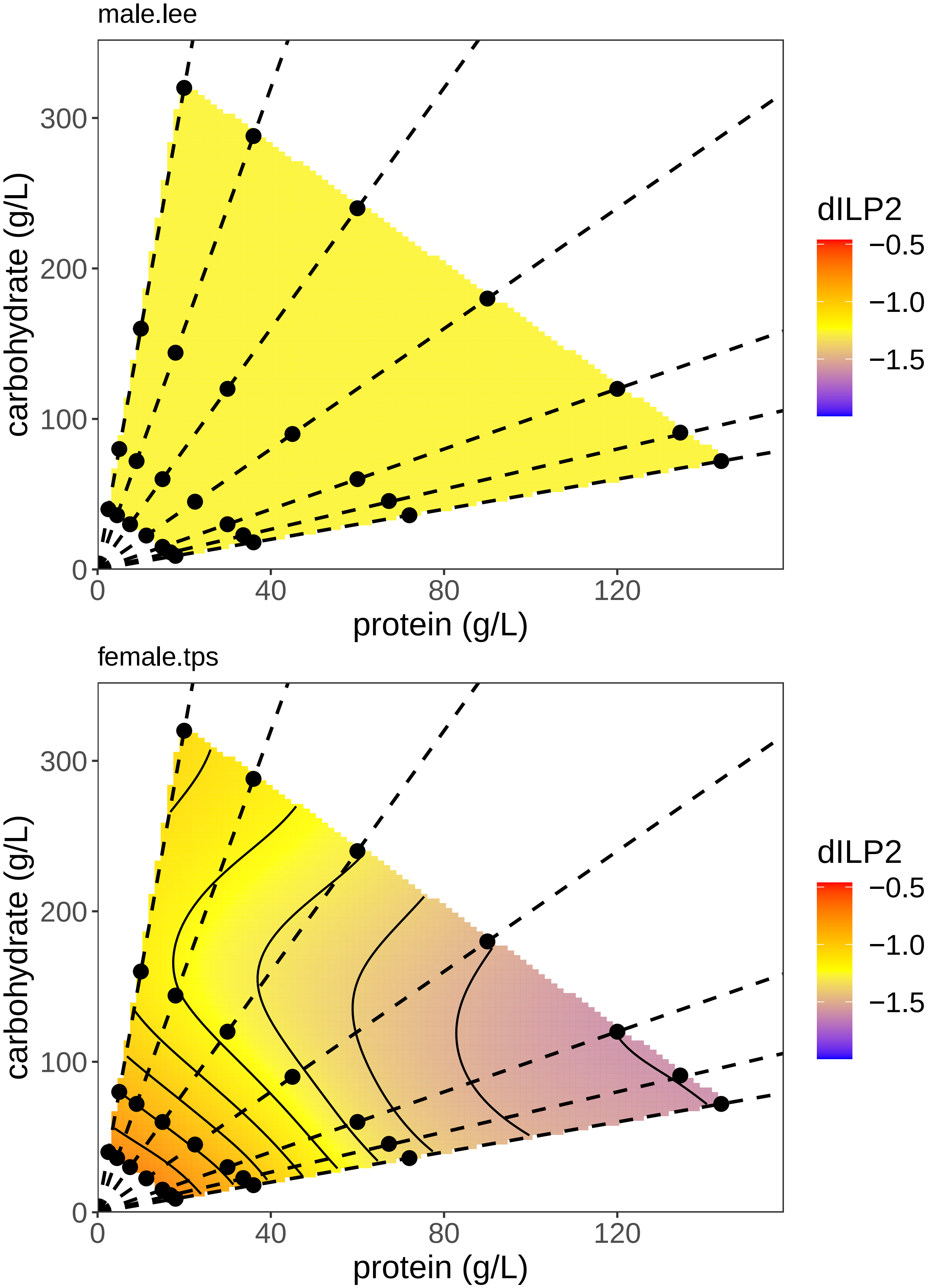
male.lee
carbohydrate (g/L)
300
200
100
0
0
40
80
120
protein (g/L)
dILP2
−0.5
−1.0
−1.5
female.tps
carbohydrate (g/L)
300
200
100
0
0
40
80
120
protein (g/L)
dILP2
−0.5
−1.0
−1.5

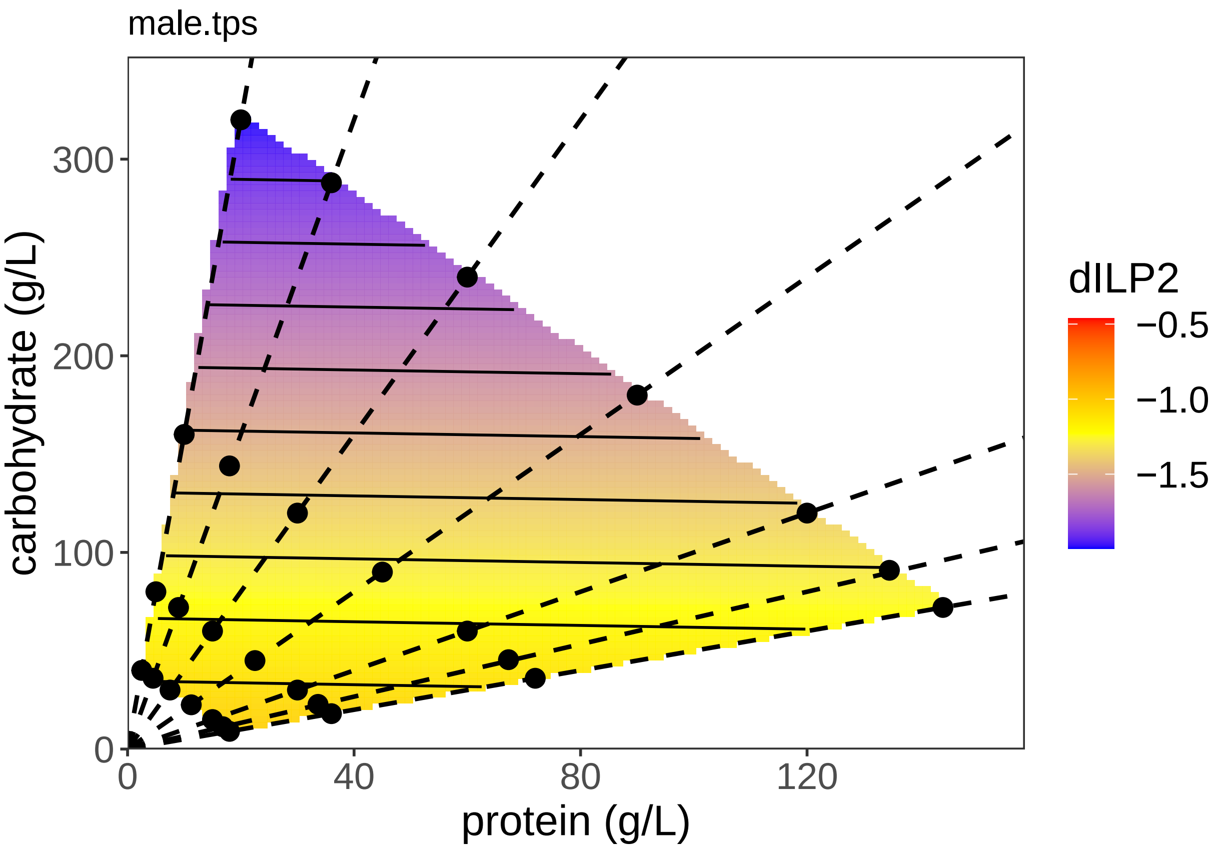


##### Analysis of dILP3

Test whether there is a difference between the sexes

```
dILP3Model1 <- lm(dILP3 ~ sex + (carb:prot + poly(carb, 2) +
    poly(prot, 2)), data = all.data)
dILP3Model2 <- lm(dILP3 ~ sex * (carb:prot + poly(carb, 2) +
    poly(prot, 2)), data = all.data)
anova(dILP3Model1, dILP3Model2)
```

```
## Analysis of Variance Table
##
## Model 1: dILP3 ~ sex + (carb:prot + poly(carb, 2) + poly(prot, 2))
## Model 2: dILP3 ~ sex * (carb:prot + poly(carb, 2) + poly(prot, 2))
##   Res.Df    RSS Df Sum of Sq      F Pr(>F)
## 1    147 131.70
## 2    142 130.48  5    1.2194 0.2654 0.9313
```

```
summary(dILP3Model1)
```

```
##
## Call:
## lm(formula = dILP3 ~ sex + (carb:prot + poly(carb, 2) + poly(prot,
##     2)), data = all.data)
##
## Residuals:
##     Min      1Q  Median      3Q     Max
```

```
## -4.2931 -0.4547  0.0255  0.5377  2.0818
##
## Coefficients:
##                   Estimate Std. Error t value Pr(>|t|)
## (Intercept)      8.210e-02  2.126e-01   0.386   0.7000
## sexM            -2.282e-01  1.528e-01  -1.494   0.1374
## poly(carb, 2)1 -2.376e+00  1.869e+00  -1.271   0.2056
## poly(carb, 2)2  4.744e-01  1.074e+00   0.442   0.6593
## poly(prot, 2)1 -4.819e+00  2.219e+00  -2.172   0.0314 *
## poly(prot, 2)2  1.064e+00  9.820e-01   1.084   0.2802
## carb:prot       5.808e-05  4.529e-05   1.282   0.2017
## ---
## Signif. codes:  0 '***' 0.001 '**' 0.01 '*' 0.05 '.' 0.1 ' ' 1
##
## Residual standard error: 0.9465 on 147 degrees of freedom
##   (5 observations deleted due to missingness)
## Multiple R-squared:  0.07479,    Adjusted R-squared:  0.03703
## F-statistic: 1.981 on 6 and 147 DF,  p-value: 0.07199
```

There is no sex*diet interaction, and expression level is the same in both sexes

Test the effect of C and P on dILP3 expression in males and females

```
dILP3.m <- lm(dILP3 ~ (carb:prot + poly(carb, 2) + poly(prot,
    2)), data = m.data)
dILP3.f <- lm(dILP3 ~ (carb:prot + poly(carb, 2) + poly(prot,
    2)), data = f.data)
summary(dILP3.m)
```

```
##
## Call:
## lm(formula = dILP3 ~ (carb:prot + poly(carb, 2) + poly(prot,
##     2)), data = m.data)
##
## Residuals:
##     Min      1Q  Median      3Q     Max
## -4.1442 -0.6863  0.0723  0.7399  2.2307
##
## Coefficients:
##                   Estimate Std. Error t value Pr(>|t|)
## (Intercept)     -2.457e-01  3.851e-01  -0.638    0.526
## poly(carb, 2)1 -1.705e+00  2.393e+00  -0.712    0.479
## poly(carb, 2)2 -2.052e-01  1.383e+00  -0.148    0.883
## poly(prot, 2)1 -4.305e+00  2.930e+00  -1.469    0.146
## poly(prot, 2)2  1.093e+00  1.288e+00   0.848    0.399
## carb:prot       7.999e-05  8.657e-05   0.924    0.359
##
## Residual standard error: 1.246 on 68 degrees of freedom
##   (1 observation deleted due to missingness)
## Multiple R-squared:  0.04876,    Adjusted R-squared:  -0.02118
## F-statistic: 0.6972 on 5 and 68 DF,  p-value: 0.6274
```

```
summary(dILP3.f)
```

```
##
## Call:
```

```
## lm(formula = dILP3 ~ (carb:prot + poly(carb, 2) + poly(prot,
##     2)), data = f.data)
##
## Residuals:
##      Min       1Q   Median       3Q      Max
## -1.57684 -0.34565  0.01863  0.38630  1.23255
##
## Coefficients:
##                  Estimate Std. Error t value Pr(>|t|)
## (Intercept)     1.751e-01  1.670e-01   1.048   0.2979
## poly(carb, 2)1 -1.591e+00  1.177e+00  -1.351   0.1807
## poly(carb, 2)2  8.782e-01  6.715e-01   1.308   0.1950
## poly(prot, 2)1 -2.547e+00  1.356e+00  -1.879   0.0642 .
## poly(prot, 2)2  4.414e-01  6.043e-01   0.730   0.4674
## carb:prot       3.746e-05  3.835e-05   0.977   0.3318
## ---
## Signif. codes:  0 '***' 0.001 '**' 0.01 '*' 0.05 '.' 0.1 ' ' 1
##
## Residual standard error: 0.5809 on 74 degrees of freedom
##   (4 observations deleted due to missingness)
## Multiple R-squared:  0.1467, Adjusted R-squared:  0.08909
## F-statistic: 2.545 on 5 and 74 DF,  p-value: 0.03512
```

Simplify the models:

```r
dILP3.m <- lm(dILP3 ~ prot, data = m.data)
dILP3.f <- lm(dILP3 ~ prot, data = f.data)
summary(dILP3.m)
```

```
##
## Call:
## lm(formula = dILP3 ~ prot, data = m.data)
##
## Residuals:
##     Min      1Q  Median      3Q     Max
## -4.4979 -0.6453  0.0609  0.8141  1.8769
##
## Coefficients:
##              Estimate Std. Error t value Pr(>|t|)
## (Intercept)  0.293628   0.210180   1.397    0.167
## prot        -0.004952   0.003563  -1.390    0.169
##
## Residual standard error: 1.225 on 72 degrees of freedom
##   (1 observation deleted due to missingness)
## Multiple R-squared:  0.02612,    Adjusted R-squared:  0.01259
## F-statistic: 1.931 on 1 and 72 DF,  p-value: 0.1689
```

```r
summary(dILP3.f)
```

```
##
## Call:
## lm(formula = dILP3 ~ prot, data = f.data)
##
## Residuals:
##      Min       1Q   Median       3Q      Max
## -1.42943 -0.40182 -0.00719  0.37409  1.41793
```

```
##
## Coefficients:
##              Estimate Std. Error t value Pr(>|t|)
## (Intercept)  0.519250   0.094713   5.482    5e-07 ***
## prot        -0.004857   0.001646  -2.950   0.0042 **
## ---
## Signif. codes:  0 '***' 0.001 '**' 0.01 '*' 0.05 '.' 0.1 ' ' 1
##
## Residual standard error: 0.581 on 78 degrees of freedom
##   (4 observations deleted due to missingness)
## Multiple R-squared:  0.1004, Adjusted R-squared:  0.08882
## F-statistic: 8.701 on 1 and 78 DF,  p-value: 0.004197
```

Re-test using the simplest model

```
dILP3Model0 <- lm(dILP3 ~ (prot), data = all.data)
dILP3Model1 <- lm(dILP3 ~ sex + (prot), data = all.data)
dILP3Model2 <- lm(dILP3 ~ sex * (prot), data = all.data)
anova(dILP3Model0, dILP3Model1, dILP3Model2)
```

```
## Analysis of Variance Table
##
## Model 1: dILP3 ~ (prot)
## Model 2: dILP3 ~ sex + (prot)
## Model 3: dILP3 ~ sex * (prot)
##   Res.Df    RSS Df Sum of Sq      F Pr(>F)
## 1    152 136.38
## 2    151 134.35  1   2.02707 2.2632 0.1346
## 3    150 134.35  1   0.00055 0.0006 0.9803
```

```
summary(dILP3Model2)
```

```
##
## Call:
## lm(formula = dILP3 ~ sex * (prot), data = all.data)
##
## Residuals:
##     Min      1Q  Median      3Q     Max
## -4.4979 -0.4718 -0.0001  0.5712  1.8769
##
## Coefficients:
##               Estimate Std. Error t value Pr(>|t|)
## (Intercept)  5.192e-01  1.543e-01   3.366  0.00097 ***
## sexM        -2.256e-01  2.240e-01  -1.007  0.31543
## prot        -4.857e-03  2.682e-03  -1.811  0.07216 .
## sexM:prot   -9.503e-05  3.843e-03  -0.025  0.98031
## ---
## Signif. codes:  0 '***' 0.001 '**' 0.01 '*' 0.05 '.' 0.1 ' ' 1
##
## Residual standard error: 0.9464 on 150 degrees of freedom
##   (5 observations deleted due to missingness)
## Multiple R-squared:  0.05618,    Adjusted R-squared:  0.0373
## F-statistic: 2.976 on 3 and 150 DF,  p-value: 0.03354
```

Still no difference between males and females There is a marginal effect of protein in females but not in females

We can do a power analysis on detecting a sex:protein interaction

```
RsqFull <- summary(dILP3Model2)$r.squared
RsqRed <- summary(dILP3Model1)$r.squared


fsq <- (RsqFull - RsqRed)/(1 - RsqFull)

predFull <- length(dILP3Model2$coefficients)
predRed <- length(dILP3Model1$coefficients)


wp.regression(n = nobs(dILP2Model2), p1 = predFull, p2 = predRed,
    f2 = fsq)
```

```
## Power for multiple regression
##
##       n p1 p2           f2 alpha      power
##     127  4  3 4.075692e-06  0.05 0.05005699
##
## URL: http://psychstat.org/regression
```

We can do a power analysis on the linear effect of protein on expression in males

```
anova(dILP3.m)
```

```
## Analysis of Variance Table
##
## Response: dILP3
##           Df  Sum Sq Mean Sq F value Pr(>F)
## prot       1   2.897  2.8972  1.9311 0.1689
## Residuals 72 108.018  1.5002
```

```
summary(dILP3.m)
```

```
##
## Call:
## lm(formula = dILP3 ~ prot, data = m.data)
##
## Residuals:
##     Min      1Q  Median      3Q     Max
## -4.4979 -0.6453  0.0609  0.8141  1.8769
##
## Coefficients:
##              Estimate Std. Error t value Pr(>|t|)
## (Intercept)  0.293628   0.210180   1.397    0.167
## prot        -0.004952   0.003563  -1.390    0.169
##
## Residual standard error: 1.225 on 72 degrees of freedom
##   (1 observation deleted due to missingness)
## Multiple R-squared:  0.02612,    Adjusted R-squared:  0.01259
## F-statistic: 1.931 on 1 and 72 DF,  p-value: 0.1689
```

```
pwr.f2.test(u = 1, v = summary(dILP3.m)$df[2], f2 = (summary(dILP3.m)$r.squared/(1 -
    summary(dILP3.m)$r.squared)), sig.level = 0.05)
```

```
##
```

```
##      Multiple regression power calculation
##
##               u = 1
##               v = 72
##              f2 = 0.02682121
##       sig.level = 0.05
##           power = 0.2847729
```

Plot the model and the TPS

```
## Warning:
## Grid searches over lambda (nugget and sill variances) with  minima at the endpoints:
##   (GCV) Generalized Cross-Validation
##    minimum at  right endpoint  lambda  =  268.0589 (eff. df= 3.00101 )
```

```
## Warning:
## Grid searches over lambda (nugget and sill variances) with  minima at the endpoints:
##   (GCV) Generalized Cross-Validation
##    minimum at  right endpoint  lambda  =  246.4776 (eff. df= 3.001005 )
```

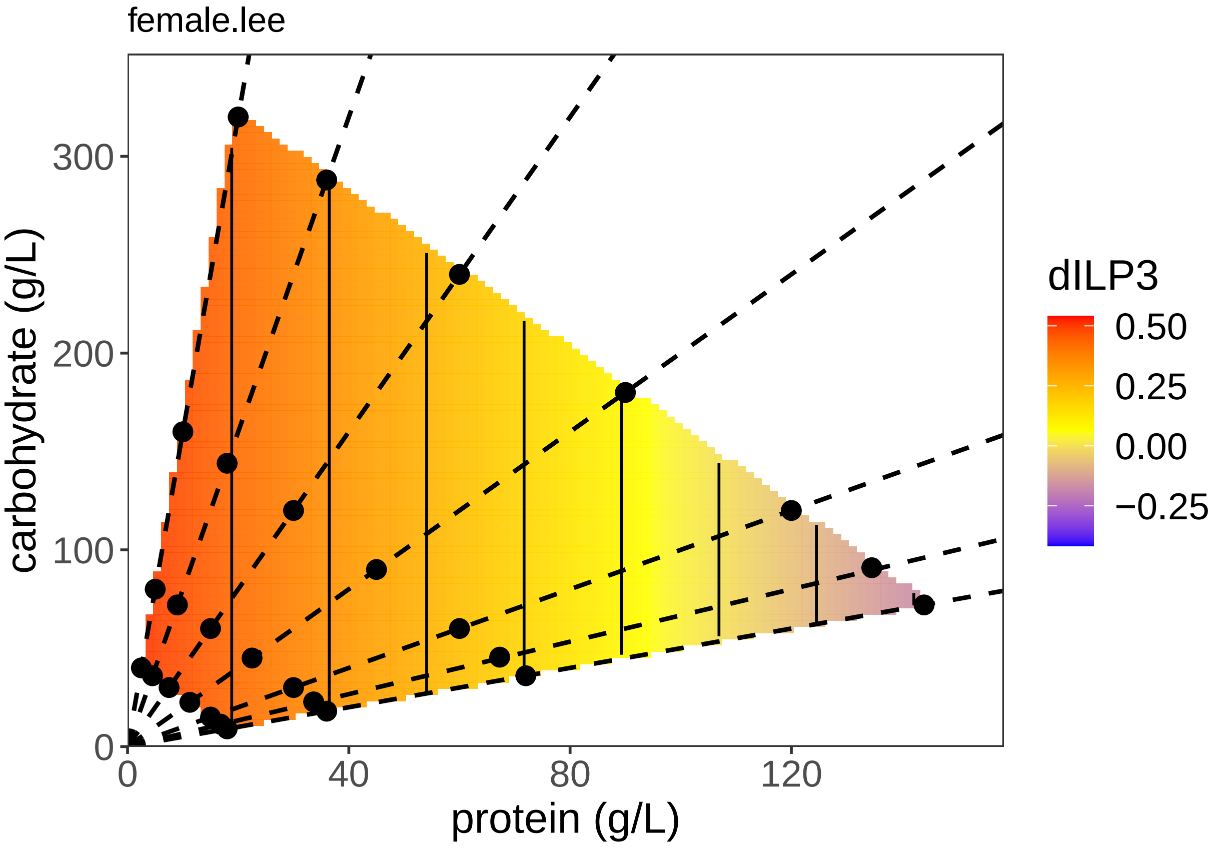

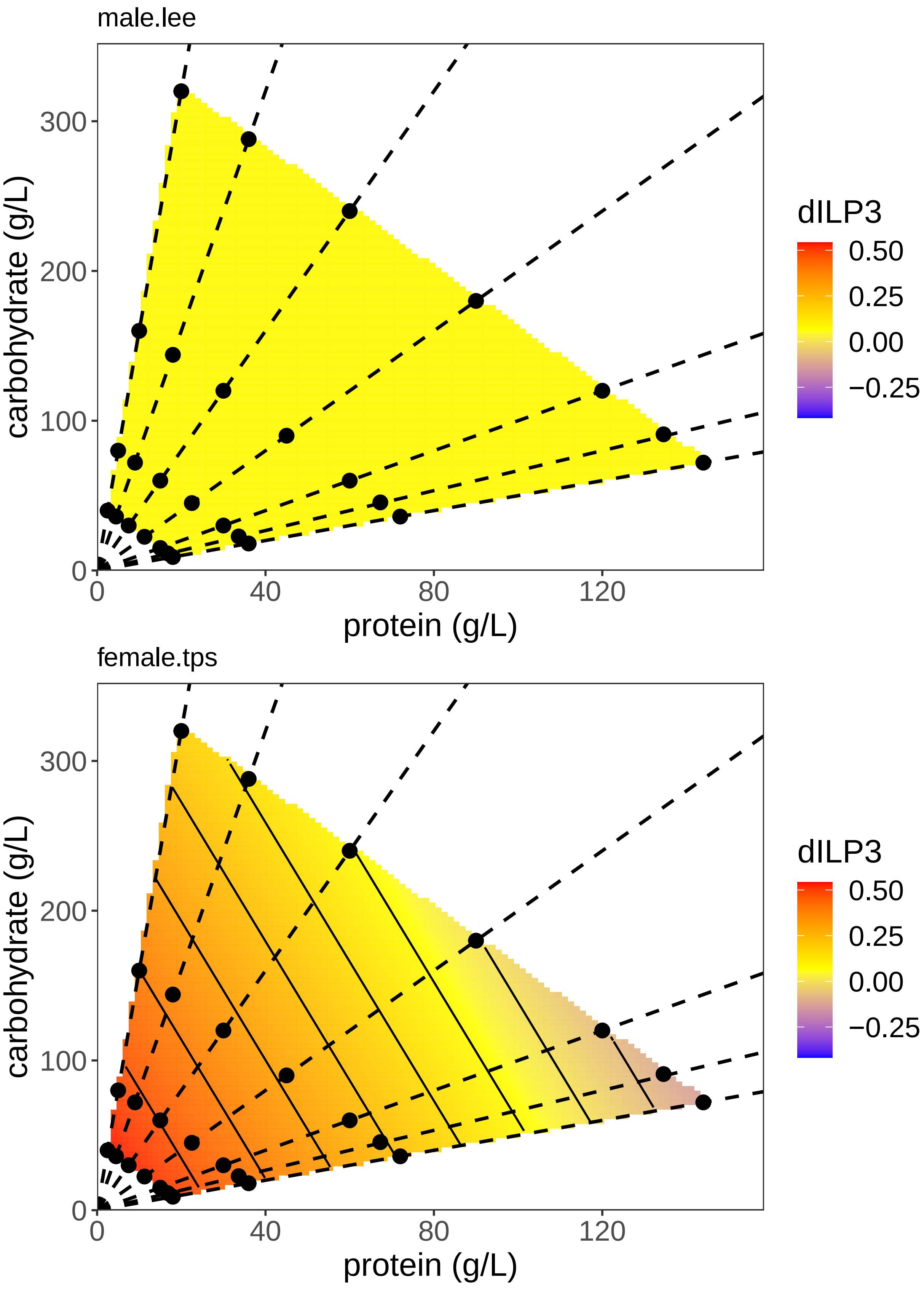
male.lee
carbohydrate (g/L)
300
200
100
0
0
40
80
120
protein (g/L)
dILP3
0.50
0.25
0.00
−0.25
female.tps
carbohydrate (g/L)
300
200
100
0
0
40
80
120
protein (g/L)
dILP3
0.50
0.25
0.00
−0.25

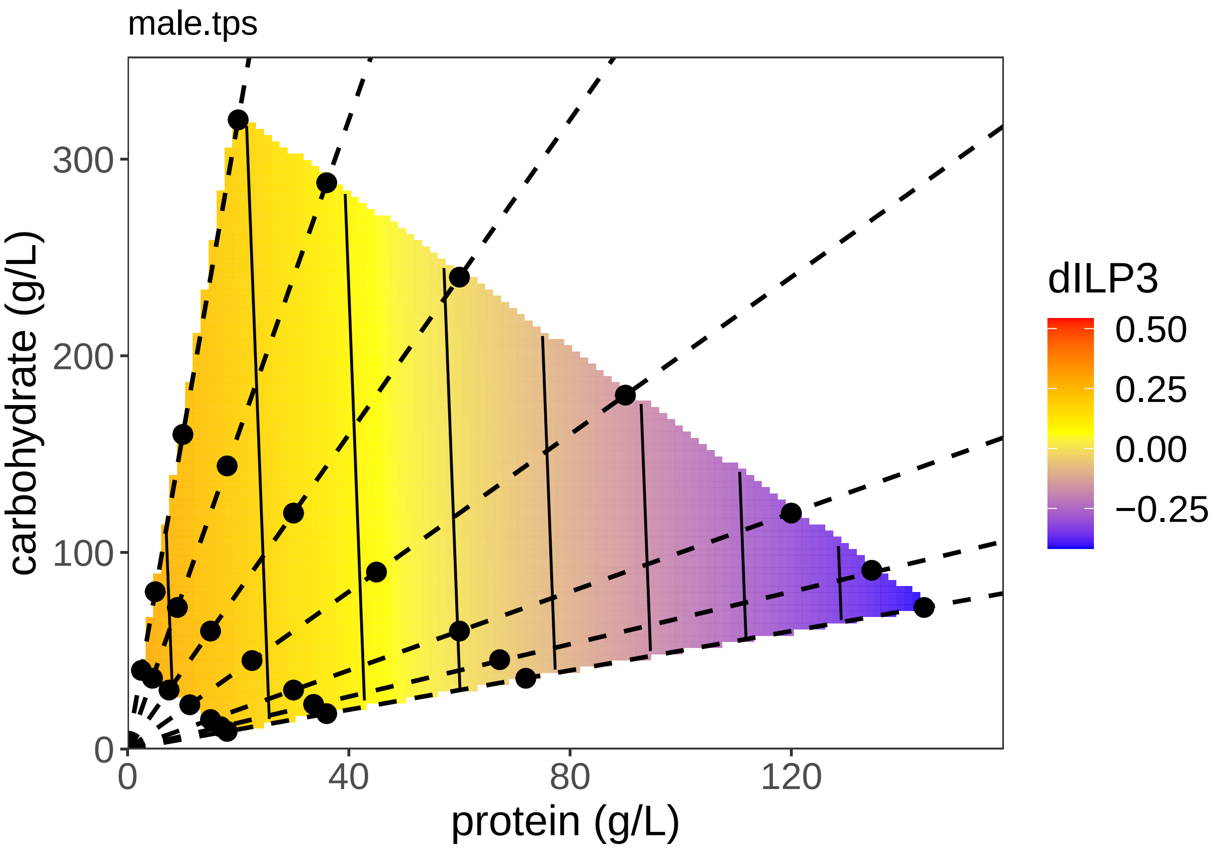


##### Analysis of dILP5

Test whether there is a difference between the sexes

```
dILP5Model1 <- lm(dILP5 ~ sex + (carb:prot + poly(carb, 2) +
    poly(prot, 2)), data = all.data)
dILP5Model2 <- lm(dILP5 ~ sex * (carb:prot + poly(carb, 2) +
    poly(prot, 2)), data = all.data)
anova(dILP5Model1, dILP5Model2)
```

```
## Analysis of Variance Table
##
## Model 1: dILP5 ~ sex + (carb:prot + poly(carb, 2) + poly(prot, 2))
## Model 2: dILP5 ~ sex * (carb:prot + poly(carb, 2) + poly(prot, 2))
##   Res.Df   RSS Df Sum of Sq      F Pr(>F)
## 1    145 138.0
## 2    140 134.9  5       3.1 0.6435 0.6669
```

There is no sex*protein interaction

Test the effect of C and P on dILP5 expression in males and females

```
dILP5.m <- lm(dILP5 ~ (carb:prot + poly(carb, 2) + poly(prot,
    2)), data = m.data)
dILP5.f <- lm(dILP5 ~ (carb:prot + poly(carb, 2) + poly(prot,
    2)), data = f.data)
summary(dILP5.m)
```

```
##
## Call:
## lm(formula = dILP5 ~ (carb:prot + poly(carb, 2) + poly(prot,
##     2)), data = m.data)
##
## Residuals:
##      Min       1Q   Median       3Q      Max
## -2.97679 -0.83305  0.00411  0.73832  2.00276
##
## Coefficients:
##                  Estimate Std. Error t value Pr(>|t|)
## (Intercept)     9.155e-01  3.277e-01   2.794  0.00681 **
## poly(carb, 2)1 -2.827e+00  2.054e+00  -1.377  0.17331
## poly(carb, 2)2  1.138e+00  1.193e+00   0.954  0.34352
## poly(prot, 2)1 -2.391e+00  2.510e+00  -0.952  0.34433
## poly(prot, 2)2  1.205e-01  1.108e+00   0.109  0.91377
## carb:prot       4.596e-05  7.355e-05   0.625  0.53419
## ---
## Signif. codes:  0 '***' 0.001 '**' 0.01 '*' 0.05 '.' 0.1 ' ' 1
##
## Residual standard error: 1.058 on 66 degrees of freedom
##   (3 observations deleted due to missingness)
## Multiple R-squared:  0.08248,    Adjusted R-squared:  0.01297
## F-statistic: 1.187 on 5 and 66 DF,  p-value: 0.3253
```

```
summary(dILP5.f)
```

```
##
## Call:
## lm(formula = dILP5 ~ (carb:prot + poly(carb, 2) + poly(prot,
##     2)), data = f.data)
##
## Residuals:
##      Min       1Q   Median       3Q      Max
## -2.05849 -0.53357 -0.02815  0.64357  1.65136
##
## Coefficients:
##                  Estimate Std. Error t value Pr(>|t|)
## (Intercept)     4.904e-01  2.610e-01   1.879   0.0642 .
## poly(carb, 2)1 -3.175e+00  1.840e+00  -1.725   0.0887 .
## poly(carb, 2)2  2.018e+00  1.050e+00   1.922   0.0584 .
## poly(prot, 2)1 -1.272e+00  2.120e+00  -0.600   0.5504
## poly(prot, 2)2 -1.695e+00  9.446e-01  -1.795   0.0768 .
## carb:prot       6.556e-05  5.994e-05   1.094   0.2776
## ---
## Signif. codes:  0 '***' 0.001 '**' 0.01 '*' 0.05 '.' 0.1 ' ' 1
##
## Residual standard error: 0.908 on 74 degrees of freedom
##   (4 observations deleted due to missingness)
## Multiple R-squared:  0.123,  Adjusted R-squared:  0.06375
## F-statistic: 2.076 on 5 and 74 DF,  p-value: 0.07794
```

There is an effect of protein in females but not in males, but only when you include higher order interactions.

Simplify:

```r
# Non-orthoganol Coef
summary(lm(dILP5 ~ (poly(carb, 1) + poly(prot, 2, raw = TRUE)),
    data = f.data))[4]
```

```
## $coefficients
##                                Estimate   Std. Error   t value   Pr(>|t|)
## (Intercept)                0.4061695267 2.226727e-01  1.824065 0.07207311
## poly(carb, 1)             -1.5896548921 1.010509e+00 -1.573123 0.11984446
## poly(prot, 2, raw = TRUE)1  0.0194485870 9.624513e-03  2.020735 0.04683010
## poly(prot, 2, raw = TRUE)2 -0.0001415936 6.783467e-05 -2.087334 0.04020999
```

```r
# Orthoganol Test
dILP5.m <- lm(dILP5 ~ (poly(carb, 1) + poly(prot, 2)), data = m.data)
dILP5.f <- lm(dILP5 ~ (poly(carb, 1) + poly(prot, 2)), data = f.data)
summary(dILP5.m)
```

```
##
## Call:
## lm(formula = dILP5 ~ (poly(carb, 1) + poly(prot, 2)), data = m.data)
##
## Residuals:
##      Min       1Q   Median       3Q      Max
## -3.07906 -0.80693 -0.00032  0.74391  2.14899
##
## Coefficients:
##                 Estimate Std. Error t value Pr(>|t|)
## (Intercept)      1.10418    0.12404   8.902 5.08e-13 ***
## poly(carb, 1)   -1.76877    1.13154  -1.563    0.123
## poly(prot, 2)1 -1.39927    1.11269  -1.258    0.213
## poly(prot, 2)2 -0.01021    1.09434  -0.009    0.993
## ---
## Signif. codes:  0 '***' 0.001 '**' 0.01 '*' 0.05 '.' 0.1 ' ' 1
##
## Residual standard error: 1.051 on 68 degrees of freedom
##   (3 observations deleted due to missingness)
## Multiple R-squared:  0.06676,    Adjusted R-squared:  0.02559
## F-statistic: 1.621 on 3 and 68 DF,  p-value: 0.1925
```

```r
summary(dILP5.f)
```

```
##
## Call:
## lm(formula = dILP5 ~ (poly(carb, 1) + poly(prot, 2)), data = f.data)
##
## Residuals:
##      Min       1Q   Median       3Q      Max
## -2.24726 -0.58188 -0.01893  0.61509  1.89785
##
## Coefficients:
##                 Estimate Std. Error t value Pr(>|t|)
## (Intercept)       0.7460     0.1032   7.225  3.3e-10 ***
## poly(carb, 1)    -1.5897     1.0105  -1.573   0.1198
## poly(prot, 2)1    0.1477     0.9469   0.156   0.8765
## poly(prot, 2)2   -1.9817     0.9494  -2.087   0.0402 *
## ---
```

```
## Signif. codes:  0 '***' 0.001 '**' 0.01 '*' 0.05 '.' 0.1 ' ' 1
##
## Residual standard error: 0.9224 on 76 degrees of freedom
##   (4 observations deleted due to missingness)
## Multiple R-squared:  0.07066,    Adjusted R-squared:  0.03397
## F-statistic: 1.926 on 3 and 76 DF,  p-value: 0.1325
```

There is an effect of protein in females but not in males, but only when you include carbohydrate.

Re-test using the simplest model

```
dILP5Model0 <- lm(dILP5 ~ (poly(carb, 1) + poly(prot, 2)), data = all.data)
dILP5Model1 <- lm(dILP5 ~ sex + (poly(carb, 1) + poly(prot, 2)),
    data = all.data)
dILP5Model2 <- lm(dILP5 ~ sex * (poly(carb, 1) + poly(prot, 2)),
    data = all.data)
anova(dILP5Model0, dILP5Model1, dILP5Model2)
```

```
## Analysis of Variance Table
##
## Model 1: dILP5 ~ (poly(carb, 1) + poly(prot, 2))
## Model 2: dILP5 ~ sex + (poly(carb, 1) + poly(prot, 2))
## Model 3: dILP5 ~ sex * (poly(carb, 1) + poly(prot, 2))
##   Res.Df    RSS Df Sum of Sq      F  Pr(>F)
## 1    148 147.40
## 2    147 142.85  1    4.5507 4.6873 0.03203 *
## 3    144 139.80  3    3.0460 1.0458 0.37431
## ---
## Signif. codes:  0 '***' 0.001 '**' 0.01 '*' 0.05 '.' 0.1 ' ' 1
```

```
summary(dILP5Model1)
```

```
##
## Call:
## lm(formula = dILP5 ~ sex + (poly(carb, 1) + poly(prot, 2)), data = all.data)
##
## Residuals:
##     Min      1Q  Median      3Q     Max
## -3.2765 -0.6640  0.0488  0.7149  2.2774
##
## Coefficients:
##                Estimate Std. Error t value Pr(>|t|)
## (Intercept)      0.7537     0.1102   6.838    2e-10 ***
## sexM             0.3468     0.1602   2.164   0.0321 *
## poly(carb, 1)   -2.3492     1.0712  -2.193   0.0299 *
## poly(prot, 2)1  -0.8738     1.0269  -0.851   0.3962
## poly(prot, 2)2  -1.4921     1.0207  -1.462   0.1459
## ---
## Signif. codes:  0 '***' 0.001 '**' 0.01 '*' 0.05 '.' 0.1 ' ' 1
##
## Residual standard error: 0.9858 on 147 degrees of freedom
##   (7 observations deleted due to missingness)
## Multiple R-squared:  0.07844,    Adjusted R-squared:  0.05336
## F-statistic: 3.128 on 4 and 147 DF,  p-value: 0.0167
```

Still no significant interaction, although males have higher expression

We can do a power analysis on the sex interaction:

```r
RsqFull <- summary(dILP5Model2)$r.squared
RsqRed <- summary(dILP5Model1)$r.squared


fsq <- (RsqFull - RsqRed)/(1 - RsqFull)

predFull <- length(dILP5Model2$coefficients)
predRed <- length(dILP5Model1$coefficients)


wp.regression(n = nobs(dILP2Model2), p1 = predFull, p2 = predRed,
    f2 = fsq)
```

```
## Power for multiple regression
##
##       n p1 p2         f2 alpha     power
##     127  8  5 0.02178796  0.05 0.2389557
##
## URL: http://psychstat.org/regression
```

Plot the model and the TPS

```
## Warning:
## Grid searches over lambda (nugget and sill variances) with  minima at the endpoints:
##   (GCV) Generalized Cross-Validation
##    minimum at  right endpoint  lambda  =  268.0589 (eff. df= 3.00101 )
```

```
## Warning:
## Grid searches over lambda (nugget and sill variances) with  minima at the endpoints:
##   (GCV) Generalized Cross-Validation
##    minimum at  right endpoint  lambda  =  242.5057 (eff. df= 3.000997 )
```

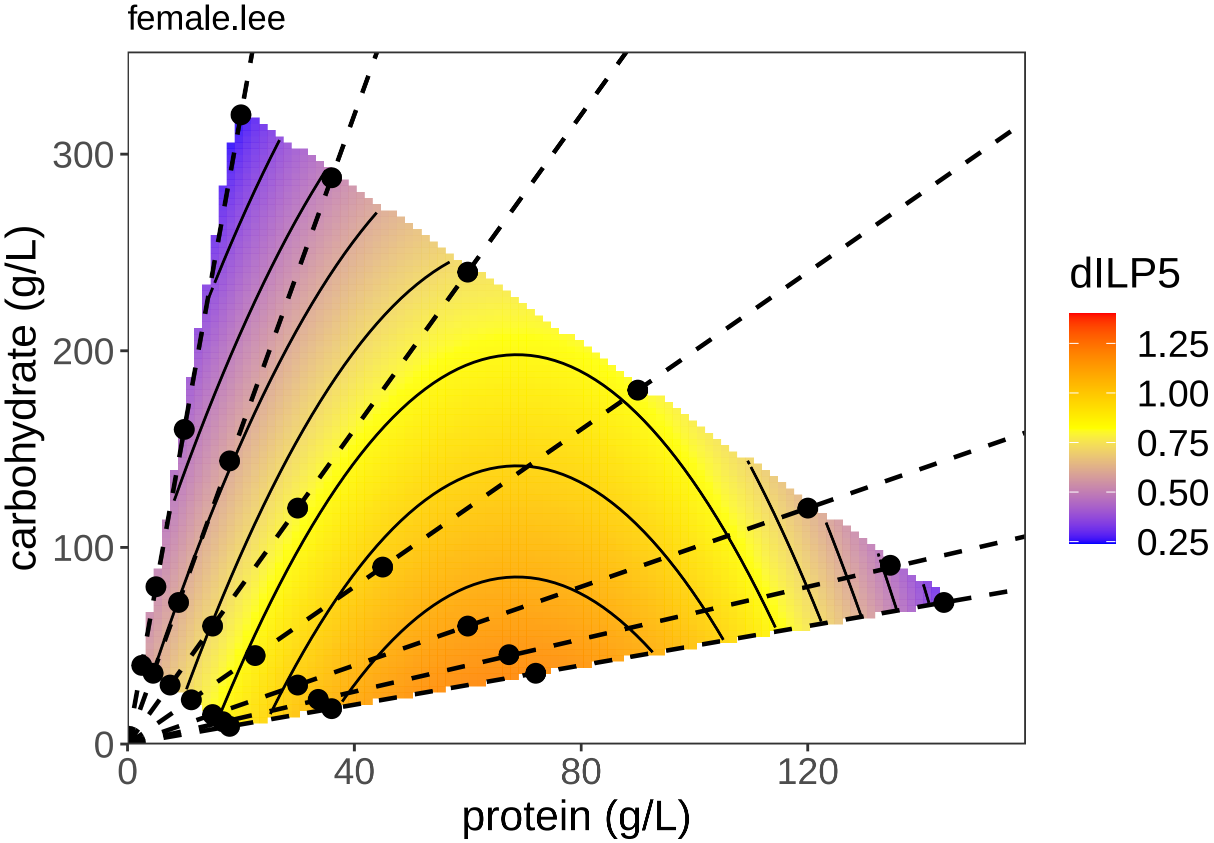
female.lee
carbohydrate (g/L)
300
200
100
0
0
40
80
120
protein (g/L)
dILP5
1.25
1.00
0.75
0.50
0.25

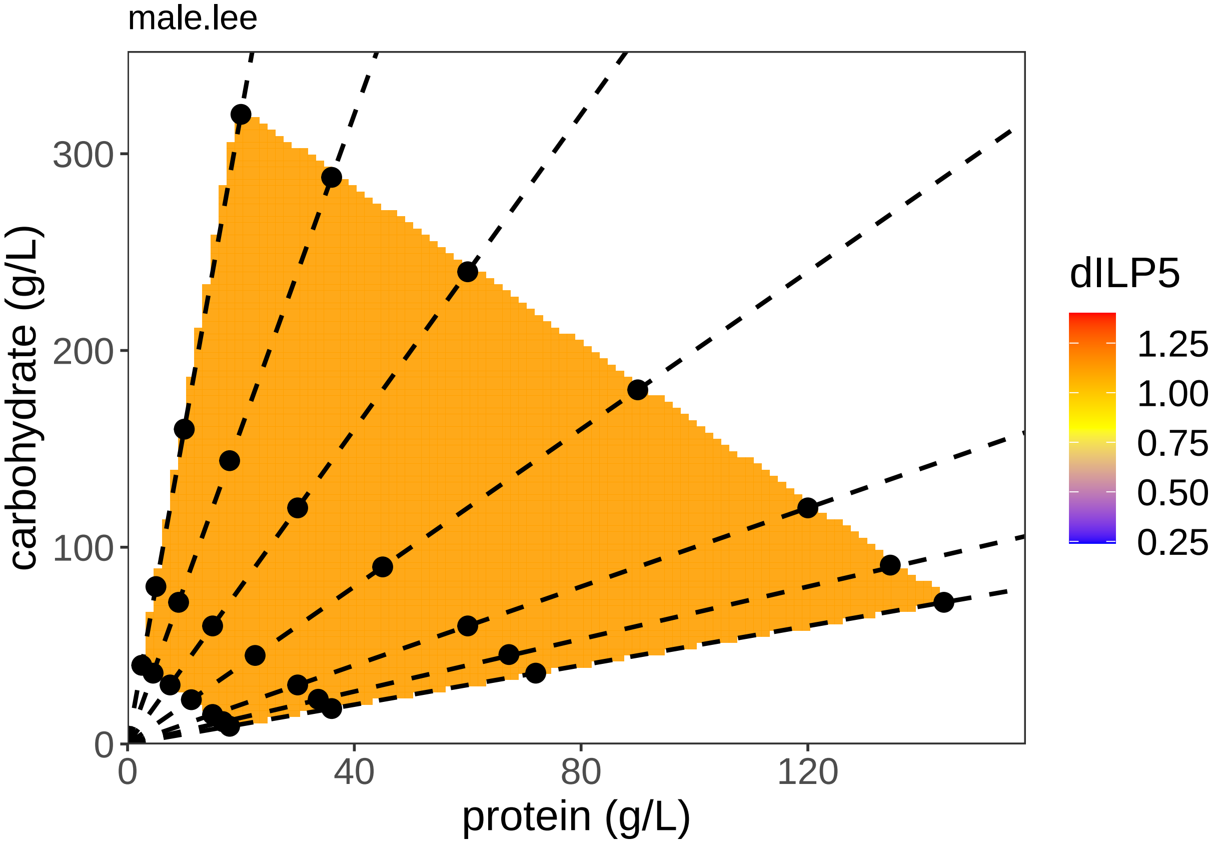
male.lee
carbohydrate (g/L)
300
200
100
0
0
40
80
120
protein (g/L)
dILP5
1.25
1.00
0.75
0.50
0.25

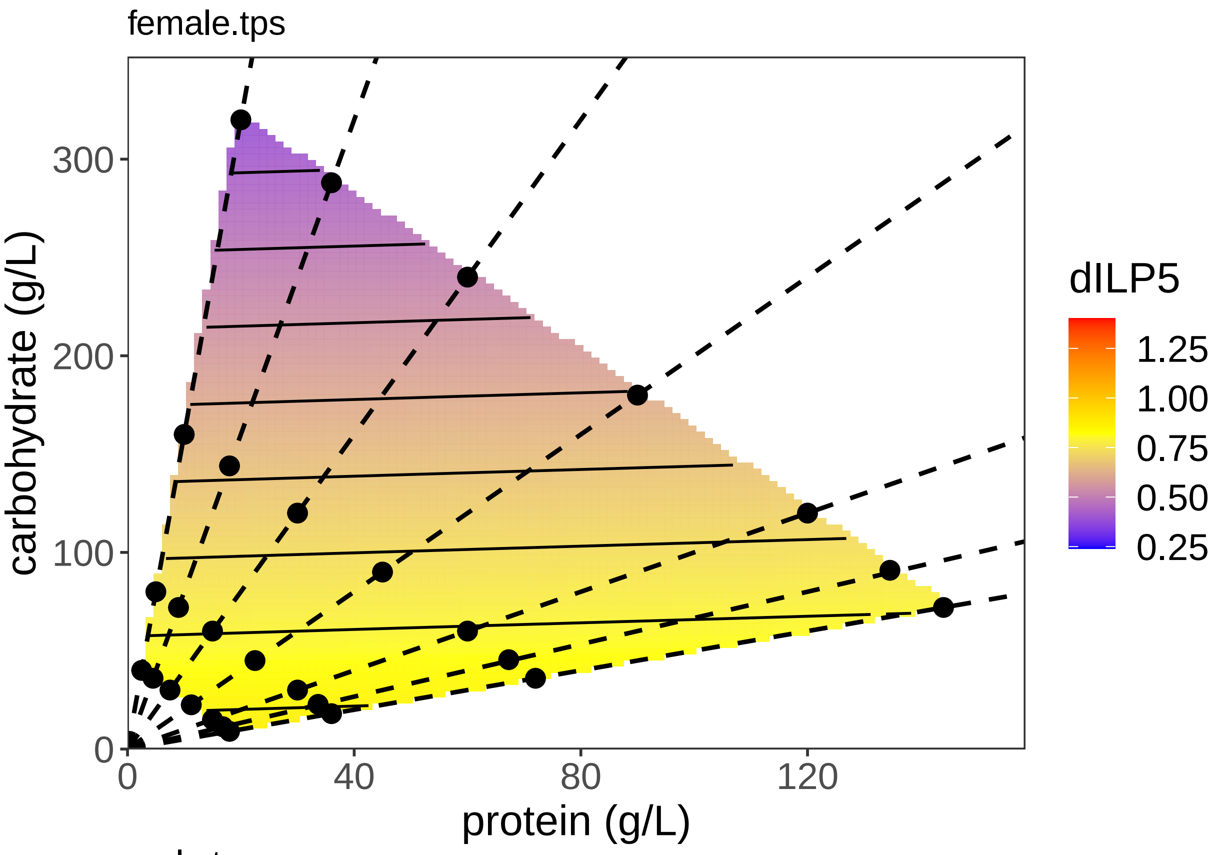
female.tps
carbohydrate (g/L)
protein (g/L)
0
100
200
300
0
40
80
120
dILP5
1.25
1.00
0.75
0.50
0.25

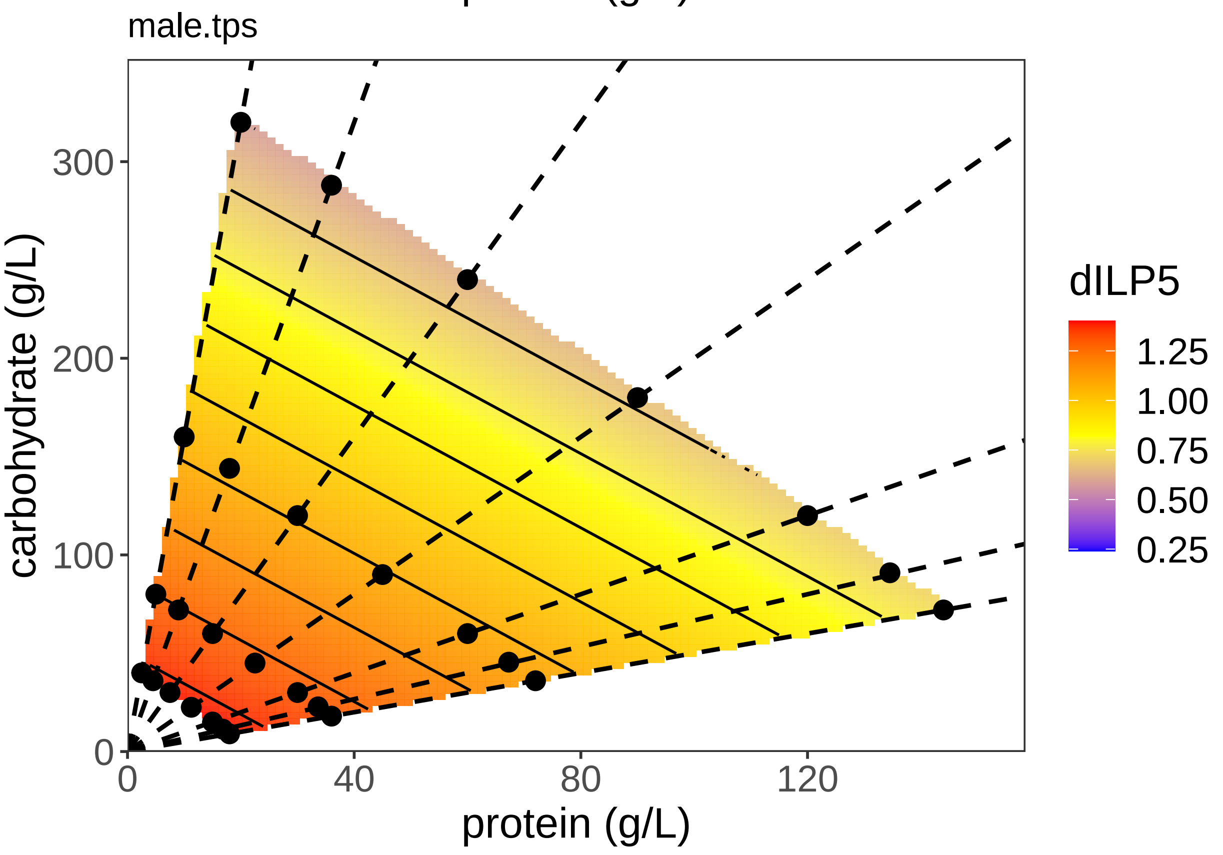
male.tps
carbohydrate (g/L)
protein (g/L)
0
100
200
300
0
40
80
120
dILP5
1.25
1.00
0.75
0.50
0.25

#### Analysis of dILP8

Test whether there is a difference between the sexes

```r
dILP8Model1 <- lm(dILP8 ~ sex + (carb:prot + poly(carb, 2) +
    poly(prot, 2)), data = all.data)
dILP8Model2 <- lm(dILP8 ~ sex * (carb:prot + poly(carb, 2) +
    poly(prot, 2)), data = all.data)
anova(dILP8Model1, dILP8Model2)
```

```
## Analysis of Variance Table
##
## Model 1: dILP8 ~ sex + (carb:prot + poly(carb, 2) + poly(prot, 2))
## Model 2: dILP8 ~ sex * (carb:prot + poly(carb, 2) + poly(prot, 2))
##   Res.Df    RSS Df Sum of Sq      F  Pr(>F)
## 1    118 203.28
## 2    113 185.70  5    17.581 2.1397 0.06576 .
## ---
## Signif. codes:  0 '***' 0.001 '**' 0.01 '*' 0.05 '.' 0.1 ' ' 1
```

```r
summary(dILP8Model1)
```

```
##
## Call:
## lm(formula = dILP8 ~ sex + (carb:prot + poly(carb, 2) + poly(prot,
##     2)), data = all.data)
##
## Residuals:
##     Min      1Q  Median      3Q     Max
## -3.1116 -0.8288 -0.1546  0.6183  3.8612
##
## Coefficients:
##                  Estimate Std. Error t value Pr(>|t|)
## (Intercept)    -3.416e+00  3.237e-01 -10.553  < 2e-16 ***
## sexM            1.367e+00  2.424e-01   5.642 1.17e-07 ***
## poly(carb, 2)1  1.050e+00  2.780e+00   0.378   0.7064
## poly(carb, 2)2 -1.282e+00  1.597e+00  -0.802   0.4239
## poly(prot, 2)1 -3.757e+00  3.486e+00  -1.078   0.2834
## poly(prot, 2)2  2.850e+00  1.561e+00   1.825   0.0705 .
## carb:prot      -5.801e-05  7.008e-05  -0.828   0.4095
## ---
## Signif. codes:  0 '***' 0.001 '**' 0.01 '*' 0.05 '.' 0.1 ' ' 1
##
## Residual standard error: 1.313 on 118 degrees of freedom
##   (34 observations deleted due to missingness)
## Multiple R-squared:  0.3376, Adjusted R-squared:  0.3039
## F-statistic: 10.02 on 6 and 118 DF,  p-value: 6.213e-09
```

There is no sex*diet interaction but male expression is higher.

Test the effect of C and P on dILP8 expression in males and females

```r
dILP8.m <- lm(dILP8 ~ (carb:prot + poly(carb, 2) + poly(prot,
    2)), data = m.data)
dILP8.f <- lm(dILP8 ~ (carb:prot + poly(carb, 2) + poly(prot,
    2)), data = f.data)
summary(dILP8.m)
```

```
##
## Call:
## lm(formula = dILP8 ~ (carb:prot + poly(carb, 2) + poly(prot,
##     2)), data = m.data)
##
## Residuals:
##     Min      1Q  Median      3Q     Max
## -2.5028 -0.5241 -0.1313  0.5150  3.8538
##
## Coefficients:
##                  Estimate Std. Error t value Pr(>|t|)
## (Intercept)    -1.5918552  0.5336418  -2.983  0.00469 **
## poly(carb, 2)1  2.6867995  3.0158676   0.891  0.37795
## poly(carb, 2)2 -1.7559316  1.6636539  -1.055  0.29711
## poly(prot, 2)1  3.9402870  4.5819990   0.860  0.39459
## poly(prot, 2)2  0.9093467  2.0203242   0.450  0.65490
## carb:prot      -0.0001507  0.0001244  -1.212  0.23230
## ---
## Signif. codes:  0 '***' 0.001 '**' 0.01 '*' 0.05 '.' 0.1 ' ' 1
##
## Residual standard error: 1.31 on 43 degrees of freedom
##   (26 observations deleted due to missingness)
## Multiple R-squared:  0.06269,    Adjusted R-squared:  -0.0463
## F-statistic: 0.5752 on 5 and 43 DF,  p-value: 0.7186
```

```r
summary(dILP8.f)
```

```
##
## Call:
## lm(formula = dILP8 ~ (carb:prot + poly(carb, 2) + poly(prot,
##     2)), data = f.data)
##
## Residuals:
##     Min      1Q  Median      3Q     Max
## -2.4393 -0.7959 -0.1933  0.4806  3.9637
##
## Coefficients:
##                  Estimate Std. Error t value Pr(>|t|)
## (Intercept)    -3.683e+00  3.680e-01 -10.010 3.84e-15 ***
## poly(carb, 2)1 -7.704e-01  2.586e+00  -0.298  0.76663
## poly(carb, 2)2 -1.464e-01  1.490e+00  -0.098  0.92202
## poly(prot, 2)1 -6.897e+00  2.976e+00  -2.317  0.02342 *
## poly(prot, 2)2  3.629e+00  1.345e+00   2.699  0.00872 **
## carb:prot       1.459e-05  8.410e-05   0.173  0.86279
## ---
## Signif. codes:  0 '***' 0.001 '**' 0.01 '*' 0.05 '.' 0.1 ' ' 1
##
## Residual standard error: 1.264 on 70 degrees of freedom
##   (8 observations deleted due to missingness)
## Multiple R-squared:  0.3177, Adjusted R-squared:  0.269
## F-statistic: 6.519 on 5 and 70 DF,  p-value: 4.963e-05
```

Protein effects dILP8 in females but not in males.

Test the simpler model on dILP8 expression in males and females

```r
# Non-orthoganol Coef
summary(lm(dILP8 ~ (poly(prot, 2, raw = TRUE)), data = f.data))[4]
```

```
## $coefficients
##                                Estimate   Std. Error   t value     Pr(>|t|)
## (Intercept)               -2.2908053812 2.940779e-01 -7.789791 3.494185e-11
## poly(prot, 2, raw = TRUE)1 -0.0536213974 1.279359e-02 -4.191272 7.680640e-05
## poly(prot, 2, raw = TRUE)2  0.0002649604 9.081879e-05  2.917462 4.687016e-03
```

```r
# Orthoganol Test
dILP8.m <- lm(dILP8 ~ (poly(prot, 2)), data = m.data)
dILP8.f <- lm(dILP8 ~ (poly(prot, 2)), data = f.data)
summary(dILP8.m)
```

```
##
## Call:
## lm(formula = dILP8 ~ (poly(prot, 2)), data = m.data)
##
## Residuals:
##     Min      1Q  Median      3Q     Max
## -2.5949 -0.8071 -0.0448  0.4390  3.7340
##
## Coefficients:
##                 Estimate Std. Error t value Pr(>|t|)
## (Intercept)      -2.1932     0.1936 -11.328 6.66e-15 ***
## poly(prot, 2)1   -0.2329     2.1245  -0.110    0.913
## poly(prot, 2)2    0.7797     1.9085   0.409    0.685
## ---
## Signif. codes:  0 '***' 0.001 '**' 0.01 '*' 0.05 '.' 0.1 ' ' 1
##
## Residual standard error: 1.305 on 46 degrees of freedom
##   (26 observations deleted due to missingness)
## Multiple R-squared:  0.00552,    Adjusted R-squared:  -0.03772
## F-statistic: 0.1277 on 2 and 46 DF,  p-value: 0.8805
```

```r
summary(dILP8.f)
```

```
##
## Call:
## lm(formula = dILP8 ~ (poly(prot, 2)), data = f.data)
##
## Residuals:
##     Min      1Q  Median      3Q     Max
## -2.4128 -0.7713 -0.2479  0.4914  3.9844
##
## Coefficients:
##                 Estimate Std. Error t value Pr(>|t|)
## (Intercept)      -3.6241     0.1423 -25.471  < 2e-16 ***
## poly(prot, 2)1   -6.4359     1.2618  -5.101 2.58e-06 ***
## poly(prot, 2)2    3.7084     1.2711   2.917  0.00469 **
## ---
## Signif. codes:  0 '***' 0.001 '**' 0.01 '*' 0.05 '.' 0.1 ' ' 1
##
## Residual standard error: 1.239 on 73 degrees of freedom
##   (8 observations deleted due to missingness)
```

```
## Multiple R-squared:  0.3165, Adjusted R-squared:  0.2978
## F-statistic:  16.9 on 2 and 73 DF,  p-value: 9.281e-07
```

Still no effect in males

Test using the simpler model

```r
dILP8Model1 <- lm(dILP8 ~ sex + (poly(prot, 2)), data = all.data)
dILP8Model2 <- lm(dILP8 ~ sex * (poly(prot, 2)), data = all.data)
anova(dILP8Model1, dILP8Model2)
```

```
## Analysis of Variance Table
##
## Model 1: dILP8 ~ sex + (poly(prot, 2))
## Model 2: dILP8 ~ sex * (poly(prot, 2))
##   Res.Df    RSS Df Sum of Sq      F   Pr(>F)
## 1    121 205.98
## 2    119 190.40  2    15.582 4.8693 0.009276 **
## ---
## Signif. codes:  0 '***' 0.001 '**' 0.01 '*' 0.05 '.' 0.1 ' ' 1
```

```r
summary(dILP8Model2)
```

```
##
## Call:
## lm(formula = dILP8 ~ sex * (poly(prot, 2)), data = all.data)
##
## Residuals:
##     Min      1Q  Median      3Q     Max
## -2.5949 -0.7920 -0.1970  0.4733  3.9844
##
## Coefficients:
##                      Estimate Std. Error t value Pr(>|t|)
## (Intercept)           -3.6506     0.1451 -25.151  < 2e-16 ***
## sexM                   1.4600     0.2361   6.185 9.05e-09 ***
## poly(prot, 2)1        -8.7656     1.7924  -4.890 3.18e-06 ***
## poly(prot, 2)2         5.0963     1.7831   2.858  0.00503 **
## sexM:poly(prot, 2)1    8.3832     3.4270   2.446  0.01590 *
## sexM:poly(prot, 2)2   -3.9572     3.2381  -1.222  0.22409
## ---
## Signif. codes:  0 '***' 0.001 '**' 0.01 '*' 0.05 '.' 0.1 ' ' 1
##
## Residual standard error: 1.265 on 119 degrees of freedom
##   (34 observations deleted due to missingness)
## Multiple R-squared:  0.3796, Adjusted R-squared:  0.3535
## F-statistic: 14.56 on 5 and 119 DF,  p-value: 4e-11
```

There is a sex*diet interaction. The negative linear effect of protein on dILP8 expression is less negative in males.

Do we have the power to detect a female protein effect-size in males?

```r
anova(dILP8.m)
```

```
## Analysis of Variance Table
##
## Response: dILP8
##               Df Sum Sq Mean Sq F value Pr(>F)
```

```
## poly(prot, 2)  2  0.435 0.21729  0.1277 0.8805
## Residuals     46 78.300 1.70217
```

```r
summary(dILP8.m)
```

```
##
## Call:
## lm(formula = dILP8 ~ (poly(prot, 2)), data = m.data)
##
## Residuals:
##     Min      1Q  Median      3Q     Max
## -2.5949 -0.8071 -0.0448  0.4390  3.7340
##
## Coefficients:
##                 Estimate Std. Error t value Pr(>|t|)
## (Intercept)      -2.1932     0.1936 -11.328 6.66e-15 ***
## poly(prot, 2)1  -0.2329     2.1245  -0.110    0.913
## poly(prot, 2)2   0.7797     1.9085   0.409    0.685
## ---
## Signif. codes:  0 '***' 0.001 '**' 0.01 '*' 0.05 '.' 0.1 ' ' 1
##
## Residual standard error: 1.305 on 46 degrees of freedom
##   (26 observations deleted due to missingness)
## Multiple R-squared:  0.00552,    Adjusted R-squared:  -0.03772
## F-statistic: 0.1277 on 2 and 46 DF,  p-value: 0.8805
```

```r
pwr.f2.test(u = 1, v = summary(dILP8.m)$df[2], f2 = (summary(dILP8.f)$r.squared/(1 -
    summary(dILP8.f)$r.squared)), sig.level = 0.05)
```

```
##
##      Multiple regression power calculation
##
##               u = 1
##               v = 46
##              f2 = 0.4630899
##       sig.level = 0.05
##           power = 0.9960259
```

Plot the model and the TPS

```
## Warning:
## Grid searches over lambda (nugget and sill variances) with  minima at the endpoints:
##   (GCV) Generalized Cross-Validation
##    minimum at  right endpoint  lambda  =  164.2074 (eff. df= 3.000996 )
```

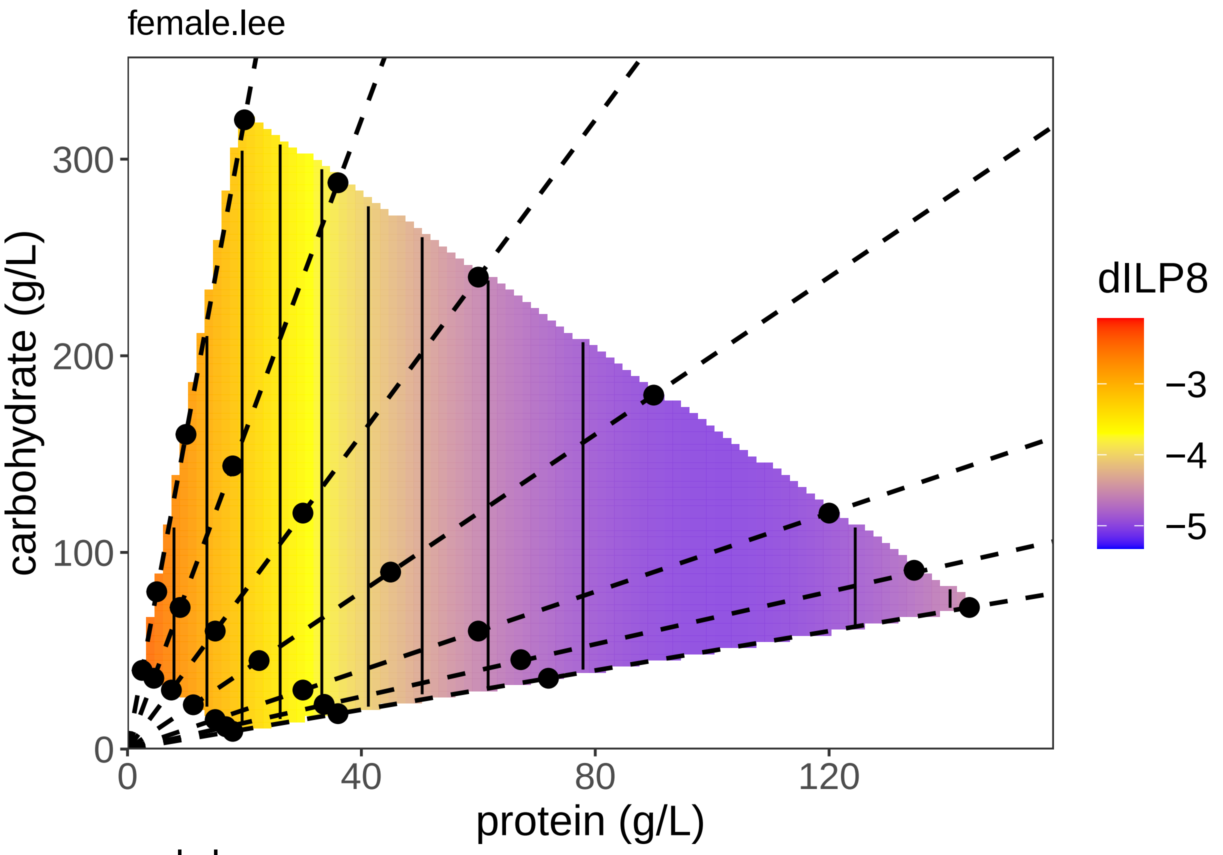
female.lee
dILP8
−3
−4
−5
carbohydrate (g/L)
300
200
100
0
0
40
80
120
protein (g/L)

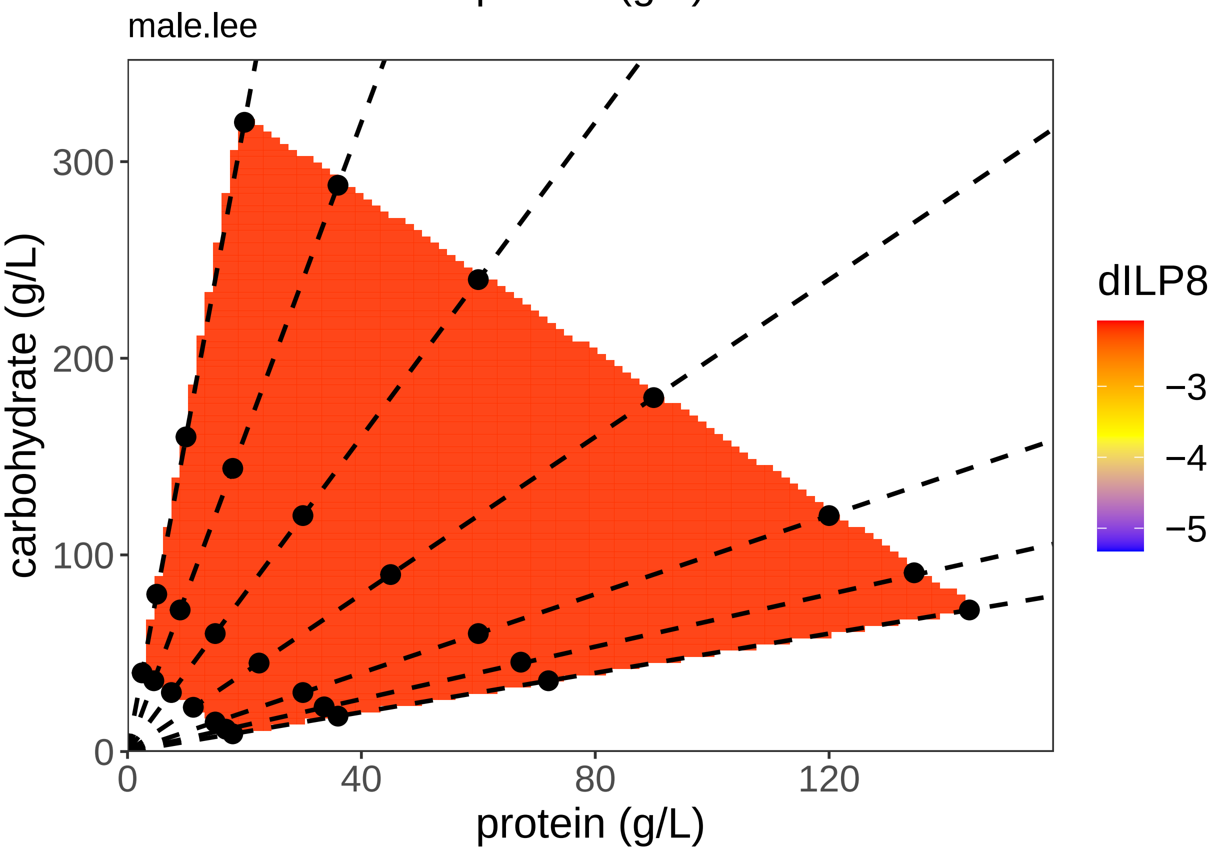
male.lee
dILP8
−3
−4
−5
carbohydrate (g/L)
300
200
100
0
0
40
80
120
protein (g/L)

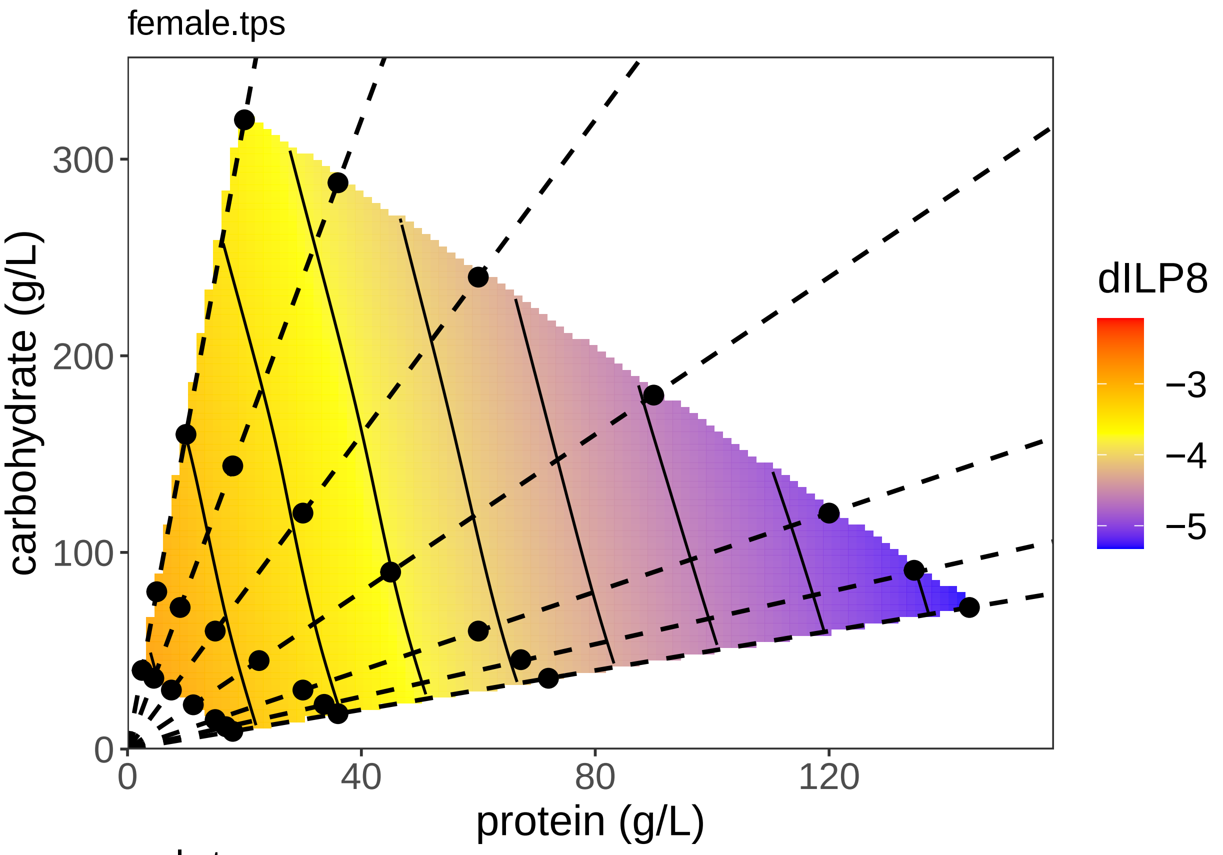
female.tps
carbohydrate (g/L)
protein (g/L)
dILP8
−3
−4
−5
0
100
200
300
0
40
80
120

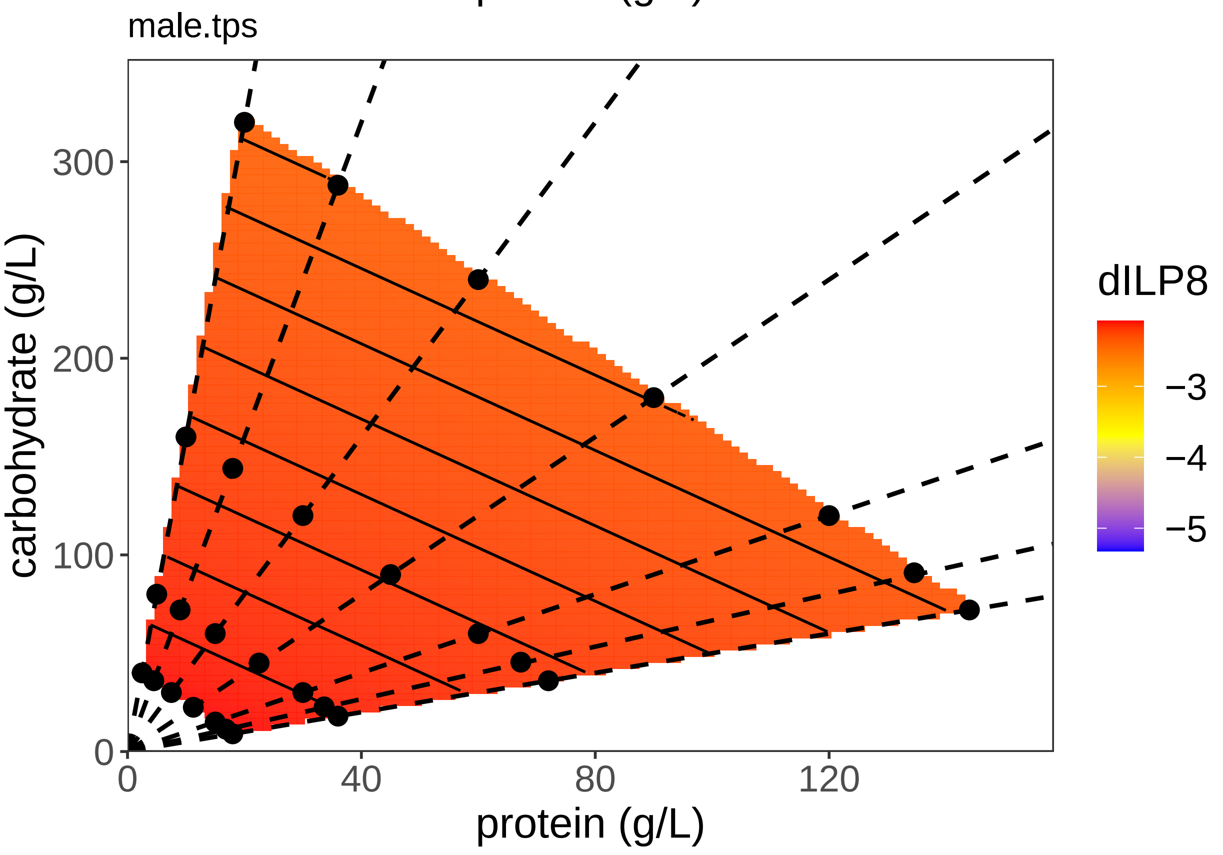
male.tps
carbohydrate (g/L)
protein (g/L)
dILP8
−3
−4
−5
0
100
200
300
0
40
80
120

#### Sample variablity

It is useful to see how much variablity there is in gene expression among samples within diets. We can inspect the EMS for each gene after fitting a one-way ANOVA of gene expression against

```
## Analysis of Variance Table
##
## Response: InR
##           Df Sum Sq Mean Sq F value    Pr(>F)
## Sample    29 54.974 1.89566  3.9012 4.853e-05 ***
## Residuals 39 18.951 0.48592
## ---
## Signif. codes:  0 '***' 0.001 '**' 0.01 '*' 0.05 '.' 0.1 ' ' 1
```

```
## Analysis of Variance Table
##
## Response: InR
##           Df Sum Sq Mean Sq F value    Pr(>F)
## Sample    27 25.274 0.93609  2.7502 0.0007759 ***
## Residuals 54 18.380 0.34037
## ---
## Signif. codes:  0 '***' 0.001 '**' 0.01 '*' 0.05 '.' 0.1 ' ' 1
```

```
##
##  F test to compare two variances
##
## data:  var.m and var.f
## F = 1.4276, num df = 39, denom df = 54, p-value = 0.2241
## alternative hypothesis: true ratio of variances is not equal to 1
## 95 percent confidence interval:
##  0.8026526 2.6121768
## sample estimates:
## ratio of variances
##           1.427616
```

```
## Analysis of Variance Table
##
## Response: X4eBP
##           Df Sum Sq Mean Sq F value   Pr(>F)
## Sample    30 18.512 0.61708  2.4286 0.003836 **
## Residuals 43 10.926 0.25409
## ---
## Signif. codes:  0 '***' 0.001 '**' 0.01 '*' 0.05 '.' 0.1 ' ' 1
```

```
## Analysis of Variance Table
##
## Response: X4eBP
##           Df  Sum Sq Mean Sq F value    Pr(>F)
## Sample    27 13.7807 0.51040  2.9154 0.0004071 ***
## Residuals 54  9.4536 0.17507
## ---
## Signif. codes:  0 '***' 0.001 '**' 0.01 '*' 0.05 '.' 0.1 ' ' 1
```

```
##
##  F test to compare two variances
##
## data:  var.m and var.f
```

```
## F = 1.4514, num df = 43, denom df = 54, p-value = 0.1941
## alternative hypothesis: true ratio of variances is not equal to 1
## 95 percent confidence interval:
##  0.8257575 2.5995086
## sample estimates:
## ratio of variances
##           1.451388

## Analysis of Variance Table
##
## Response: CG3071
##           Df  Sum Sq Mean Sq F value  Pr(>F)
## Sample    29 13.1052 0.45190  1.9006 0.04864 *
## Residuals 27  6.4197 0.23777
## ---
## Signif. codes:  0 '***' 0.001 '**' 0.01 '*' 0.05 '.' 0.1 ' ' 1

## Analysis of Variance Table
##
## Response: CG3071
##           Df Sum Sq Mean Sq F value    Pr(>F)
## Sample    26 43.087  1.6572  6.4482 2.403e-08 ***
## Residuals 46 11.822  0.2570
## ---
## Signif. codes:  0 '***' 0.001 '**' 0.01 '*' 0.05 '.' 0.1 ' ' 1

##
##  F test to compare two variances
##
## data:  var.m and var.f
## F = 0.92516, num df = 27, denom df = 46, p-value = 0.8457
## alternative hypothesis: true ratio of variances is not equal to 1
## 95 percent confidence interval:
##  0.4813848 1.8904364
## sample estimates:
## ratio of variances
##          0.9251612

## Analysis of Variance Table
##
## Response: Ash2L
##           Df Sum Sq Mean Sq F value Pr(>F)
## Sample    31 16.353 0.52751  1.1681 0.3145
## Residuals 43 19.418 0.45159

## Analysis of Variance Table
##
## Response: Ash2L
##           Df Sum Sq Mean Sq F value    Pr(>F)
## Sample    27 30.069 1.11367  4.0666 7.176e-06 ***
## Residuals 52 14.241 0.27385
## ---
## Signif. codes:  0 '***' 0.001 '**' 0.01 '*' 0.05 '.' 0.1 ' ' 1

##
##  F test to compare two variances
##
```

```
## data:  var.m and var.f
## F = 1.649, num df = 43, denom df = 52, p-value = 0.08531
## alternative hypothesis: true ratio of variances is not equal to 1
## 95 percent confidence interval:
##  0.9322235 2.9639623
## sample estimates:
## ratio of variances
##           1.649003
```

```
## Analysis of Variance Table
##
## Response: dILP2
##           Df Sum Sq Mean Sq F value Pr(>F)
## Sample    28 65.930  2.3546  1.0647 0.4414
## Residuals 24 53.079  2.2116
```

```
## Analysis of Variance Table
##
## Response: dILP2
##           Df Sum Sq Mean Sq F value    Pr(>F)
## Sample    25 33.984 1.35935  5.3362 3.514e-07 ***
## Residuals 48 12.228 0.25474
## ---
## Signif. codes:  0 '***' 0.001 '**' 0.01 '*' 0.05 '.' 0.1 ' ' 1
```

```
##
##  F test to compare two variances
##
## data:  var.m and var.f
## F = 8.6819, num df = 24, denom df = 48, p-value = 3.266e-10
## alternative hypothesis: true ratio of variances is not equal to 1
## 95 percent confidence interval:
##   4.466349 18.347965
## sample estimates:
## ratio of variances
##           8.681866
```

```
## Analysis of Variance Table
##
## Response: dILP3
##           Df Sum Sq Mean Sq F value Pr(>F)
## Sample    30 43.789  1.4596   0.935 0.5706
## Residuals 43 67.126  1.5611
```

```
## Analysis of Variance Table
##
## Response: dILP3
##           Df  Sum Sq Mean Sq F value    Pr(>F)
## Sample    27 20.8985 0.77402  4.8086 6.261e-07 ***
## Residuals 52  8.3702 0.16096
## ---
## Signif. codes:  0 '***' 0.001 '**' 0.01 '*' 0.05 '.' 0.1 ' ' 1
```

```
##
##  F test to compare two variances
##
## data:  var.m and var.f
```

```
## F = 9.6982, num df = 43, denom df = 52, p-value = 2.442e-13
## alternative hypothesis: true ratio of variances is not equal to 1
## 95 percent confidence interval:
##   5.482642 17.431812
## sample estimates:
## ratio of variances
##           9.698206

## Analysis of Variance Table
##
## Response: dILP5
##           Df Sum Sq Mean Sq F value Pr(>F)
## Sample    30 41.347 1.37824  1.4424 0.1367
## Residuals 41 39.176 0.95552

## Analysis of Variance Table
##
## Response: dILP5
##           Df Sum Sq Mean Sq F value    Pr(>F)
## Sample    27 42.556 1.57615  3.0338 0.0002938 ***
## Residuals 52 27.015 0.51953
## ---
## Signif. codes:  0 '***' 0.001 '**' 0.01 '*' 0.05 '.' 0.1 ' ' 1

##
##  F test to compare two variances
##
## data:  var.m and var.f
## F = 1.8392, num df = 41, denom df = 52, p-value = 0.03822
## alternative hypothesis: true ratio of variances is not equal to 1
## 95 percent confidence interval:
##  1.033923 3.339588
## sample estimates:
## ratio of variances
##           1.839221

## Analysis of Variance Table
##
## Response: dILP8
##           Df Sum Sq Mean Sq F value  Pr(>F)
## Sample    27 56.234  2.0827  1.9439 0.06102 .
## Residuals 21 22.500  1.0714
## ---
## Signif. codes:  0 '***' 0.001 '**' 0.01 '*' 0.05 '.' 0.1 ' ' 1

## Analysis of Variance Table
##
## Response: dILP8
##           Df Sum Sq Mean Sq F value  Pr(>F)
## Sample    27 80.795  2.9924  1.7261 0.04876 *
## Residuals 48 83.215  1.7336
## ---
## Signif. codes:  0 '***' 0.001 '**' 0.01 '*' 0.05 '.' 0.1 ' ' 1

##
##  F test to compare two variances
##
```

```
## data:  var.m and var.f
## F = 0.61803, num df = 21, denom df = 48, p-value = 0.2308
## alternative hypothesis: true ratio of variances is not equal to 1
## 95 percent confidence interval:
##  0.3108393 1.3686834
## sample estimates:
## ratio of variances
##            0.61803
```
